# Systematic benchmarking of mass spectrometry-based antibody sequencing reveals methodological biases

**DOI:** 10.1101/2024.11.11.622451

**Authors:** Maria Chernigovskaya, Khang Lê Quý, Maria Stensland, Sachin Singh, Rowan Nelson, Melih Yilmaz, Konstantinos Kalogeropoulos, Pavel Sinitcyn, Anand Patel, Natalie Castellana, Stefano Bonissone, Stian Foss, Jan Terje Andersen, Geir Kjetil Sandve, Timothy Patrick Jenkins, William S. Noble, Tuula A. Nyman, Igor Snapkow, Victor Greiff

## Abstract

The circulating antibody repertoire is crucial for immune protection, holding significant immunological and biotechnological value. While bottom-up mass spectrometry (MS) is the most widely used proteomics technique for profiling the sequence diversity of circulating antibodies (Ab-seq), it has not been thoroughly benchmarked. We quantified the replicability and robustness of Ab-seq using six monoclonal antibodies with known protein sequences in 70 different combinations of concentration and oligoclonality, both with and without polyclonal serum IgG background. Each combination underwent four protease treatments and was analyzed across four experimental and three technical replicates, totaling 3,360 LC-MS/MS runs. We quantified the dependence of MS-based Ab-seq identification on antibody sequence, concentration, protease, background signal diversity, and bioinformatics setups. Integrating the data from experimental replicates, proteases, and bioinformatics tools enhanced antibody identification. *De novo* peptide sequencing showed similar performance to database-dependent methods for higher antibody concentrations, but *de novo* antibody reconstruction remains challenging. Our work provides a foundational resource for the field of MS-based antibody profiling.

## Introduction

Immunoglobulins (Ig) provide the foundation for the humoral immune system by binding to a diverse and ever-changing array of threats known as antigens. There exist two forms of Ig: (1) transmembrane B-cell receptors (BCR) when displayed on the surface of B cells and (2) antibodies (Ab) when secreted in a soluble truncated form into the serum and mucosa. BCRs and antibodies possess mostly identical structures of two heavy chains (HC or IGH) and two light chains (LC), where the LCs can be either of the kappa (IGK) or lambda (IGL) type ^1^. Each chain consists of a constant region and a variable region. While the constant region determines effector functions, the variable region determines specificity to antigens and is composed of V, (D), and J genes assembled in a process called V(D)J recombination. The variable region is subdivided into the lower diversity framework regions and the higher diversity complementary determining regions, known as CDR1, CDR2, and CDR3 ^2^. Most of the diversity in Ig sequences is concentrated in the CDR3, which is often utilized as a unique identifier of clonal lineage amongst Ig molecules ^3–5^ and has been shown to be crucial for antigen binding ^6,7^. In this manuscript, CDR3 regions of HC and LC are referred to as CDRH3 and CDRL3, respectively. During the response to antigens, the affinity maturation process introduces point mutations that further enhance Ig sequence diversity ^8,9^. The stochastic and combinatorial nature of these processes leads to a theoretical diversity of over 10^13^ human Ig sequences ^3,10^.

Over the last decade, high-throughput BCR sequencing (BCR-seq) has provided unprecedented insights into the genetic diversity of the B-cell immune response ^11–17^. BCR-seq enabled the study of B-cell repertoires using sequence-based computational methods, such as repertoire similarity, diversity, clonality, and architecture analysis ^16^. In contrast to BCRs, the sequence diversity and antigen-specific dynamics of the mucosal and serological antibody repertoire have remained underexplored ^18–23^. The lack of a detailed description of the sequence identities and relative amounts of circulating antibodies in the repertoire prevents the establishment of the fundamental principles of humoral adaptive immunity. This, in turn, has hindered the development of next-generation therapeutics, vaccines, and diagnostics, as well as the profiling of antibody sequence diversity in infection, cancer, and autoimmunity ^18,19,24–44^.

Since antibodies are soluble proteins that circulate freely in body fluids, they cannot be sequenced directly as opposed to BCRs. To analyze the sequence diversity of polyclonal antibody mixtures, mass spectrometry (MS) is required – hereafter referred to as Ab-seq ^18,21^. Specifically, Ab-seq signifies the use of bottom-up proteomics analysis with LC-MS/MS to recover amino acid sequences from digested Ab molecules, in contrast to intact mass analysis of native Ab molecules in top-down MS ^25,26^. To enable sequence recovery, BCR-seq data is usually required to build the reference database, from which an *in silico* spectrum as a basis to evaluate the experimental spectra, formally referred to as peptide-spectrum match (PSM) (Figure 1a). Alternatively, *de novo* peptide sequencing tools may be used ^45–47^. Bottom-up proteomics is by far the most common approach to capturing the amino acid sequence diversity ^18^, however, recently, other approaches combining both bottom-up and top-down approaches have been introduced ^32^.

**Figure 1:**
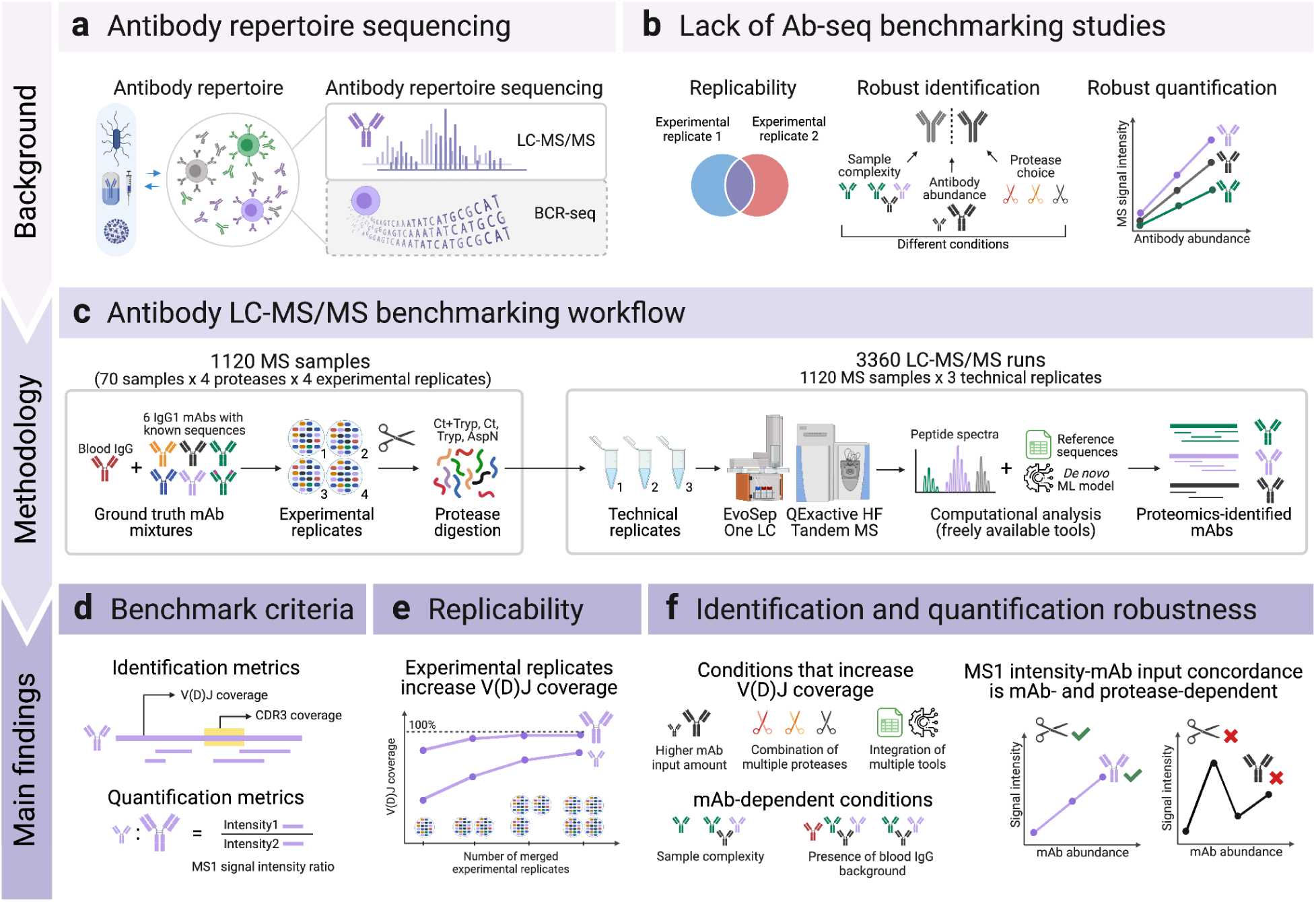
An integrated experimental and computational approach to Ab-seq benchmarking. (a) Antibody repertoire sequencing (Ab-seq) usually involves tandem analysis of LC-MS/MS and BCR-seq data to enable the proteomic characterization of antibody repertoire diversity, unless de novo peptide sequencing is employed. (b) There is a lack of benchmarking studies investigating Ab-seq replicability and robustness of identification and quantification, hindering the reliability of Ab-seq data. (c) To address this knowledge gap, we utilized six IgG1 mAb spike-ins to systematically benchmark antibody bottom-up proteomics. Our study included 1120 MS samples (70 samples × 4 experimental replicates × 4 protease treatments) and 3360 LC-MS/MS runs (1120 MS samples × 3 technical replicates). The MS samples varied in terms of antibody input amount, antibody sequence similarity, protease treatments, and the presence of blood IgG background. The raw data were analyzed using peptide-spectrum match (PSM) and de novo proteomics computational platforms that are freely available. (d) We used V(D)J and CDR3 coverage as metrics of identification quality and MS1 signal intensity ratio as a quantification estimate. (e) We found that combining multiple experimental replicates improves the replicability of mAb sequence detection and increases sequence coverage. (f) To study Ab-seq robustness, we investigated how different conditions influence the coverage of antibody sequences as well as the extent to which MS1 signal intensity may be used to infer relative antibody abundance.

Ab-seq is a special case of the protein inference problem in bottom-up proteomics, where proteins in a sample are identified based on the detected peptides. While protein inference has been extensively benchmarked in general cases ^48^ and serves as the basis for protein quantification ^49^, Ab-seq poses unique challenges. Unlike standard protein inference, V(D)J recombination and somatic hypermutation produce a highly degenerate protein database, where two antibodies in the repertoire may differ by only a single mutation. In general protein inference, detecting one or two peptides is often sufficient to confirm the presence of a protein. However, in Ab-seq, the high frequency of shared peptides between similar antibodies means that accurate identification requires the detection of multiple peptides, particularly those covering the CDR3s.

While many reports exist on B- and T-cell receptor sequencing standardization ^50–55^, benchmarking studies for Ab-seq are virtually absent, despite numerous reports on this subject ^18,21,29,32,36,56^. Benchmarking of high-throughput technologies depends on ground truth data ^57^, where such data consists of, for example, known cell lines, or known nucleic or protein sequences (i.e., “spike-in” sequences), allowing for estimation of identification accuracy. Indeed, a recent large-scale shotgun proteomics study performed on cell lines established baselines for the correspondence of transcriptome and translated proteins ^58^. While some previous Ab-seq studies utilized spike-in, their main purpose was relative quantification of sample input rather than benchmarking performance ^32,36,39,59,60^. Thus, as of present, the extent to which bottom-up Ab-seq can accurately recover antibody sequences across a wide range of experimental conditions varying by background, antibody concentration, and complexity of the polyclonal mixture has not been investigated systematically (Figure 1b). Additionally, the replicability and robustness of Ab-seq remain largely unknown (Figure 1b). Quantifying the impact of these factors could facilitate the joint analysis of BCR sequencing and Ab-seq ^61^, with the ultimate goal of achieving accurate and comprehensive characterization of serum antibody diversity.

In this study, we devised a large-scale, sequence- and input amount-controlled experimental design with multiple independent repetitions of the entire experiment (experimental replicates) and multiple sample injections into the mass spectrometer (technical replicates) (Figure 1c). This experimental design was used to assess the performance of LC-MS/MS for Ab-seq sequence reconstruction with respect to (1) experimental and technical replicability, (2) protease-dependent variation, (3) presence of blood-derived Ig background, (4) antibody input amount, (5) antibody sequence similarity, (6) freely available protein identification and quantification tools, and (7) *de novo* peptide sequencing from MS data. We show that (i) performing experimental replicates improves peptide discovery and sequence coverage (Figure 1e), (ii) detection of monoclonal antibodies is affected by sample polyclonal complexity and presence of blood IgG background (Figure 1f), (iii) antibody concentration differences may be inferred using Ab-seq’s MS1 signal intensities (Figure 1f), and (iv) *de novo* Ab-seq remains inferior in performance compared to PSM methods in V(D)J sequence coverage (Figure 1f). Overall, our Ab-seq dataset, obtained from 3360 MS runs, provides a foundation for future refinement of both experimental and computational Ab-seq workflows.

## Results

### Experimental design for Ab-seq benchmarking

To benchmark the Ab-seq method, we designed an experiment incorporating a diverse source of variations. As ground truth, we used six monoclonal IgG1 antibodies (mAbs) with known amino acid sequence: h9C12 WT (wild type), h9C12 Q97A (mutant), PGT121, and PGDM1400 as well as recombinant forms of Briakinumab (short: Brimab), Ustekinumab (short: Umab), (Supplementary Table 1, Supplementary Table 2). The mAbs are well-characterized and have known specificities: h9C12 and its variant h9C12 Q97A bind the hexon surface protein of human adenovirus 5 ^62^, Brimab and Umab bind human IL12/23 ^63,64^ while PGT121 and PGDM1400 target the HIV gp120 envelope protein ^65,66^. Except for h9C12 WT and h9C12 Q97A, which differed by only one amino acid residue in the CDRH3 region, all tested mAbs exhibited low pairwise sequence similarity of 0% to 44.4% within the CDRH3 region (Supplementary Table 3), where CDR3 was defined by the IMGT numbering scheme for the V-REGION ^67^. In addition, we verified that none of the six mAbs except h9C12 WT and h9C12 Q97A share common subsequences within CDRH3 regions more than 5 amino acids long (Supplementary Table 4). The intentional use of diverse antibodies in the experiment was to minimize collisions in peptide detection and quantification among different mAbs.

In the main experiment, we varied the mAb input amount across three orders of magnitude (from 10 ng/mL to 10 μg/mL in 100 μL volume), the number of mAbs included (from 1 to 6 mAbs), and the presence of background noise (polyclonal serum IgGs), resulting in a total of 70 unique mAbs combinations, hereafter referred to as samples. We repeated the experiment, starting from digestion to LC-MS/MS analysis, for all 70 samples across four experimental replicates, ensuring a high degree of experimental consistency (Supplementary Table 5). As suggested by findings by Bondt and colleagues ^32,68^, the majority of IgG1 and IgA1 repertoires consist of tens to hundreds of the most abundant clones. To approximate clonal complexity, we used six mAbs and varied the clonal composition by adjusting the number of mAbs in each sample. The range of mAb input (from 1 ng to 1 μg) was chosen to reflect the physiological range at which IgG antibody clonotypes would be present in the serum, based on previously measured serum IgG concentration ^69^ and the estimated number of clonotypes present in the serum ^70^. Briefly, single Ab clonotypes have been reported to be within the range of 10 ng to 1 μg per 1 mL of human plasma ^32^, while 1 ng of input (equivalent to 6.7 fmol of antibodies) is used to examine the detection limit of bottom-up proteomics for Ab clonotype identification. Polyclonal IgGs isolated from the blood of a healthy donor were added to half of the samples (1 μg in samples 1-35, 65, 67, and 68; 50 μg in samples 69 and 70) to simulate the effect of polyclonal IgG background noise (Figure 2a, Supplementary Table 6).

**Figure 2:**
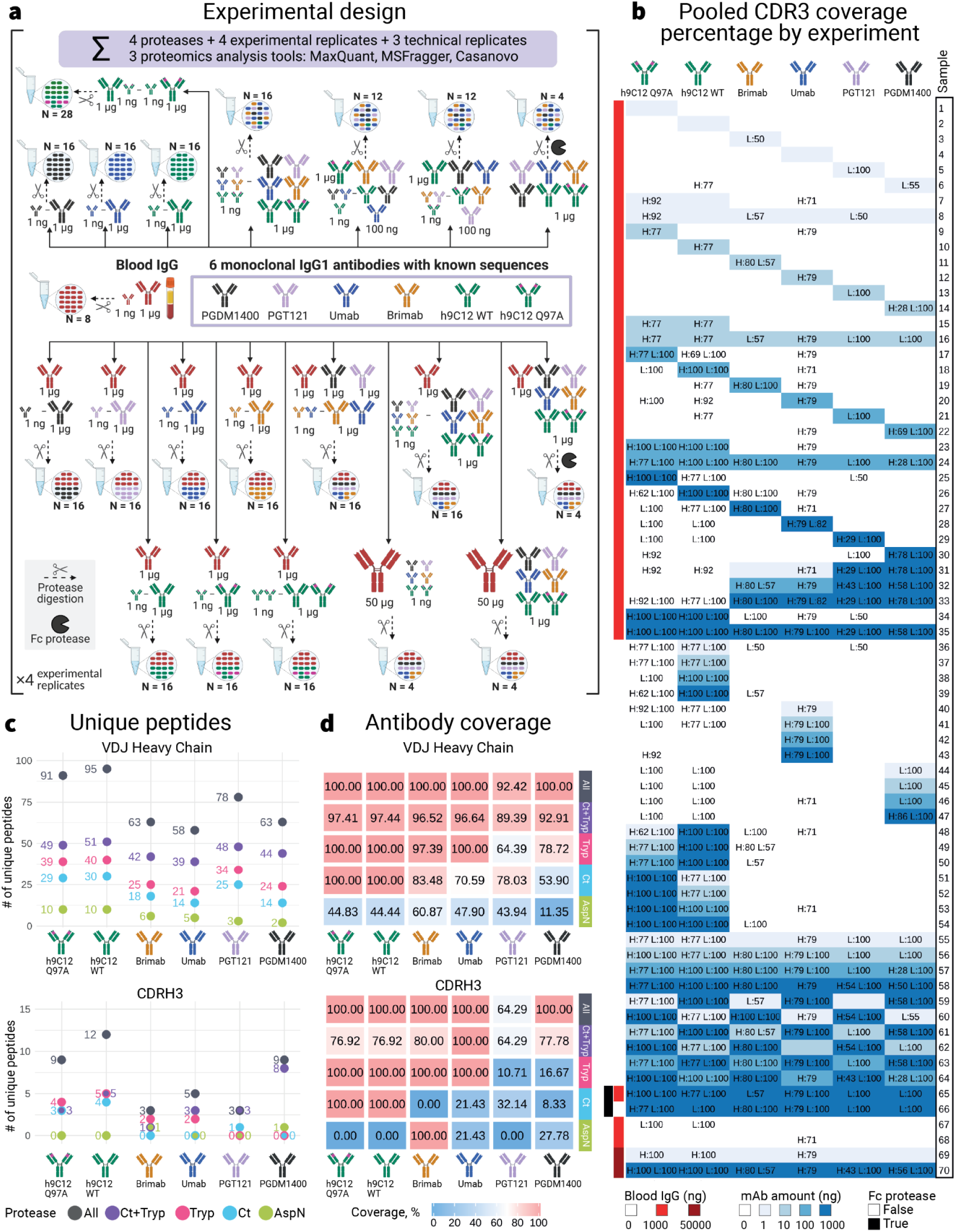
The choice of protease and antibody sequence strongly affects Ab-seq variable region sequence coverage. **(a)** The benchmarking dataset consists of 70 unique antibody combinations (samples) with varying complexity, including the number of mAb spike-ins (1–6 mAbs from the list of six mAbs of known sequences — h9C12 Q97A, h9C12 WT, Brimab, Umab, PGT121, and PGDM1400) and their input amount (1–1000 ng), along with the optional addition of peripheral blood IgG polyclonal antibodies (0 ng, 1000 ng, or 50000 ng). Each sample was treated with four digestion strategies — Ct, Tryp, Ct+Tryp, AspN, and samples 65 and 66 were additionally treated with the Fc cleavage protease. The whole experiment setup was replicated four times (experimental replicates). The output of each LC-MS/MS run was analyzed using three proteomics analysis tools — MaxQuant (MQ), MSFragger (MSF), and Casanovo. **(b)** The number of unique CDR3 peptides, i.e., peptides with at least a three amino acid overlap with the CDR3 region of the mAb according to the IMGT numbering scheme for V-REGION ^67^ was identified in every sample and for every mAb. Peptides were pooled from all experimental and technical replicates, all proteases, and all computational tools (MQ, MSF, Casanovo). The heatmap cell is colored based on the mAb input (ranging from 1 ng to 1000 ng), while a white background indicates the absence of the mAb in the sample. H:xx L:yy in the heatmap cell represents the number of peptides in the HC and the LC, correspondingly. The number of peptides displayed in the blue fields are considered true positives, while the numbers displayed in the white fields are considered false positives. (**c)** The number of unique true positive peptides in the full-length VDJ region and overlapping with the CDRH3 region of antibody HCs. Peptide counts are pooled for all experimental, all technical replicates, and all proteomics analysis tools. **(d)** Full-length HC VDJ and CDRH3 coverage (the percentage of the reference mAb sequence detected by Ab-seq) of true positive peptides pooled from all experimental replicates, all technical replicates, and all proteomics analysis tools achieved by every protease digestion. See also **Supplementary Figure 13** for peptide count and sequence coverage for LC peptides.

Each sample was digested separately by four commonly used protease treatments ^71,72^: Trypsin (Tryp), Chymotrypsin alone (Ct), Chymotrypsin followed by Trypsin (Ct+Tryp), or AspN, resulting in a total of 1120 MS samples (70 samples x 4 experimental replicates x 4 protease treatments) (Figure 1c). The rationale was that aside from Trypsin, the most common choice for protease treatment ^73^, Chymotrypsin and AspN can be used to complement Trypsin to achieve a better distribution of cleavage sites peptides in the target sequence ^74^. In addition to the four proteases, samples 65 and 66 were treated with Fc protease followed by purification of the derived Fab fragments to evaluate whether Fab-only characterization would result in better performance compared to intact mAbs. However, we did not observe that Fc protease increased the detection of mAb-related peptides from the samples (Supplementary Figure 1). All digested peptides were fractionated with the EvoSep One LC system, followed by detection in QExactive HF mass spectrometer, with each sample analyzed in technical triplicates, resulting in a total of 3360 LC-MS/MS runs (1120 MS samples x 3 technical replicates). The raw output files were searched against a reference database containing the ground truth mAb sequences, along with decoy protein and antibody sequences (details in Methods) using MaxQuant (MQ) ^75^ and MSFragger (MSF) ^76^, as well as *de novo* peptide sequencing tool Casanovo ^45^, all of which are freely available software packages. Although other computational tools for proteomics analysis exist ^77^, our study focused solely on freely available software.

To evaluate Ab-seq performance, we examined both its replicability and robustness in identification and quantification (Figure 1b). Here, replicability is defined as the consistency of Ab-seq results across experimental and technical replicates (Figure 1e), while robustness refers to the impact of various conditions on Ab-seq performance (with benchmarking metrics illustrated in Figure 1d). These conditions include experimental factors, (such as background noise, mAb input quantity, and mAb mixture complexity) and computational factors (such as the choice between PSM or *de novo* approaches and the selection of computational tools) (Figure 1f). The low variation in signal intensities and peptide detection between technical replicates, chromatography method, and consumables (Supplementary Figure 4, Supplementary Figure 2, Supplementary Figure 3) allowed us to better discern the variability across experimental replicates and conditions. Consequently, unless otherwise specified, the three technical replicates for each sample are pooled in all reported results in the main or supplementary figures.

Overall, the choice of sample content, preparation methods, experimental and computational conditions, along with the selected performance metrics, allowed us to conduct an in-depth analysis of Ab-seq as a tool to characterize the circulating antibody repertoire sequence diversity.

### False peptide identifications may result from choice of software and antibody sequence similarity

In our benchmarking dataset, we had ground truth knowledge of the mAbs present in each sample, enabling us to detect incorrectly identified peptides. We defined false positive matches as peptides that mapped to the V(D)J region of a mAb not present in the sample. Notably, these false positive peptides systematically appeared across experimental replicates (Supplementary Figure 5, Supplementary Figure 6) (experimental replicate 1: 18.98%, experimental replicate 2: 20.64%, experimental replicate 3: 22.58%, experimental replicate 4: 15.5%). Additionally, the false positive matches were consistent across computational tools used (Supplementary Figure 7), with MSFragger producing the highest number of false positive peptides overall (23.06%), compared to MaxQuant (13.32%) and Casanovo (16.26%). Brimab and Umab were the mAbs most frequently misidentified by all tools in the CDR3 (Supplementary Figure 7). This excludes the false positives between h9C12 WT and h9C12 Q97A, due to their identical LC and 99.14% HC similarity (Supplementary Table 3).

In general, samples containing blood IgGs exhibited a higher percentage of falsely identified peptides (20.68%) compared to samples containing mAbs-only (4.62%). However, the frequency of false positive matches was lower among CDR3 peptides, both in blood IgG-containing (1.08%) and mAb-only samples (0.64%) (Supplementary Figure 8). We hypothesized that the presence of these false positive peptides can be attributed to several factors: (i) the presence of polyclonal IgGs in blood IgG-containing samples, which may share common subsequences with the mAbs, particularly in the V or J gene regions, (ii) shared subsequences among different mAbs, and (iii) incomplete removal of previous peptides during the cleaning cycles.

To address these issues, we implemented a preprocessing pipeline, with the procedure detailed in the Methods section and Supplementary Figure 9. Briefly, (i) we subtracted false positive peptides aligning to the IMGT V and J gene reference database from the samples containing blood IgGs, (ii) reassigned false positive peptides shared between mAbs to the correct mAb, and (iii) subtracted false positive peptides if they were detected in the last step of the previous cleaning cycle. One cleaning cycle was performed after every three MS runs, and three cleaning cycles were performed after samples containing the highest input amount of mAbs (Supplementary Table 6).

The majority of false positive peptides removed during filtering (Supplementary Figure 10) were Ab-related peptides corresponding to the IMGT V(D)J database (75.26%). A smaller proportion of false positives were either shared between mAbs or identified during intermediate cleaning cycles (0.17% and 2.05% correspondingly). However, despite the implementation of this filtering procedure, a residual number of false positives persisted (22.52%), indicating potential unidentified sources of contamination (Supplementary Figure 11). To minimize bias from these remaining false positives, subsequent analyses focused exclusively on true positive matches — peptides that aligned with the mAbs expected to be present in each sample. The only exception was Figure 2b, which includes CDR3-related peptides that passed the filtering process.

### Coverage of antibody variable region is influenced by the protease and antibody sequence

We first evaluated the effects of key experimental conditions, such as protease choice and mAb sequence, on the Ab-seq identification (Figure 2). Although the number of unique peptides mapped to a mAb sequence can provide insights into Ab-seq performance, applications such as mAb cloning and expression of mAb sequence of interest require full V(D)J sequence characterization. This highlights the need for a more thorough assessment of mAb sequencing performance. For this reason, we prioritized sequence coverage, defined as the percentage of covered positions in the corresponding reference sequence, as the primary performance metric over the number of unique peptides. This choice reflects the fact that peptides of varying lengths can map to the same positions in the reference V(D)J sequence, making sequence coverage a more comprehensive measure of Ab-seq performance (see Supplementary Figure 12 for a detailed explanation of coverage). From this point on, sequence coverage will be the sole performance metric used in our analysis.

Figure 2 provides an overview of the primary features measured in the study, with the sample contents detailed in Figure 2a, and results pooled from all samples, replicates, proteases, and proteomics software tools. These include the CDRH3 and CDRL3 coverage per sample for all protease treatments (Figure 2b), the number of unique peptides aligning to the VDJ and CDRH3 sequences (Figure 2c), and the VDJ and CDRH3 sequence coverage per protease treatment in all samples combined (Figure 2d). For each mAb, the CDR3 coverage increased with higher input amounts (Figure 2b). The detection limit of mAb-specific CDR3 peptides, indicated by non-zero CDR3 coverage, was 10 ng in the presence of background blood IgGs and 1 ng without background blood IgGs on both HCs and LCs (Figure 2b). In addition, CDRH3 coverage was similar or better in samples without blood IgGs when comparing samples with equal input amounts (Figure 2b, Figure 4c).

In terms of the number of unique peptides obtained from each protease treatment, digestion by Ct+Tryp consistently yielded the largest number of unique VDJ peptides per mAb (39–51 unique peptides), followed by Tryp (21–40 peptides) and Ct (14–30 peptides). ApsN digestion, however, consistently resulted in the fewest number of unique mAb peptides (2–10 peptides) (Figure 2c). A similar order in protease efficiency was observed for CDRH3 peptides, as well as LC VJ and CDRL3 peptides (Supplementary Figure 13a). Notably, CDRH3 peptides were absent in certain protease treatments from all experimental replicates: h9C12 Q97A, h9C12 WT, Umab, and PGT121 for AspN; Brimab, Umab, and PGDM1400 for Ct; PGT121 and PGDM1400 for Tryp (Figure 2c), despite the theoretical cleavage predictions indicating the presence of CDRH3 peptides from all four proteases (Supplementary Figure 14). Additional details, including the peptide length distribution and miscleavage rates, are provided in Supplementary Figure 15.

By combining the sequence coverage from all proteases, we achieved over 90% VDJ region coverage for all six mAbs and 100% CDRH3 region coverage for five out of six mAbs, except PGT121 having 64.29% CDRH3 coverage (Figure 2d). Of note, theoretical coverage as determined by *in silico* cleavage was expected to be 100% (Supplementary Figure 14, Supplementary Figure 16, Supplementary Figure 17). Each protease treatment contributed differently to the overall sequence coverage, depending on the specific mAb. Ct+Tryp protease provided the highest overall VDJ sequence coverage across all six mAbs (89.39–97.44%), followed by Tryp peptides (64.39–100%), Ct peptides (53.90–100%), and AspN peptides (11.35–60.87%) (Figure 2d). Of note, the two mAbs PGT121 and PGDM1400, which had lower CDRH3 coverage with Tryp and Ct, also had the longest CDRH3 sequences out of the six mAbs (Supplementary Table 1). For the LC, full coverage was achieved for all mAbs when combining all proteases (Supplementary Figure 13b). As estimated by *in silico* digestion, AspN consistently delivered the lowest sequence coverage among the four proteases, producing longer peptides that are typically less amenable to mass spectrometry (Supplementary Figure 14). However, including AspN improved overall sequence coverage for certain mAbs, such as Brimab and PGDM1400, where Ct or Tryp alone were insufficient (Figure 2d, Supplementary Figure 14).

In summary, we show that both protease choice and antibody sequence have a major impact on the coverage of the variable region in Ab-seq. Among the tested conditions, Ct+Tryp digestion produced the highest number of unique peptides and the most comprehensive sequence coverage.

### Experimental replicates in Ab-seq improve sensitivity at low mAb input

To evaluate the potential additive effects of utilizing multiple experimental replicates, we cumulatively merged experimental replicates from one to four (Figure 3a), while pooling together peptides from three technical replicates, four protease treatments, and all computational tools. For each sample, V(D)J and CDR3 coverage were calculated for each mAb, grouping the results by mAb input amount (6 mAbs x 4 input amounts = 24 mAb-input groups). Our primary focus was on the HC sequence coverage (Figure 3b, c), similar trends were observed for the LC sequence coverage (Supplementary Figure 13).

**Figure 3:**
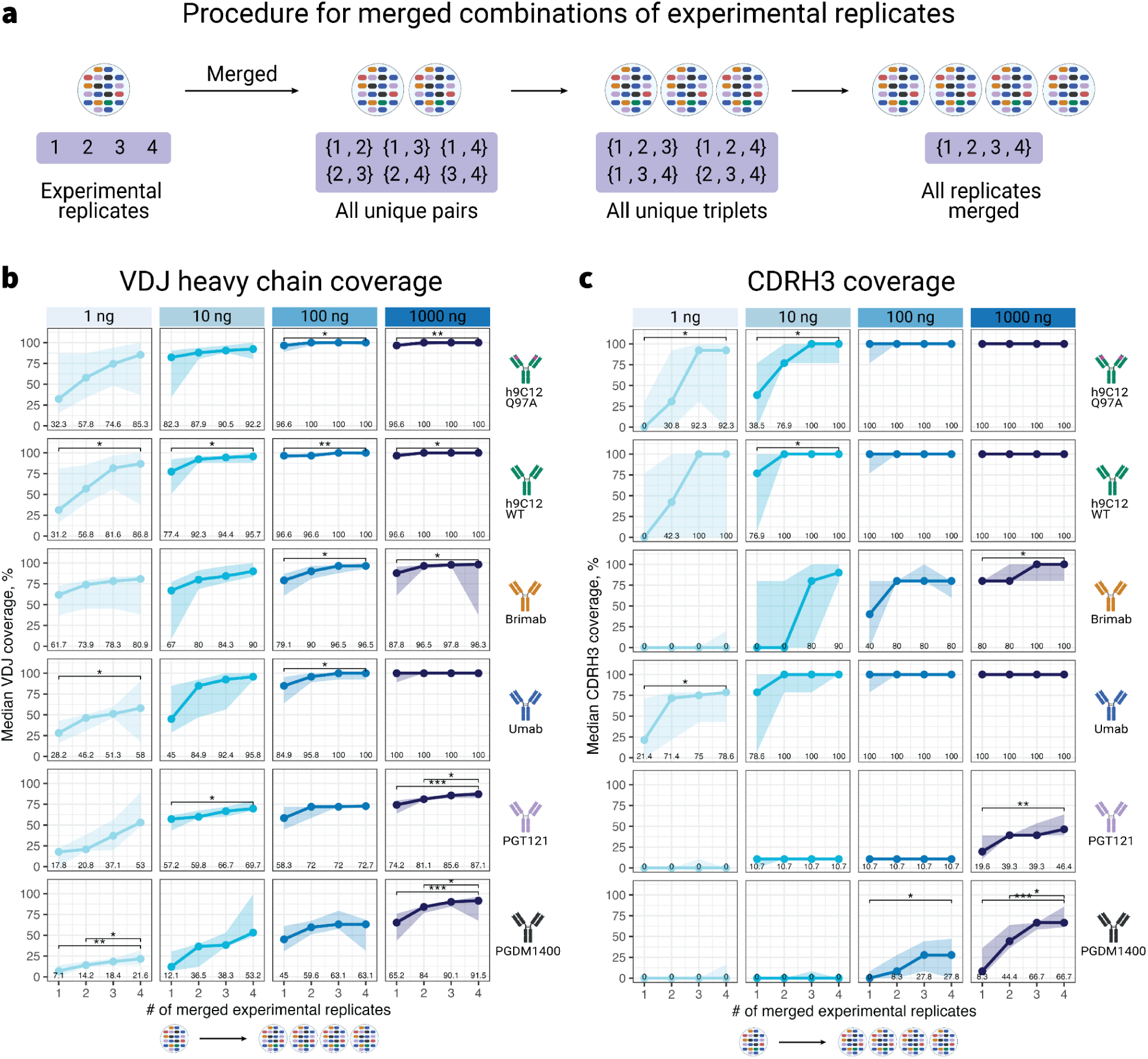
Experimental replicates are required for comprehensive V(D)J and CDR3 coverage. **(a)** Four experimental replicates were merged in all possible combinations: 1 (individual replicates), 2 (all unique pairs of replicates merged), 3 (all unique triplets of replicates merged), and 4 (all experimental replicates merged). True positive preprocessed peptides from all proteases, technical triplicates, and proteomics analysis software programs were pooled and separated by mAb and input amounts (ranging from 1 ng to 1000 ng), resulting in 24 mAb-input groups (6 mAbs x 4 input amounts). For each mAb-input group, we calculated the median **(b)** VDJ coverage and **(c)** CDRH3 coverage with respect to the number of merged experimental replicates. Median coverage for each number of merged replicates is shown below within each mAb-input group. The difference between median coverage values within each mAb-input group was tested using the Kruskal–Wallis test and adjusted for multiple testing across all 24 mAb-input groups using the Benjamini–Hochberg method. For mAb-input groups showing significant differences with the adjusted Kruskal–Wallis test results, we compared pairwise differences between individual experimental replicates, pairs, triplets, with all replicates merged (3 comparisons in total for each mAb-input group) using the Wilcoxon Rank Sum test, with p-values adjusted for multiple testing by Benjamini–Hochberg correction within each mAb-input group. All adjusted p-values lower than 0.05 are displayed above brackets as symbols: *(p.adj < 0.05), **(p.adj < 0.01), ***(p.adj < 0.001). Analogous stratification and analysis of samples on the LC are presented in **Supplementary Figure 19**. Coverage of the V(D)J and CDR3 regions with similar groupings for each experimental replicate individually and for all experimental replicates pooled together (∑) is shown in **Supplementary Figure 18**. The number of unique peptide sequences for each experimental replicates and their pairwise overlap of peptide sets is shown in **Supplementary Figure 20**.

By merging experimental replicates cumulatively and comparing pairwise differences between individual experimental replicates, pairs, triplets, with all replicates merged (3 comparisons in total for each mAb-input group) (Figure 3a), we observed a marked increase in sequence coverage, especially at lower mAb input amounts. At the lowest input of 1 ng, accumulating peptides from an increasing number of experimental replicates led to an increase in median VDJ coverage of 14.54–55.56 %pt, depending on the mAb sequence. The impact of merging replicates diminished at higher input amounts, with the gap in coverage reduced to 9.91–50.84%pt at 10 ng, 3.42–18.08%pt at 100 ng, and 0–26.24%pt at 1000 ng (Figure 3b). Each added replicate improved the median VDJ sequence coverage by 4.84%pt to 18.52%pt at 1 ng, 3.30%pt to 16.95%pt at 10 ng, 1.14%pt to 6.03%pt at 100 ng, while at 1000 ng the increase fluctuated between 0%pt and 8.75%pt.

Next, we evaluated whether the increase in VDJ sequence coverage from merging experimental replicates was statistically significant (adjusted p-value < 0.05). In 14 out of 24 mAb-input groups, VDJ coverage was significantly higher in four merged replicates than individual replicates (Figure 3b). However, when comparing two merged experimental replicates to four, only 3 out of 24 mAb-input groups showed a statistically significant increase. Finally, no significant difference in VDJ coverage was observed between three merged experimental replicates and four merged experimental replicates.

Similar to the VDJ region, CDRH3 coverage improved when experimental replicates were merged. In 8 out of 24 mAb-input groups, merging four experimental replicates significantly increased CDRH3 coverage compared to individual experimental replicates. However, only 1 out of 24 mAb-input groups showed a significant difference in CDRH3 coverage when comparing two versus four merged experimental replicates, largely due to the analytical challenge of samples exhibiting either zero or full CDRH3 coverage. Once again, PGT121 and PGDM1400, which had lower CDRH3 coverage, coincided with their longer CDRH3 length (Figure 3c, Supplementary Table 1).

The highest median VDJ coverages were reported more often in experimental replicates 2 (12/24 mAb-input groups) and 3 (9/24 mAb-input groups), compared to replicates 1 (5/24 mAb-input groups) and 4 (0/24 mAb-input groups). However, no single experimental replicate was universally superior in sequence coverage for all six mAbs surveyed (Supplementary Figure 18). Nevertheless, aggregating sequence data from all experimental replications invariably resulted in greater coverage than any single experimental instance, except where coverage had attained completeness at 100% (Figure 3b,c). Similar results were observed for the LC sequence coverage (Supplementary Figure 19).

To summarize, consolidating peptide sequence data from multiple experimental replicates improves V(D)J and CDR3 coverage, especially at low input amounts. Our results highlight the importance of using experimental replicates to enhance the robustness and comprehensiveness of serum antibody proteomic analyses.

### Presence of blood polyclonal IgG background may impact mAb CDRH3 coverage

Since Ab-seq is typically performed on polyclonal antibody mixtures, we investigated whether the presence of blood-derived polyclonal IgG could interfere with the identification of mAb-specific peptides. To explore this, we divided the samples into two groups (Figure 4a): mAb-only samples (samples 36–64, 66) and those containing blood-derived IgG background (samples 1–35, 65, 67–70). We pooled together peptides from technical triplicates, protease treatments, and proteomics software programs. As in Figure 3, we quantified VDJ and CDRH3 coverage across 24 different mAb-input groups (6 mAbs x 4 input amounts).

**Figure 4:**
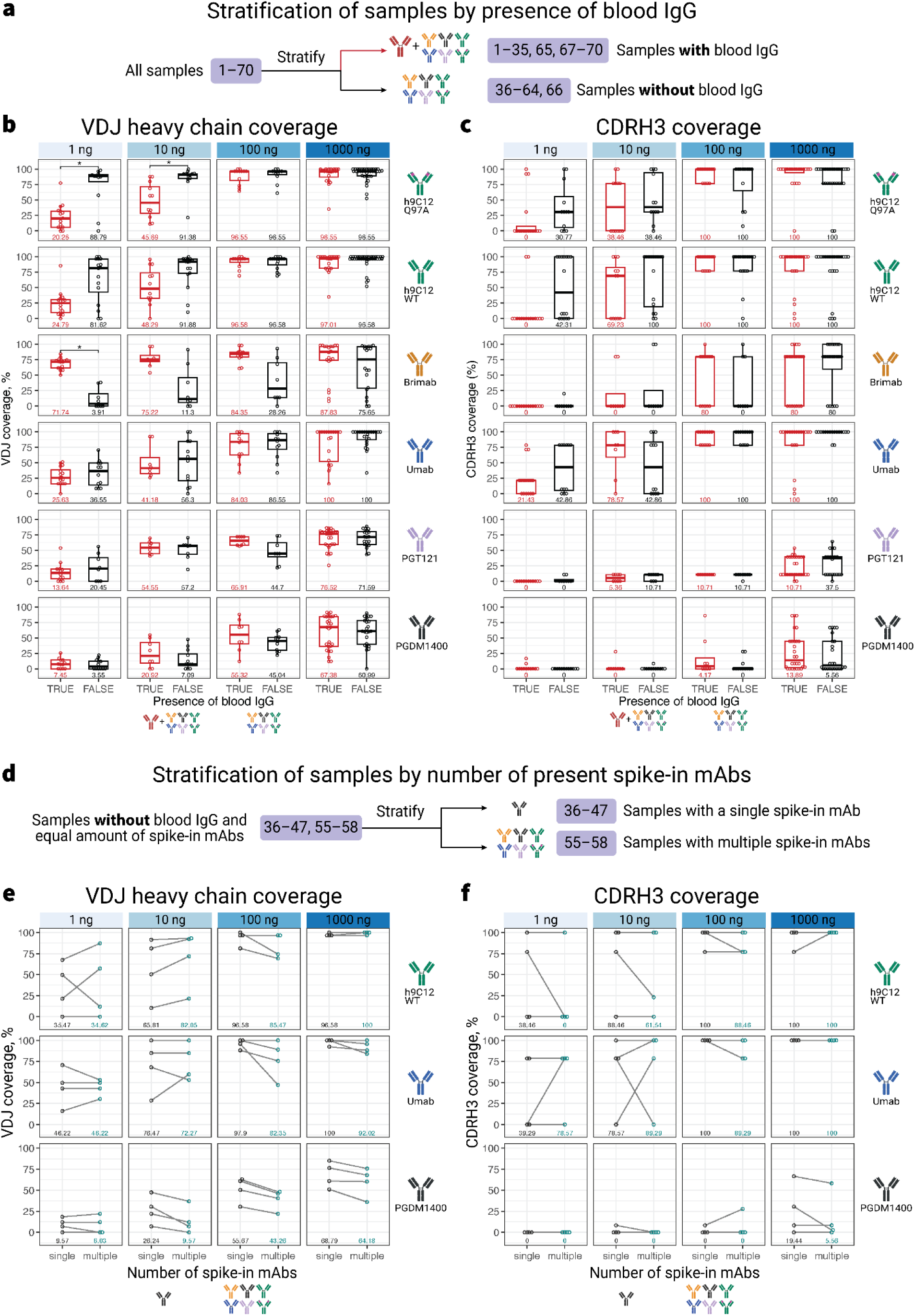
Diversity of antibodies in a sample influences sequence coverage in Ab-seq analysis. **(a)** We stratified 70 samples based on the presence (sample 1–35, 65, 67–70) or absence (sample 36–64, 66) of blood IgG background. True positive preprocessed peptides from all protease treatments, technical replicates, and proteomics software programs were pooled together. Within each of the 24 mAb-input groups, coverage metrics were computed for **(b)** the VDJ and **(c)** the CDRH3 region. **(d)** Samples without blood IgG that also contain spike-in mAbs in equal amounts (sample 36–47, 55–58) were further subdivided into single mAb (sample 36–47) and multiple mAb samples (sample 55–58). Coverage metrics were computed for **(e)** the VDJ and **(f)** the CDRH3 region. Segment line represents the same experimental replicate. Pairwise differences were determined using the Mood’s median test, with p-values adjusted for multiple testing by Benjamini–Hochberg correction. All adjusted p-values that were statistically significant (p.adj < 0.05) were displayed above each bracket as symbols: *(p.adj < 0.05), **(p.adj < 0.01), ***(p.adj < 0.001). Similar stratification and analysis for the LC is presented in **Supplementary Figure 21**.

We found that in three out of six mAbs (h9C12 Q97A, h9C12 WT, and Umab), the VDJ sequence coverage was equal to or higher in mAb-only samples compared to those with blood IgG. This difference was most pronounced at lower mAb inputs of 1 ng and 10 ng, before leveling off at 100 ng and 1000 ng (Figure 4b). However, Brimab, PGT121, and PGDM1400 were exceptions to this trend. At the lowest input amount of 1 ng, the median VDJ coverage for Brimab was 67.83%pt higher in samples with blood-derived IgG (adjusted p-value = 0.04), decreasing to increments of 63.92%pt, 55.99%pt, and 12.18%pt with increasing input amounts. Similarly, for PGDM1400, the difference in median VDJ coverage between samples with and without blood IgG was 3.9%pt, 13.83%pt, 10.28%pt, and 6.39%pt at 1 ng, 10 ng, 100 ng, and 1000 ng, respectively, although the differences were not statistically significant. This can be partially attributed to the sequence of Brimab and PGDM1400 themselves, which are made up of common human V genes, IGHV3-33 and IGHV1-2, respectively (Supplementary Table 1). Due to the frequent occurrence of these V genes in the population ^78^, it is likely that blood IgGs also possess them, thereby impacting the VDJ coverage of Brimab and PGDM1400. Furthermore, the highest number of false positive peptides removed from blood-containing samples corresponded to Brimab and PGDM1400, suggesting the impact of blood IgGs on their identification (Supplementary Figure 10). On the other hand, for PGT121, the median VDJ coverage for samples without blood at 1 ng and 10 ng of input was 6.81%pt and 2.65%pt higher than samples with blood, respectively; but at higher inputs of 100 ng and 1000 ng, the pattern reversed, with samples containing blood showing median VDJ coverage that was 21.21%pt and 4.91%pt higher than samples without blood. These results suggest the effect of blood IgG on sequence coverage is not uniform due to the polyclonal nature of the blood-derived IgGs.

CDRH3 coverage followed similar patterns to VDJ coverage when examined with and without blood IgG background. At 1 ng input, blood IgG-containing samples showed no detectable CDRH3 peptides for five of the six mAbs (h9C12 Q97A, h9C12 WT, Brimab, PGT121, and PGDM1400), whereas this occurred in only three of the mAbs for mAb-only samples (Brimab, PGT121, and PGDM1400), (Figure 4c). In addition, in most cases, the median CDHR3 coverage was equal to or lower in samples with blood IgG compared to those without, except for PGDM1400, where its longer CDRH3 (36 amino acids) may have increased the probability of non-specific sequence alignment (Supplementary Table 1). At higher mAb input levels of 100 ng and 1000 ng, PGDM1400 samples with blood IgG 4.17%pt and 8.33%pt higher median CDRH3 coverage, respectively, compared to samples without blood IgG. These findings collectively highlight the role of background polyclonal antibodies in obscuring CDRH3 peptide detection, with the magnitude of this effect diminishing at higher mAb concentrations.

After observing that high-diversity, i.e., polyclonal, sequence background decreases the CDRH3 detection accuracy, we asked whether the number of mAb spike-ins in the sample also impacts sequence coverage. To this end, we only investigated those samples without blood IgG background, and subdivided samples into those containing a single mAb (samples 36–47, containing either h9C12 WT, or Umab, or PGDM1400) versus multiple mAbs (samples 48–64, 66, containing all six mAbs at equal input amount) (Figure 4d). We found that for Umab and PGDM1400, having a single mAb present led to higher VDJ and CDRH3 coverage in most cases compared to samples with multiple mAbs (Figure 4e, f). However, h9C12 WT displayed higher sequence coverage in mixed mAb samples, likely due to the peptide alignment overlap with the mutant h9C12 Q97A, which shares all but one amino acid with the wild type (Supplementary Table 1). Similar results were observed for the LC sequence coverage (Supplementary Figure 21). Overall, our findings suggest that the sample complexity may negatively impact antibody sequence coverage.

### Mass spectrometry signal intensity is a biased estimator for mAb input quantity

Each peptide identified from the mass spectrum has a defined amino acid sequence and quantitative data related to mass peak signal intensity. Quantifying specific peptides of mAbs within a polyclonal mixture using MS1 signal intensity would provide information about the relative abundance of clonotype peptides of interest in the serum antibody repertoire. Therefore, we aimed to determine whether the signal intensities of measured peptides can accurately estimate the corresponding quantities of the original mAbs, allowing the inference of the relative abundance of mAbs in samples, with the expected linear relationship of the two variables in log-log scale to have a slope of 1 (see *Intensity ratio* section in Methods). To this end, we considered peptide pairs with identical amino acid sequences that mapped to the same mAb, originated from a single experimental replicate, and received the same protease treatment, but differed in the mAb input amount. Furthermore, within each pair, both peptides either came from samples containing blood IgG background or from mAb-only samples to prevent blood IgG-derived peptides from affecting the peptide pairs unevenly (Figure 5a). Additionally, h9C12 WT and h9C12 Q97A were excluded from this analysis because their nearly identical sequences would skew the MS1 signal intensity ratio on the peptide level, and peptides derived from the protease AspN were excluded due to their low number (Figure 2c, Supplementary Figure 22). We confirmed the a priori feasibility of MS1 signal intensity to estimate protein input amount using standard BSA protein tryptic digest (Supplementary Figure 2).

**Figure 5:**
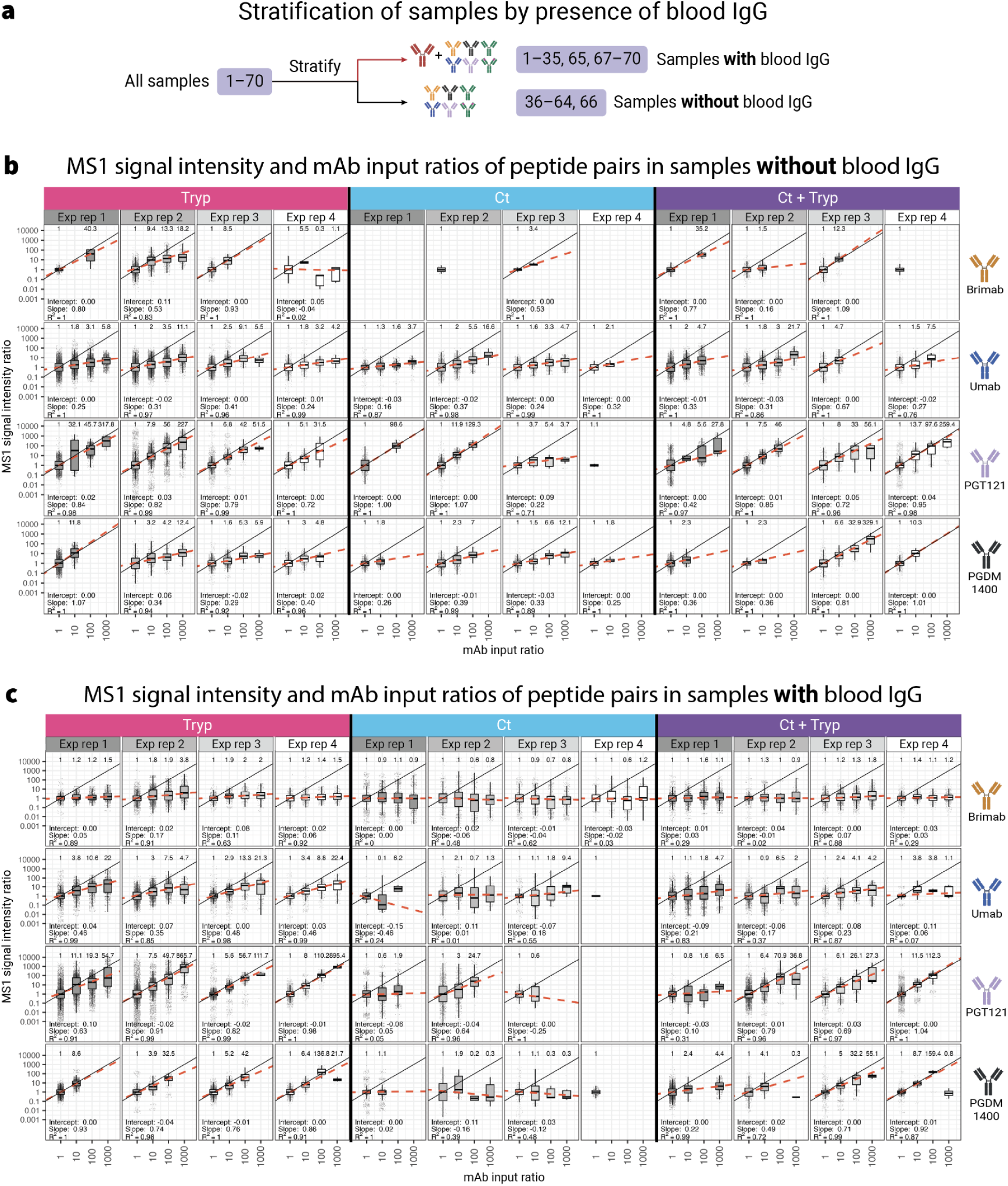
MS1 signal intensity ratio is a biased estimator of relative mAb input amount, and is biased by presence of blood IgG background, mAb sequence, and proteases utilized in digestion. **(a)** We stratified 70 samples based on the presence (sample 1–35, 65, 67–70) or absence (sample 36–64, 66) of blood IgG background. True positive preprocessed peptides from all protease treatments, technical replicates, and proteomics software programs (MQ and MSF only, as Casanovo does not report MS1 signal intensity for each peptide) were pooled together. Pairs of HC peptides aligning to the same mAb from the same experimental replicate (here denoted as exp rep for brevity) with identical amino acid sequences were formed and grouped according to the ratio of mAb input (1:1 to 1000:1, denoted as 1 to 1000 on the axis labels), and their reported MS1 signal intensity ratio was calculated (see Methods). h9C12 WT and h9C12 Q97A were excluded from this analysis because their nearly identical sequences would skew the MS1 signal intensity ratio on the peptide level. Peptides derived from ApsN were also excluded from this analysis due to a lack of data points. **(b)** MS1 signal intensity ratios in relation to mAb input ratios in peptide pairs from samples without blood IgG. **(c)** MS1 signal intensity ratios in relation to mAb input ratios in peptide pairs from samples with blood IgG. The values of MS1 signal intensity ratios and mAb input ratios were log10-transformed, and the median MS1 signal intensity ratios of each mAb-experimental replicate group were used for linear regression, weighted by the number of data points in each mAb input ratio. The dashed bright orange line represents the linear model that predicts the MS1 log10-signal intensity ratios from the log10-mAb input ratio, while the black solid represents the expected linear relationship of the two variables in log-log scale (slope 1, intercept 0). The median MS1 signal intensity ratio for each mAb input ratio is shown above each boxplot, while the intercept, slope, and the goodness-of-fit of the linear model – denoted by the coefficient of determination (R^2^), are shown below each boxplot. MS1 signal intensity ratios by mAb input for peptides on the LC is shown in Supplementary Figure 23.

For paired samples without blood IgG and equivalent mAb inputs (1:1 ratio), the median of signal intensity ratios conformed closely to the input ratio (Figure 5b). However, as the input differences increased (10:1 to 1000:1), the median signal intensity ratios deviated further from the input ratios, with the most drastic deviation occurring at 1000:1, where the median signal intensity ratios ranged 1.1–329.1 (Figure 5b). This result aligns with the findings from our preliminary experiment using standard BSA digest, where the median MS1 signal intensity ratio was also 342.8–368.5 at 1000:1 input ratios (Supplementary Figure 2b). Of note, since the mAb input range in our study varies from 1 ng to 1000 ng, peptide pairs at the highest 1000:1 input ratio are only observed when the same peptide sequence is detected at both the 1 ng and 1000 ng levels. Given the low number of detected peptides at the 1 ng level (Figure 2b), this may introduce potential bias in the estimation of the signal intensities at the 1000:1 input ratio.

The signal intensity ratio bias also varied by mAb and protease in samples without blood IgG (Figure 5c). Specifically, for PGT121 and PGDM1400, there was a clear upward trend in signal intensity ratios corresponding to increasing input ratios, from 1:1 up to 1000:1. The regression lines for PGT121 had slopes ranging 0.22–0.95, while for PGDM1400, they ranged 0.25–1.1. For Umab, the regression lines showed slopes ranging 0.16–0.67, and for Brimab, the slopes ranged 0.16–1.1, with the 1.1 slope value largely due to the absence of two out of four mAb inputs (Figure 5c). Among the proteases, Tryp provided the most peptide pairs across different mAb input ratios, followed by Ct+Tryp, while Ct had several missing mAb input ratios, and AspN was excluded due to insufficient data points overall. Notably, at the highest contrast of 1000:1 input ratio, the median signal intensity ratios were predominantly well below 1000, and, in some cases, fell beneath those observed at 1:1 input ratios (0.9–1.5 for Brimab, 1.1–22 for Umab, 6.5–895.4 for PGT121, and 0.8–55.1 for PGDM1400).

A similar trend was observed in blood-containing samples, although with greater divergence from the anticipated MS1 signal intensity ratios, leading to regression line slopes with reduced values. PGT121, PGDM1400, and Umab regression lines had slopes ranging −0.25–1, −0.16–0.93, and −0.46–0.48, respectively. For Brimab in particular, the MS1 signal intensity ratios did not change even with increasing mAb input. This is evident from the consistent persistence of the median MS1 signal intensity ratios between 0.6 and 3.8, despite the rise in the mAb input ratio, with slopes ranging −0.0073–0.17 (Figure 5c) Similar results were observed for the MS1 signal intensity ratios of LC peptides (Supplementary Figure 23). To summarize, we demonstrate that inferring relative abundance of specific mAbs through label-free quantification has promising but limited feasibility, especially in samples with polyclonal Ig background.

### *De novo* peptide sequencing demonstrates lower performance compared to peptide-spectrum matching but enables template-based V(D)J sequence assembly under certain experimental conditions

In our efforts to deduce peptide sequences from MS data, we employed not only the standard PSM proteomics platforms such as MaxQuant and MSFragger, but also integrated a machine learning-based *de novo* sequencing tool called Casanovo (Figure 1b, Supplementary Figure 9). The potential advantage of *de novo* sequencing tools is the identification of peptide sequences without the requirement of a reference database. Our objective was to assess the uniqueness of peptides identified by *de novo* sequencing compared to those from PSM methods. Of note, given that InstaNovo, another *de novo* sequencing tool ^46^, showed comparable performance to Casanovo in pilot experiments, we proceeded with only Casanovo (Supplementary Figure 24). Each peptide *de novo* sequenced by Casanovo is assigned a prediction score based on how confident the model is on average for each amino acid it has predicted in a peptide sequence (Figure 6a). Peptides derived from Tryp were predicted using the tryptic peptide model, while peptides derived from Ct, Ct+Tryp, and AspN were predicted using a non-enzymatic model. We chose the score threshold of ≥0.8 (Figure 6b) in the preprocessing pipeline to retain peptides for downstream analyses (described in Supplementary Figure 9).

**Figure 6:**
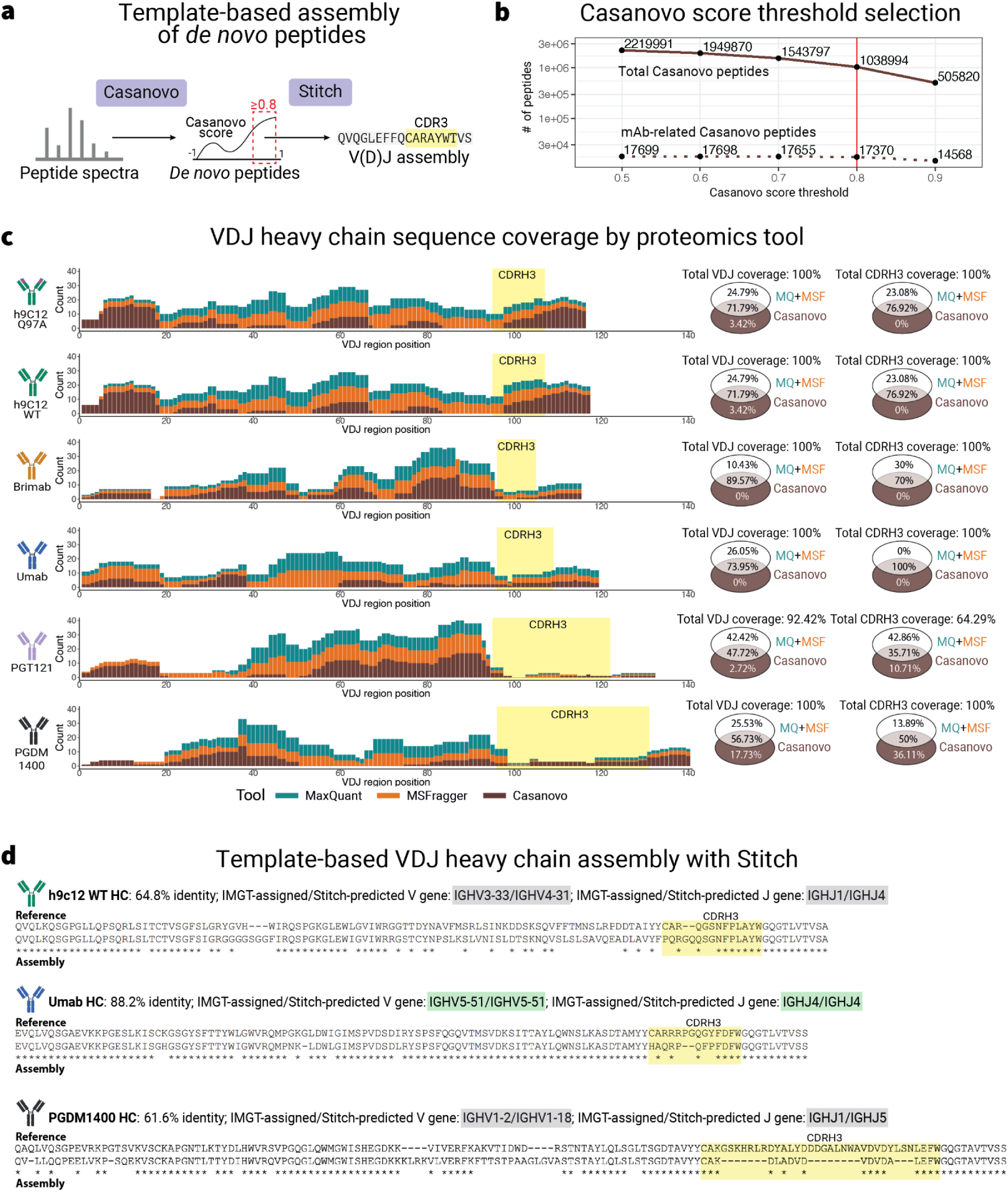
De novo peptide sequencing demonstrates lower performance compared to peptide-spectrum matching but enables template-based V(D)J sequence assembly under certain experimental conditions. **(a)** Each peptide *de novo* sequenced by Casanovo is assigned a prediction score; Tryp peptides were predicted from the tryptic model, while Ct, Ct+Tryp, and AspN peptides were predicted from a non-enzymatic model. Peptides were pooled from all experimental and technical replicates, mAb inputs, and protease digestions. All Casanovo peptides with prediction score ≥0.8 were subsequently used in template-based sequence assembly by Stitch **(b)** Number of total Casanovo peptides compared to number of mAb-related Casanovo peptides at different score thresholds from 0.5 to 0.9. We chose the prediction score threshold of ≥0.8 in the preprocessing pipeline to retain peptides for downstream analyses (described in Supplementary Figure 9). **(c)**. VDJ and CDRH3 (in yellow) coverage per tool — MQ (green), MSF (orange), and Casanovo (brown). Venn diagrams demonstrate the percentage of VDJ and CDRH3 positions covered by PSM tools (MQ+MSF) and Casanovo. The number of positions on the HC & LC covered by each proteomics software was shown in Supplementary Figure 28 and Supplementary Figure 27. Alignment of Casanovo peptides to the corresponding mAb reference is shown for the HC in Supplementary Figure 25 and for the LC in Supplementary Figure 26, with further breakdown by mAb input and presence of blood IgG in Supplementary Figure 29. **(d)** Template-based assembly of *de novo* sequencing Casanovo peptides from mAb-only samples (samples 36–47) by Stitch. Stitch predicted the most probable germline V and J genes, and reconstructed the V(D)J sequence, using the IMGT germline database as a template^79^. The assembled sequences were aligned to the corresponding mAb reference sequence. V and J genes predicted by Stitch were compared with IMGT-assigned ones, with overlapping predictions shown in green and non-overlapping predictions shown in gray. The yellow region represents the CDRH3, the “-” symbol represent a gap in the sequence, and the “*” symbol represents matching amino acids. Assembly of V(D)J sequences for both the HC and LC for PGDM1400, Umab, and h9C12WT is shown in Supplementary Figure 30.

Analysis of the sequence coverage across different proteomics software tools demonstrated substantial differences in their ability to cover the VDJ and CDRH3 regions. We observed that Casanovo provided less coverage than PSM tools (MQ+MSF), with the total VDJ sequence coverage provided by Casanovo peptides ranged 69.70–89.57%, while VDJ sequence coverage provided by PSM tools (MQ+MSF) ranged 82.26–100% (Figure 6c, Supplementary Figure 25). In particular, 0–17.73% of the VDJ region was covered solely by Casanovo, in contrast to 10.43–42.42% of the VDJ sequence covered exclusively by PSM tools, and 47.72–89.57% covered by all three tools (Figure 6c). Similar patterns were observed in the CDRH3 region, with the total CDRH3 sequence coverage provided by Casanovo peptides ranged 39.29–100%, while CDRH3 sequence coverage provided by PSM tools (MQ+MSF) ranged 63.89–100% (Figure 6c, Supplementary Figure 25), in which 0–36.11% of the CDRH3 sequence covered exclusively by Casanovo, 42.86–42.42% of covered solely by PSM tools, and 35.71–100% covered by all three tools. On a per-position basis, Casanovo exhibited gaps in coverage totaling 12–31 amino acids in the VDJ region, and 3–8 amino acids in the CDRH3. However, for PGT121 and PGDM1400, Casanovo peptides covered additional 3–13 positions in their CDRH3 that were not covered by PSM tools (Figure 6c, Supplementary Figure 27a). For the LC, where sequence coverage for the VJ and CDRL3 was generally higher than those of the HC sequence for all tools (Supplementary Figure 13), Casanovo did not provide any additional coverage, and exhibited gaps in coverage totaling 23–42 amino acids for the VJ region and 2–9 amino acids for the CDRL3 (Supplementary Figure 27b). Coverage of peptide positions by each proteomics tool are outlined in Supplementary Figure 28, and for Casanovo peptides only in Supplementary Figure 25 and Supplementary Figure 26. Further coverage analysis showed that Casanovo achieved its highest coverage at higher mAb input levels (1000 ng), both in presence and absence of blood IgG. (Supplementary Figure 29). In contrast, PSM tools reached their peak coverage at lower inputs (10 or 100 ng) and were less impacted by blood IgG (Supplementary Figure 29).

To investigate whether Casanovo peptides can be assembled into a full length HC or LC antibody sequence without prior knowledge of ground truth sequences, we used the software Stitch ^22^, which was specifically designed for antibody peptides sequence assembly based on pre-defined templates (template-based). In contrast to all prior analyses done with Casanovo peptides which were based on mAb-related peptide, the entirety of the *de novo* peptides produced by Casanovo with prediction score ≥0.8 were used for Stitch-based template-based assembly ^22^. Briefly, Stitch matches *de novo* (Casanovo) peptides to V and J genes germline sequences from the IMGT database ^79^ (Figure 6a). For Stitch assembly, we selected samples consisting of only one mAb (samples 36–47), which comprise three out of six mAbs (PGDM1400, Umab, and h9C12 WT) (Figure 2a, Figure 6d, Supplementary Figure 30). This is done because in polyclonal mixtures, reconstructing the CDR3 sequence would be especially difficult for Stitch, since it is mostly designed for monoclonal samples. Additionally, the results would require extensive manual fine-tuning and validation. The fidelity of the assembled sequence was observed to be heavily reliant on the sequence coverage in most cases. The Stitch-assembled VDJ sequences exhibited sequence identities of 64.8%, 88.2%, and 61.6% to the reference sequences for h9C12 WT, Umab, and PGDM1400, respectively. Discrepancies between the assembled sequences and the reference for h9C12 WT and PGDM1400 were found at positions where Casanovo peptides also lacked coverage (Figure 6d, Supplementary Figure 25). However, this association did not hold for Umab, as despite having coverage gaps in the VDJ region (primarily from position 39–59), the assembled sequence still maintained a high sequence identity to the reference (Figure 6d, Supplementary Figure 25). This may be owed to the fact that Stitch predicted the V and J genes in agreement with IMGT’s gene assignments for Umab (IMGT-defined/Stitch-predicted: IGHV5-51/IGHV5-51 & IGHJ4/IGHJ4), while being incorrect for h9C12 WT (IMGT-defined/Stitch-predicted: IGHV3-33/IGHV4-31 & IGHJ1/IGHJ4) and PGDM1400 (IMGT-defined/Stitch-predicted: IGHV1-2/IGHV1-18 & IGHJ1/IGHJ5) (Figure 6d, Supplementary Figure 30).

For the CDRH3 region, which rarely resembles its germline configuration, Stitch relied on overlapping peptides with residues that start or end outside the CDRH3. Consequently, peptides entirely within the CDRH3, which are crucial for high CDRH3 sequence coverage, are difficult to utilize for Stitch assembly, posing challenges for long CDRH3 sequences such as those found in PGDM1400 (Supplementary Table 1). This reliance led to numerous gaps and errors in the Stitch-assembled CDRH3 sequence, such as the starting 3 residues in h9C12 WT CDRH3 where Casanovo had a coverage gap, or the missing residues in Umab (2 aa) and PGDM1400 (18 aa) CDRH3 despite high sequence coverage due to peptides being fully within the CDRH3 (Figure 6c, Figure 6d, Supplementary Figure 25). In contrast, on the LC, where the task of template-based assembly is comparatively easier due to the lack of the highly variable D gene, the sequence identity between the assembly and the reference was comparatively higher, with sequence identity for h9C12 WT, Umab, and PGDM1400 being 67.0%, 92.5%, and 96.4%, respectively, the assembled sequences had fewer errors in the CDRL3 sequence (Supplementary Figure 30). Nonetheless, Stitch demonstrated commendable accuracy in sequence assembly, especially considering it was performed without prior knowledge of the mAb sequences.

In conclusion, *de novo* sequencing tools such as Casanovo are as yet insufficient for achieving comprehensive antibody sequence coverage, as evidenced by its inferior performance in comparison to PSM tools. However, *de novo* sequencing presents opportunities for template-based assembly.

## Discussion

To better understand the principles by which the adaptive immune repertoire evolves, there is a need to profile the serum antibody repertoire ^18,21^, which is one of the cornerstones of the adaptive immune system. Here, we investigated the performance and limitations of bottom-up proteomics-based serum antibody repertoire profiling (Ab-seq). Briefly, Ab-seq profiling of mAbs in polyclonal antibody mixtures is affected by extensive variation across different antibody sequences, proteases, input amounts and computational tools. Importantly, the observations made necessitated prior knowledge of the mAb sequence studied, demonstrating the importance of ground truth data for systems biology benchmarking^57^.

### Determination of replicability of antibody LC-MS/MS experiments using technical triplicates, experimental quadruplicates and freely available software

Ab-seq replicability is one of the key focus areas of this study. We demonstrated minimal variability across technical replicates (Supplementary Figure 4), chromatography method (Supplementary Figure 2) and consumables (Supplementary Figure 3). This technical consistency can broadly be attributed to the precision of the MS instrumentation and the sample preparation process. It lays the foundation for this study’s findings on the influence of various experimental factors on LC/MS-MS performance, such as antibody amino acid sequence, antibody concentration, protease, and polyclonal antibody complexity levels.

In contrast to technical replicates, individual experimental replicates revealed a substantial level of variation. This is underscored by our findings that show not just variability in the number of shared peptides (Supplementary Figure 20), but also in sequence coverage (Supplementary Figure 18). We showed that combining output from multiple experimental replicates considerably enhances sequence coverage (Figure 3), indicating that combining such replicates may bridge inconsistency gaps imparted by variable digestion efficiency.

By assessing the performance of proteomic analysis tools utilizing peptide-spectrum match, we found that MaxQuant and MSFragger generally yielded comparable results when operated under default settings. Nonetheless, a discernible difference emerged in the propensity for false positives, where MSFragger exhibited a higher rate of false positive peptides than MaxQuant. This effect was more pronounced or V(D)J peptides and less for CDR3 peptides (Supplementary Figure 7, Supplementary Figure 8), suggesting that the filtering criteria in each software suite influence the accuracy of peptide identification. This potential divergence between the software platforms emphasizes the necessity of stringent validation measures, such as filtering for false positives, contaminants, and common peptide sequences among the six mAbs when interpreting Ab-seq data. Consequently, further development of antibody-focused mass spectrometry software tools, along with concerted community efforts to measure and benchmark their performance, is crucial to advancing the accuracy and reliability of Ab-seq.

### Observed technological limitations of Ab-seq

In our study, the LC-MS/MS analysis was performed using an Evosep One liquid chromatography system connected to a high-resolution quadrupole-Orbitrap mass spectrometer. Given that our benchmarking study included 3360 MS/MS samples, using a standard nano-LC would have been time-consuming and impractical, typically allowing for a throughput of only 14–20 samples per day. In addition, the small microscopic diameter of the nano-LC columns makes them prone to clogs and breakage, which would severely disrupt the workflow. Meanwhile, the Evosep One system is both robust and high throughput – tested in this study with up to 100 samples per day (Supplementary Figure 2), and was thus the LC instrument of choice for our experiments. However, this setup, combined with the stochastic nature of peptide precursor selection in data-dependent acquisition (DDA), raises inherent challenges in mass spectrometry-based proteomics. While this technique holds substantial analytical power, it is nonetheless constrained by the propensity for multiple peptide species to co-elute within a single MS1 scan. When a multitude of peptides are present, DDA inherently favors the most abundant peaks for MS2, leading to a systematic under-representation of low-abundance peptides ^80,81^. As a result, divergent fragmentation subgroups across different samples are created and introduces significant variability between replicates, particularly evident through the diminished identification of low abundant peptides and, consequently, incorrect quantification of peptides in complex mixtures ^82,83^ (Figure 5). Such variability underscores the limitations of reliance on DDA for comprehensive proteomic profiling, highlighting the necessity for advanced analytical strategies to enhance peptide detection, particularly for those of lower abundance in a complex mixture. In response to these shortcomings, employing methods capable of diminishing the effects of stochastic precursor selection and interference susceptibility such as data-independent acquisition (DIA) ^84,85^, coupled with DIA-compatible software tools ^86,87^, may be important for achieving a more complete and reproducible capture of the secreted antibody repertoire in future studies ^88,89^.

### Antibody peptide discovery is protease and Ab-dependent

The characterization of antibody peptide sequences using Ab-seq was found to be markedly influenced by both the specificity of protease cleavage and the intrinsic properties of the antibodies themselves (Figure 2). For example, PGT121 failed to achieve complete coverage even when peptides from all proteases were pooled, despite substantial mAb inputs of 1000 ng (Figure 2d), diverging from the *in silico* predicted digestion of the mAb sequences based on the reported protease specificities (Supplementary Figure 14). This attests to a dependency not only on input amount but also on the inherent sequence characteristics of each mAb. To enhance sequence detection, particularly within the highly variable CDR3, the strategic use of specific proteases is crucial — AspN, for instance. Although its standalone performance may not be remarkable (Figure 2c,d), AspN proved indispensable for achieving in-depth CDRH3 profiling when used in conjunction with other proteases. Similar recommendations have been made in prior studies advocating for the tandem use of AspN with other proteases for increased coverage ^74,90^. Therefore, protease diversity is a key determinant for coverage enhancement ^71,74,91^.

Protease specificities for peptide generation were also found to be of considerable importance. Specifically, allowing for controlled miscleavages through the concurrent use of chymotrypsin and trypsin — specifically with a shortened chymotrypsin digestion phase — resulted in superior overall coverage compared to using either protease individually (Figure 2d). This suggests that complementary tandem protease digestion could potentially unlock more complete peptide sequences, an approach also being explored in other studies ^90^. In addition to site-specific digestion by proteases, other studies also introduced the notion that nonspecific cleavage techniques, such as microwave-assisted digestion, can produce a diverse collection of peptide fragments ^92^. However, non-specific digestion may jeopardize the reproducibility pivotal to mass spectrometry, and thus must be weighed against the need for reliable and repeatable results.

### Polyclonal antibody background necessitates preprocessing of peptide sequence results and may impact detection of monoclonal antibody sequences

Ab-seq data collection faces the challenge of discerning true signals amidst false positives, which may arise from (i) misidentification made by proteomics software, (ii) incorrect peptide assignment due to high similarity between antibody sequences, or (iii) carryover contamination. While incorrect peptide identification can be mitigated by adjusting the false discovery rate, shared peptides between antibodies, particularly those derived from homologous non-CDR3 regions (Supplementary Figure 10), should be reassigned to the correct antibody or removed as false positives during the preprocessing steps (Supplementary Figure 9e). In addition, carryover contamination from liquid chromatography effluent can be a serious problem when multiple samples are run sequentially. However, this biological contamination can be mitigated by designing LC/MS-MS experiments where blank run cycles are interspersed after true sample runs. This approach not only helps elute any remaining antibodies from the previous run, but also by using the peptides detected in the effluent of blank cycles to filter out detectable carryover contamination (Supplementary Figure 9e).

The benchmarking nature of our study allowed us to further filter out false positive peptides, since the sequence of each mAb of interest was known beforehand. In a real-world Ab-seq study, however, the Ab sequence content of each sample is usually unknown, making the peptide preprocessing challenging. Without knowing which antibodies should be present in each sample, it is impossible to determine false positive peptides and further filter shared peptides between different antibodies. In specific cases where particular clonotypes of interest are tracked, prior knowledge of these clonotype sequences can serve as a reference, enabling filtration of false positive peptides for tracked clones. While this type of false positive filtration is typically limited, blank cleaning cycles can still help to reduce carry-over peptides in a real-world Ab-seq study, especially if cleaning cycles are performed after each true sample run.

In the context of our benchmarking study, blood-derived polyclonal IgG has been identified as a major deteriorating factor in sequence coverage performance. Not only does it artificially inflate V(D)J sequence coverage for certain mAbs (Figure 4b) — likely a consequence of shared framework region sequences with spiked-in mAbs—but it also diminishes the level of CDR3 coverage (Figure 4c). Furthermore, the interference of blood IgGs has manifested in the diminished detection of CDR3 peptides at lower input amounts (Figure 2b), an outcome attributable to spectral signal interference discussed in previous sections. Even in the absence of blood IgG components, the detection of CDR3 in samples containing a mixture of six mAbs presented lower coverage compared to individual mAb samples — albeit this reduction was not statistically significant (Figure 4e). This observation suggests that even without the confounding variable of blood IgG, the signal complexity due to multiple mAbs may impact CDR3 detection.

In conclusion, our benchmarking framework demonstrated how systematic filtering and preprocessing measures enhance the fidelity of Ab-seq data. However, challenges in detecting CDR3 sequences persists even when accounting for the blood IgGs, showcasing the challenges of accurate antibody repertoire characterization at full polyclonal complexity.

### Antibody abundance across different samples estimated using MS1 signal intensity is biased

Quantifying specific antibodies in a polyclonal mixture offers valuable insights into clonotype abundance and facilitates mAb monitoring, with MS1 signal intensity serving as an indirect metric to estimate relative antibody abundance across samples, potentially aiding in therapeutic monitoring and adaptive immune responses (e.g., post-vaccination). We showed that the MS1 signal intensity ratio captures the exponential growth in the mAb abundance across samples, though the parameters of the exponential function may be biased (Figure 5). The challenge in deducing relative antibody abundances at high input ratios, like 1000:1, is exacerbated by both scarcity of data points at 1 ng (10 ng/ml), due to the limit of quantitation of LC-MS/MS (established at 15 ng/ml in previous studies ^93^. MS1 signal saturation at 1000 ng may occur due to our choice to use EvoSep instead of other nanoLC methods, a phenomenon influenced by ion suppression effects previously documented in the literature ^94^. Additionally, label-free quantification at the peptide level, our chosen methodological approach, is limited in quantification accuracy compared to label-based methods ^95^. However, label-free quantification stands out for its simplicity, scalability, and utility — a critical aspect in the study of serum antibody dynamics at scale.

### *De novo* Ab-seq performance is currently limited but holds potential for improvements

Given the high degree of individualization and high diversity of the antibody repertoire ^96–98^, it is impossible to create a reference database that includes all existing antibodies for all individuals. This limitation makes it challenging for database-search methods to comprehensively capture the full diversity of the antibody repertoire. While BCR-seq data from an individual can be used as a reference database for LC-MS/MS analysis of the same individual, the extent of overlap between the genomic compartment (BCR) and the proteomic compartment (Ab) remains unclear ^25,29,56^. Another challenge for any reference-dependent methods is the heterogeneity between different anatomical sites, such as the blood, lymph node, and bone marrow. Thus, the ability to sequence and ideally reassemble Abs without prior knowledge is crucial for future efforts in this field. That said, PSM of MS spectra is an easier computational problem than *de novo* protein sequencing.

When using the machine learning-based *de novo* sequencing tools Casanovo, the resulting mAb-related peptides were predominantly short (Supplementary Figure 15). Additionally, Casanovo appeared to overlook numerous peptides that are successfully identified by both MQ and MSF (Figure 6c, Supplementary Figure 27). The reasons for these discrepancies merit further investigation. However, Casanovo (and potentially Instanovo, although it was only tested on a small sample size) was able to detect mAb-related peptides within the CDRH3 region with high confidence score, even where coverage in this region was low in MQ and MSF results (Figure 6c, Supplementary Figure 28).

For this work, we utilized Casanovo ^45^ for *de novo* sequencing and also tested Instanovo ^46^, albeit to a lesser extent. However, there exists several other tools capable of *de novo* sequencing, which has either been extensively compared by others ^99,100^, or novel methods yet to be benchmarked ^101–103^. In addition, modifications to the sample preparation ^43,104^ and fragmentation process ^72^ could potentially yield better results for *de novo* sequencing specifically.

### Template-based V(D)J sequence assembly can achieve reasonable accuracy, but still limited to monoclonal samples

Significant progress is still required for accurate and complete antibody sequence assembly with bottom-up proteomics. Although peptide identification efforts have approached near-complete coverage of large polypeptide chains, mapping PSMs to a reference sequence is not equivalent to *de novo* protein sequence assembly. This challenge is somewhat mitigated in the antibody space, where mapping to a known germline gene segment is possible and the region of interest (CDR3) is substantially smaller. Despite major improvements in recent years ^47,99,105^, the task of sequence assembly from *de novo* sequencing peptides remains challenging, and are even pronounced when false positives complicate the assignment of a complete sequence using standard assembly methods, or when oligo- and polyclonal mixtures are involved. Achieving successful and robust template-based assembly of *de novo* peptides will likely require a combination of comprehensive sample preparation workflows involving multiple proteases to generate overlapping fragments ^43^, confident peptide sequence assignment using *de novo* tools with accurate false discovery rate (FDR) filtering, and robust algorithms for protein assembly and selection of final assembled products.

Given the diversity inherent in the antibody variable regions, assembling peptides using a template is more practical than a completely *de novo* approach. However, it is crucial to acknowledge that template-based assembly could be biased by the references used. This issue is exacerbated by the incompleteness of current antibody germline gene databases in fully representing the genetic diversity of human and animal populations, notwithstanding ongoing efforts to mitigate this issue ^106^. Additionally, many errors in the assembled sequences are specific to mass spectrometry and frequently emerge in *de novo* sequencing results (Figure 6d). Such errors include Isoleucine/Leucine ambiguities and incorrect sequence order (e.g., SG instead of GS) when certain mass peaks exhibit low intensity, as previously documented ^107^.

### Final remarks and future outlook

This work provides a resource for antibody immunology and proteomics communities. Owing to its scope, depth, and coverage, our dataset provides a robust foundation to drive future methods that enable studying the human antibody proteome, both experimentally and computationally, including AI-driven approaches. Specifically, our dataset may be used for training and testing novel *de novo* sequencing approaches.

Our work demonstrates the importance of ground-truth data for systems biology benchmarking. In the future, it may be desirable to develop antibody-specific methods for mass spectrometry data processing, based on extensive benchmarking on ground-truth Ab samples at controlled inputs and complexity levels. Specifically, the first step in protein inference is peptide identification. Peptide identification using a reference database, as opposed to *de novo* peptide sequencing, is the most commonly used technique in Ab-seq. However, the highly degenerate antibody repertoire breaks key assumptions for controlling false PSM calls ^108^. Using standard peptide identification tools for Ab-seq may lead to higher than expected false positive PSMs and consequently a high rate of false antibody identifications.

Profiling serum antibody sequence diversity through bottom-up, top-down, or a combination of proteomics approaches is not only inherently useful as a complement for genomic profiling of BCR diversity ^16,17,61,109^ Additionally, it may be combined in the future with structural antibody profiling where sequence identification is needed to improve antibody structure reconstruction ^6,110–112^. Furthermore, sequence identification would complement antibody profiling methods such as PhIP-seq ^113^ where the binding of a polyclonal antibody mixture to a large antigen landscape is recorded in a sequence-agnostic fashion.

In conclusion, our findings underscore several critical considerations for advancing research in antibody mass spectrometry. (1) Stringent filtering of false positive peptide hits emerges as particularly crucial in the context of polyclonal antibody samples, where the probability of false positives are elevated. (2) Ensuring the use of multiple experimental replicates is imperative to enhance the reliability and reproducibility of results, reducing inherent experimental variability on the results. (3) Caution must be exercised when drawing biological inferences, particularly concerning antibody diversity, as various methodological biases may skew interpretations. (4) Estimation of mAb amount from MS1 signal intensities is feasible but biased. (5) *De novo* sequencing showed lower performance than PSM, but the difference will likely decrease with larger amounts of training data. Already now, *de novo* sequencing is *de novo* sequence assembly under certain experimental conditions. Prioritization of experimental and computational methodological refinements in future studies is necessary to propel the field towards a more accurate and comprehensive understanding of the serum antibody repertoire.

## Methods

### Monoclonal antibodies

Six monoclonal IgG1 antibodies were used: PGDM1400, PGT121, h9C12 WT ^62^, h9C12 Q97A ^62^ and recombinant forms of Ustekinumab (Umab) and Briakinumab ^114^ (Brimab). Vectors encoding the HCs and LCs of the mAbs were made and used for transient transfection of Expi293 cells grown in a serum-free system followed by purification, as previously described ^115^.

All the antibodies used are human monoclonal antibodies of the IgG1 isotype. PGDM1400 and PGT121 are antibodies against HIV envelope protein gp120, Umab and Brimab target the p40 protein subunit of IL-12 and IL-23, h9C12 WT and h9C12 Q97A bind to the AdV5 hexon protein. The range of input for spike-in mAbs ranged from 1 μg (6.67 pmol), 100 ng (0.67 pmol), 10 ng (0.067 pmol), and 1 ng (6.67 fmol).

Amino acid sequences of the recombinant mAbs’ CDR3 region are presented in Supplementary Table 1, and the V(D)J region amino acid sequences are presented Supplementary Table 2.

### Blood isolated human IgG

A whole blood sample from a healthy volunteer was used for plasma separation. Blood was collected in a K_2_EDTA vacutainer (367525, BD) and centrifuged for 15 min at 800 g. The plasma layer was then transferred into a clean vial and used for subsequent IgG purification.

Affinity purification of plasma IgG was performed using NAb Protein A/G Spin Kit (89980, Thermo Fisher Scientific) according to the manufacturer’s instructions following the spin purification protocol.

Purification of Fab fragments was performed with GingisKHAN Fab kit (B0-GFK-020, Genovis) according to the manufacturer’s protocol with a 4 hours digestion step.

### Sample preparation and LC-MS/MS analysis

For each of the four experimental replicate, samples were initially prepared in a 96-well master plate (Protein LoBind, cat no 0030504119, Eppendorf) in 0.1M ammonium bicarbonate (09830, Sigma-Aldrich) solution with subsequent aliquoting into four experimental plates. The pipetting was performed using Electronic Pipettes (Mettler-Toledo).

DTT was added to each sample well in a final concentration of 10 mM followed by incubation at 37 °C for 60 min to reduce cysteines. Cysteines were further alkylated with 15mM IAA for 30 minutes in the dark at room temperature. To the four plates 0.5 µg trypsin (Tryp), 0.5 µg chymotrypsin (Ct), 0.5 µg AspN, and a combination of 0.5 µg chymotrypsin and 0.5 µg trypsin (Ct+Tryp) per well was added, respectively. Plates with trypsin, chymotrypsin and AspN were incubated at 37 °C for 18 hours, and the Ct+Tryp treated plate was first incubated with chymotrypsin for 4 hours before trypsin was added for the 18-hour incubation (Supplementary Table 5).

After the incubation, each sample was transferred to Evotips (Evosep) in triplicates for desalting prior to the LC-MS/MS analysis. The standard protocol from the manufacturer was used.

The LC-MS/MS analysis was performed on an EvosepOne liquid chromatography system connected to a quadrupole – Orbitrap mass spectrometer (QExactive HF, ThermoElectron, Bremen, Germany) equipped with a nanoelectrospray ion source (EasySpray/Thermo). For liquid chromatography separation, an 8 cm C18 column (Dr Maisch C18 AQ, 3 μm beads, 100um ID, 8 cm long, Evosep) was used. The standard 100 samples/day method was used. One blank sample run were performed between each numbered sample, and three blank sample runs were performed between samples of different mAb spike-ins with 1000 ng input to reduce carry over contamination, and the peptide data obtained from the blank samples was utilized to remove contaminant peptides (Supplementary Figure 9).

The mass spectrometer was operated in the data-dependent mode to automatically switch between MS and MS/MS acquisition. Survey full scan MS spectra (from m/z 375 to 1,500) were acquired in the Orbitrap with resolution R = 60,000 at m/z 200 (after accumulation to a target of 3,000,000 ions in the quadruple). The method used allowed sequential isolation of the most intense multiply-charged ions, up to twelve, depending on signal intensity, for fragmentation on the HCD cell using high-energy collision dissociation at a target value of 100,000 charges or maximum acquisition time of 50 ms. MS/MS scans were collected at 30,000 resolution at the Orbitrap cell. Target ions already selected for MS/MS were dynamically excluded for 30 seconds. General mass spectrometry conditions were: electrospray voltage, 2.0 kV; no sheath and auxiliary gas flow, heated capillary temperature of 250 °C, normalized HCD collision energy 28%.

### Proteomics software analysis

We applied three freely available proteomics software tools to MS raw data — two PSM tools for peptide identification and quantification, MaxQuant ^75^ and MSFragger ^76^, and a reference free (*de novo*) tool for peptide sequencing, Casanovo ^45^. PSM search (MaxQuant and MSFragger) was performed against the following databases: (i) HC and LC sequences of the 6 mAbs, (ii) human UniProt database ^116^ from trEMBL and Swiss-Prot (downloaded March 2021), (iii) sequences of V, D, and J genes downloaded from IMGT (downloaded July 2020) ^67^, and (iv) 10000 synthetic IGH, IGK, and IGL simulated with immuneSIM ^117^.

MS raw files were submitted to MaxQuant software version 2.1.3.0 for protein identification and quantification, with parameters set to default values unless specified, and the protease option in MaxQuant matching the protease used to digest the samples. A minimal peptide length was set to 7 amino acids, with up to two allowed miscleavages. Minimal unique peptides were set to 1, and FDR allowed was 1% for peptide and protein identification. Generation of reversed sequences was selected to assign FDR rates. An example MaxQuant configuration file can be found in Supplementary File 1.

MSFragger version 3.5 was run through FragPipe version 18.0 using default values unless specified. Parameters included Carbamidomethylation as a constant modification, N-acetylation and methionine oxidation as variable modifications, minimal peptide length 7 amino acids and maximum peptide length 50 amino acids, with up to two allowed miscleavages and only fully-enzymatic termini allowed. See Supplementary File 2 for more configuration details.

*De novo* peptide sequencing was performed with Casanovo version 3.2.0 ^45^. For replicates produced with trypsin, predictions were made with a model trained on 28 million tryptic PSMs. For non-tryptic replicates, predictions were made with a non-enzymatic model fine-tuned on a dataset of 1M PSMs with uniform distributions of terminal amino acids. All predictions were made using a beam size of 5. An additional post-processing step was applied to Casanovo output peptides, where post-translational modifications (PTMs) were removed from the peptide sequences and after that the identical peptides within each sample and technical replicates were merged. The prediction score of each merged peptide was assigned as the maximum score among all the merged peptides. Finally, peptides with prediction score below 0.8 were eliminated. Training sets and training procedures for both models are described in prior works ^45,118^, and both sets of model weights are publicly available at https://github.com/Noble-Lab/casanovo/releases. The Casanovo configuration file is available as Supplementary File 3.

### Data preprocessing

Data preprocessing involved multiple steps, outlined in Supplementary Figure 9. Initially, peptide sequences from four experimental replicates and three technical replicates were merged, including Casanovo-predicted peptides, which were further processed as detailed in the “Proteomics Software Analysis” section. All identified peptides were aligned to the six mAb reference sequences, both HC and LC. Peptides that did not align to either the HC or LC sequences of the mAbs were deemed non-mAb peptides and excluded from further analysis. Peptides were considered mAb-related if they matched the reference HC or LC sequences with 100% accuracy or if they overlapped by at least 5 AAs at the beginning or end of HC or LC sequences. CDR3-related peptides were defined as mAb-related peptides overlapping by at least 3 AA within the CDR3 region.

All mAb-related peptides were categorized as either false positives (FP) or true positives (TP). A peptide was considered a false positive if it mapped to an mAb not present in the sample by design, and a true positive otherwise. False positive peptides were further filtered based on the following criteria: (i) FP peptides present in samples with blood IgG and aligning to the V, D, or J segments in the IMGT database, or (ii) FP peptides corresponding to multiple mAb, where one mAb is present in the sample design and the other is not present (Supplementary Table 5), or (iii) FP peptides detected during previous cleaning cycles (blank sample runs) interspersed between true sample runs. Only peptides that passed all filtering steps were retained for downstream analysis. Figure 2b shows results for both TP and FP peptides, whereas Figure 2b-c, Figure 3, Figure 4, Figure 5, and Figure 6 display only TP peptide results to minimize potential bias from FP peptides. Unless explicitly noted, technical replicates for each sample were merged, as these replicates showed high consistency in mass spectrometry signals (Supplementary Figure 4).

### Analytical methods

#### In silico digestion of mAb sequences

Prior to the experiment, we digested the V(D)J sequences of the six mAbs *in silico* using the R package *cleaver* ^119^, which cleaves polypeptide sequences according to the ExPASy cleavage rules^120^(Tryp, Ct, Ct+Tryp, AspN), with up to 2 miscleavages allowed. The resulting set of peptides with length of at least 6 aa were then compared with preprocessed peptides derived from experimental data.

#### Ab-seq coverage

Ab-seq coverage is calculated as the sum of amino acid positions covered by Ab-seq peptides divided by the length of the reference sequence (either the V(D)J region or the CDR3) (Supplementary Figure 12). We used Ab-seq sequence coverage and not Ab-seq peptide count as a measure of Ab-seq performance because our focus in this work was on measuring the extent of the recovery of mAb V(D)J sequences. As visualized on Supplementary Figure 12, distinct peptides can cover the same mAb positions and, therefore, meaning that the number of unique peptides may not accurately reflect the true extent of V(D)J sequence recovery. Consequently, an increasing Ab-seq peptide count may not lead to higher recovery of the antibody V(D)J sequence.

#### Intensity ratio

To calculate the MS1 intensity ratio, we selected pairs of peptides aligning to the same mAb with identical amino acid sequences from different MS samples within the same experimental replicate, which were digested using the same protease treatment and were identified using the same tool — either MaxQuant or MSFragger. Additionally, within each pair, both peptides belonged to either blood-containing samples or mAbs-only samples. We did not restrict the peptides to contain the same modification, as most peptides were unmodified (Supplementary Figure 22).

The intensity ratio for each peptide pair was calculated as the ratio between the MS1 intensity of a peptide (as reported from the output of MaxQuant and MSFragger) from a sample with higher mAb input and that from a sample with lower mAb concentration. In this case, the true concentration ratio will correspond to 1:1, 10:1, 100:1, or 1000:1 input ratios. We excluded h9C12 WT and h9C12 Q97A from the intensity ratio analysis because they co-occurred in most samples and the intensity of peptides not covering position Q97A will sum up from both antibodies, thereby distorting the expected 1:1, 10:1, 100:1, 1000:1 input ratios. Peptides cleaved by AspN were also excluded due to their low abundance (Supplementary Figure 22).

For each mAb, protease treatment, and experimental replicate, we calculated the median of log10-transformed intensity ratio within each concentration ratio subgroup (1:1, 10:1, 100:1, 1000:1). To assess how accurately the intensity ratio estimates the concentration ratio, we performed a weighted linear regression using the four log10-transformed MS1 signal intensity ratio medians and the four log10-transformed mAb input amounts, the regression formula is calculated as log(y) = intercept + slope*log(x). The weights of the regression were determined by the number of data points used to calculate each median. If the MS1 signal intensity ratios were perfectly concordant with the mAb input ratios, the regression line would have a slope of 1 and intercept of 0.

#### Template-based assembly of de novo peptides into V(D)J sequence

Only samples with one mAb present and no blood IgG background (samples 36–47) were utilized for assembly with Stitch version 1.5, which was run in monoclonal mode ^22^. All Casanovo peptides with prediction score ≥0.8 were used as input for Stitch. Template-matching of peptides using mass-based alignments utilized the IMGT database internal to Stitch, with cut-off score 15 (mean segment score per position needed for a path to be included), enforce unique threshold 0.9, and ambiguity threshold 0.9. Assembly of aligned peptides into V(D)J sequence utilized the highest scoring V, (D), and J gene segments, with a cut-off score of 10 (mean score per position required for the gene segment to be included in the recombination). The resulting assembled sequence was then aligned with the reference mAb sequence using the SIM-Alignment Tool for Protein Sequences (https://web.expasy.org/sim/), and sequence identity percentage was calculated. An example Stitch configuration file for h9C12 WT is available as Supplementary File 4.

### Graphics

Plots were generated using the R packages ggplot2 v.3.4.2 ^121^, ComplexHeatmap v.2.2.0 ^122^, and UpSetR ^123^. Figure 1 (https://BioRender.com/s33n780), Figure 2a (https://BioRender.com/n55a473), and Supplementary Figure 12 (https://BioRender.com/e25s512) were made using BioRender.com. Plots were arranged using Adobe Illustrator 2023 v.27.4.1.

### Data and code availability

All mass spectrometry data are available online via ProteomeXchange with identifier PXD055846, including raw files, MQ, MSF, and Casanovo output files, parameter files, search database files, and sample descriptions files. Processed data and corresponding scripts are available on github: https://github.com/csi-greifflab/Systematic-benchmarking-of-mass-spectrometry-based-antibody-sequencing-reveals-methodological-biases

## Acknowledgements

We thank Dr. Avinash Yadav (GSK, Siena, Italy) for valuable suggestions early in the project. We are grateful to Dr. Albert Bondt (Utrecht University, The Netherlands) for helpful input on the manuscript; Dr. Joost Snijder (Utrecht University, The Netherlands) and Douwe Schulte (Utrecht University, The Netherlands) for productive discussions on *de novo* sequencing.

## Supplementary Materials

**Supplementary Table 1:** IMGT Annotation of the six monoclonal antibodies (V and J gene name, CDR3H/L regions) used in the study.

| Antibody | Chain | CDR3 aa sequence | V gene | J gene |
| --- | --- | --- | --- | --- |
| Briakinumab (Brimab) | Brimab_HC | CKTHGSHDNW | IGHV3-33 | IGHJ1 |
|  | Brimab_LC | CQSYDRYTHPALLF | IGLV1-40 | IGLJ6 |
| Ustekinumab (Umab) | Umab_HC | CARRRPGQGYFDFW | IGHV5-51 | IGHJ4 |
|  | Umab_LC | CQQYNIYPYTF | IGKV1D-16 | IGKJ2 |
| PGT121 | PGT121_HC | CARTLHGRRYIGIVAFNEWFTYFYMDVW | IGHV4-4 | IGHJ6 |
|  | PGT121_LC | CHIWDSRVPTKWVF | IGLV3-21 | IGLJ3 |
| PGDM1400 | PGDM1400_HC | CAKGSKHRLRDYALYDDD GALNWAVD VDYLSNLEFW | IGHV1-2 | IGHJ1 |
|  | PGDM1400_LC | CMQGRESPWTF | IGKV2-29 | IGKJ1 |
| h9C12 WT | h9C12-WT_HC | CARQGSNFPLAYW | IGHV3-33 | IGHJ1 |
|  | h9C12_LC | CHQDYSSPLTF | IGKV4-1 | IGKJ4 |
| h9C12 Q97A | h9C12-Q97A_HC | CARAGSNFPLAYW | IGHV3-33 | IGHJ1 |
|  | h9C12_LC | CHQDYSSPLTF | IGKV4-1 | IGKJ4 |

**Supplementary Table 2:**
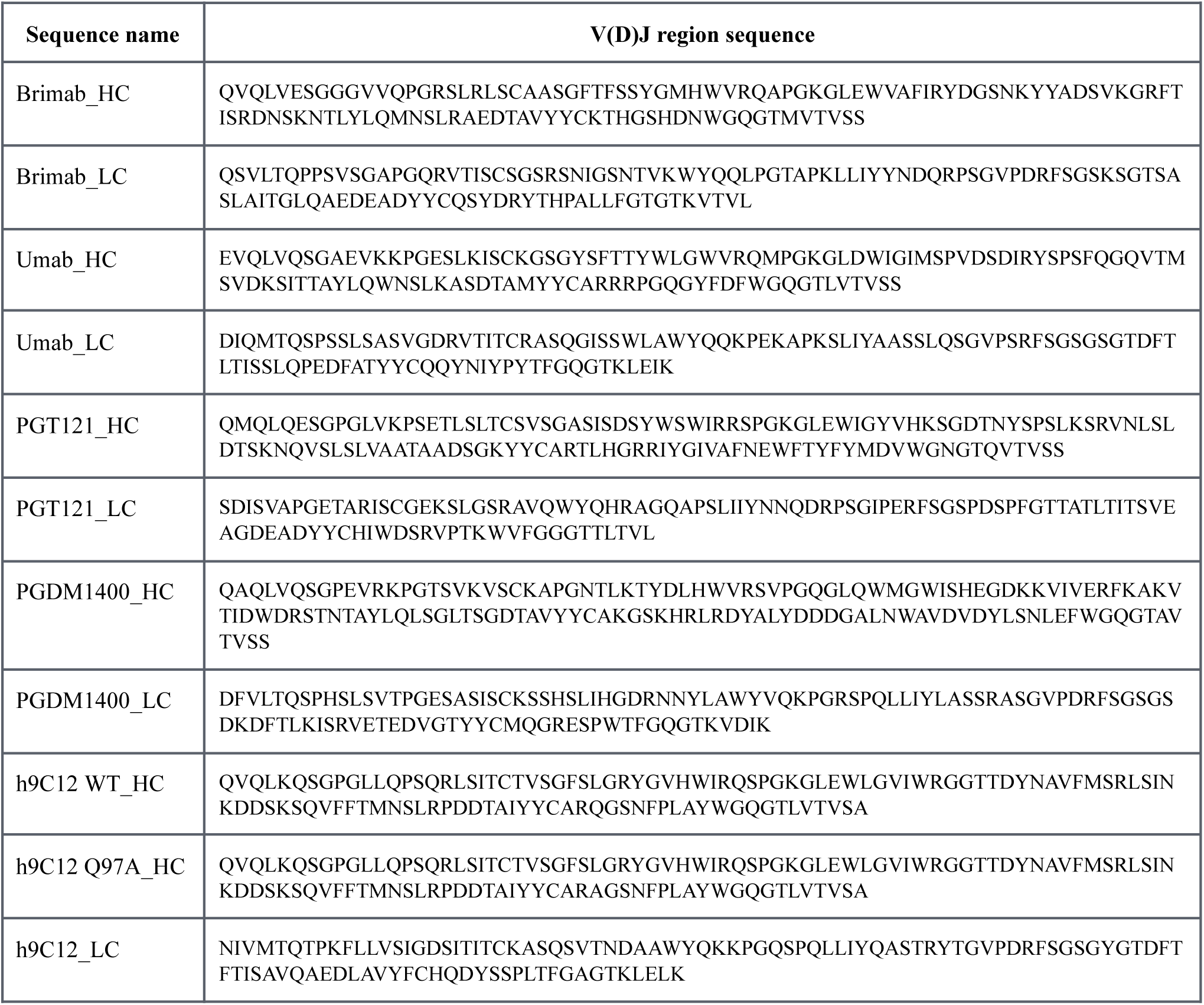
Antibody sequences used in the study.

**Supplementary Table 3:** Pairwise Ab sequence identity between the V(D)J, CDR3, and non-CDR3 (V(D)J except CDR3) regions. Sequence identity (as percentage) is calculated after pairwise alignment of the Ab sequences by the following formula:

| Antibody 1 | Antibody 2 | Identity of V(D)J region (%) | Identity of CDR3 region (%) | Identity of non-CDR3 region (%) |
| --- | --- | --- | --- | --- |
| PGDM1400_HC | PGDM1400_LC | 25.52 | 13.04 | 29.73 |
| PGDM1400_HC | PGT121_HC | 34.04 | 20.59 | 38.32 |
| PGDM1400_HC | PGT121_LC | 27.41 | 21.05 | 31.68 |
| PGDM1400_HC | Umab_HC | 44.68 | 17.65 | 53.27 |
| PGDM1400_HC | Umab_LC | 21.28 | 10.34 | 24.3 |
| PGDM1400_HC | Brimab_HC | 42.55 | 13.33 | 52.34 |
| PGDM1400_HC | Brimab_LC | 27.59 | 25 | 30.63 |
| PGDM1400_HC | h9C12 WT_HC | 34.51 | 40 | 40.19 |
| PGDM1400_HC | h9C12 Q97A_HC | 34.75 | 40 | 40.57 |
| PGDM1400_HC | h9C12_LC | 24.11 | 17.86 | 26.17 |
| PGDM1400_LC | PGT121_HC | 30.28 | 20 | 33.62 |
| PGDM1400_LC | PGT121_LC | 41.59 | 8.33 | 44.55 |
| PGDM1400_LC | Umab_HC | 31.25 | 8.33 | 32.76 |
| PGDM1400_LC | Umab_LC | 54.46 | 33.33 | 56.31 |
| PGDM1400_LC | Brimab_HC | 27.87 | 11.11 | 29.2 |
| PGDM1400_LC | Brimab_LC | 46.96 | 0 | 50.49 |
| PGDM1400_LC | h9C12 WT_HC | 33.33 | 30 | 33.91 |
| PGDM1400_LC | h9C12 Q97A_HC | 32.54 | 20 | 33.91 |
| PGDM1400_LC | h9C12_LC | 53.57 | 44.44 | 54.37 |
| PGT121_HC | PGT121_LC | 25.69 | 18.18 | 27.97 |
| PGT121_HC | Umab_HC | 40.6 | 30.77 | 42.99 |
| PGT121_HC | Umab_LC | 25.53 | 16.67 | 27.83 |
| PGT121_HC | Brimab_HC | 39.85 | 15.38 | 45.79 |
| PGT121_HC | Brimab_LC | 29.41 | 21.43 | 32.73 |
| PGT121_HC | h9C12 WT_HC | 43.94 | 26.09 | 49.06 |
| PGT121_HC | h9C12 Q97A_HC | 44.27 | 26.09 | 49.52 |
| PGT121_HC | h9C12_LC | 20.44 | 9.09 | 23.58 |
| PGT121_LC | Umab_HC | 31.15 | 0 | 32.71 |
| PGT121_LC | Umab_LC | 38.1 | 11.11 | 41.94 |
| PGT121_LC | Brimab_HC | 30.58 | 12.5 | 32.11 |
| PGT121_LC | Brimab_LC | 50.44 | 8.33 | 55.45 |
| PGT121_LC | h9C12 WT_HC | 26.27 | 16.67 | 27.36 |
| PGT121_LC | h9C12 Q97A_HC | 26.27 | 18.18 | 27.36 |
| PGT121_LC | h9C12_LC | 35.4 | 33.33 | 36.63 |
| Umab_HC | Umab_LC | 31.2 | 8.33 | 33.63 |
| Umab_HC | Brimab_HC | 51.26 | 16.67 | 55.14 |
| Umab_HC | Brimab_LC | 32 | 0 | 35.4 |
| Umab_HC | h9C12 WT_HC | 43.9 | 26.67 | 45.37 |
| Umab_HC | h9C12 Q97A_HC | 42.86 | 16.67 | 45.79 |
| Umab_HC | h9C12_LC | 24.17 | 16.67 | 25.93 |
| Umab_LC | Brimab_HC | 29.37 | 0 | 31.62 |
| Umab_LC | Brimab_LC | 50 | 33.33 | 52 |
| Umab_LC | h9C12 WT_HC | 33.33 | 22.22 | 34.78 |
| Umab_LC | h9C12 Q97A_HC | 32.54 | 9.09 | 34.78 |
| Umab_LC | h9C12_LC | 57.94 | 33.33 | 60.2 |
| Brimab_HC | Brimab_LC | 31.97 | 15.38 | 33.94 |
| Brimab_HC | h9C12 WT_HC | 50.83 | 25 | 54.13 |
| Brimab_HC | h9C12 Q97A_HC | 51.26 | 18.18 | 54.63 |
| Brimab_HC | h9C12_LC | 27.05 | 11.11 | 28.32 |
| Brimab_LC | h9C12 WT_HC | 31.15 | 8.33 | 33.64 |
| Brimab_LC | h9C12 Q97A_HC | 31.15 | 8.33 | 33.64 |
| Brimab_LC | h9C12_LC | 45.61 | 25 | 47.06 |
| h9C12-WT_HC | h9C12 Q97A_HC | 99.14 | 90.91 | 100 |
| h9C12-WT_HC | h9C12_LC | 25.86 | 30 | 25.71 |
| h9C12-Q97A_HC | h9C12_LC | 25.86 | 30 | 25.71 |

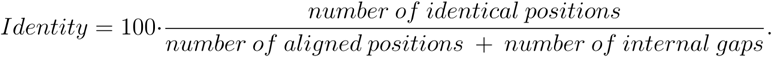

**Supplementary Table 4:**
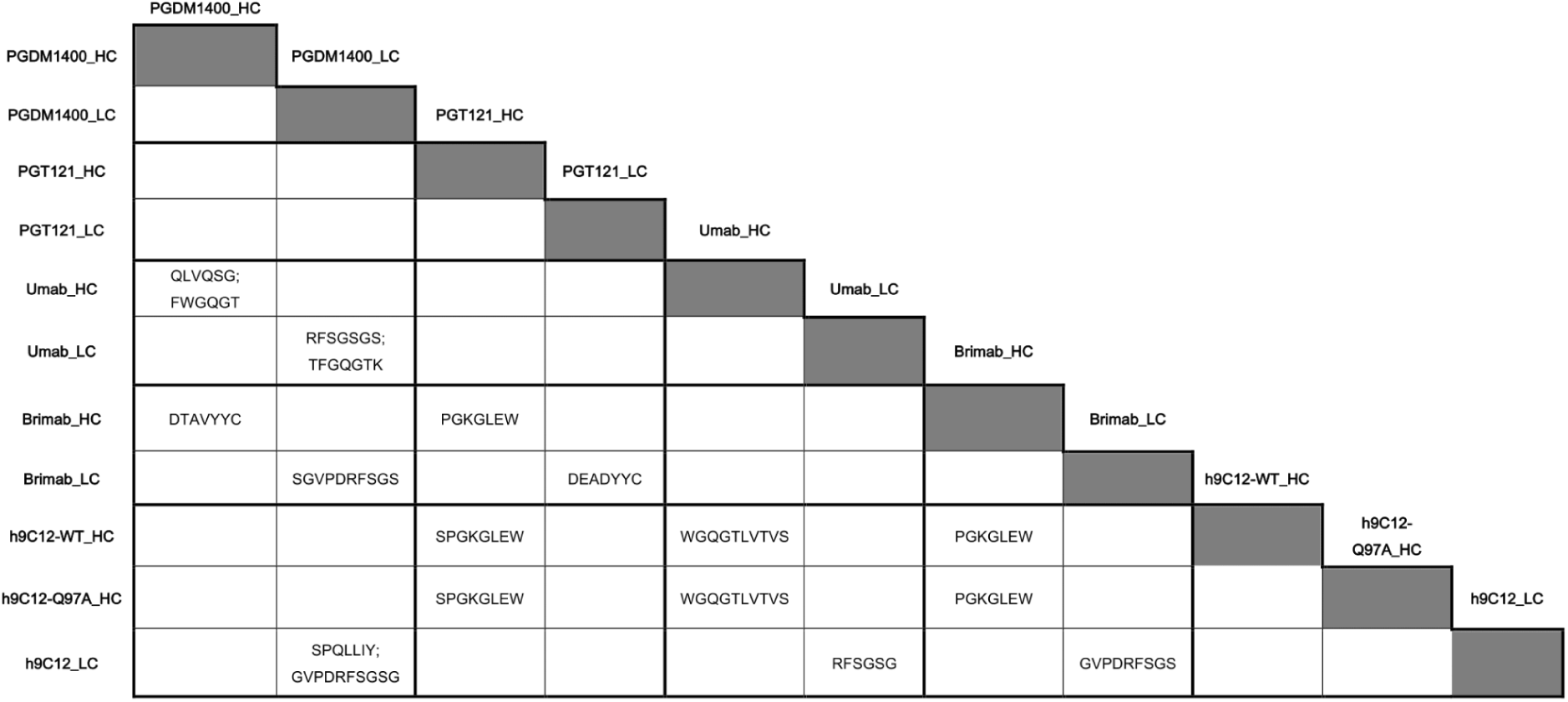
The longest common shared sub-sequences in the V(D)J region between pairs of monoclonal antibodies. Common sub-sequences are considered if the sequence is >5 aa. The sequences of h9C12 WT_HC and h9C12 Q97A_HC are not compared due to having only 1 AA difference between them (Q97A). All the shared sub-sequences are not within the CDR3 region.

**Supplementary Table 5:** Experimental consistency across the four experimental replicates (submissions). Experimental variables were kept as constant as possible across replicates, however, some parameters were changed across runs. These include protease supplier and 96-well plate well volume. We verified that the plate well volume did not have an impact on the number of mAb-related peptides discovered (Supplementary Figure 3).

| Experimental replicate | Protease treatment (in run order) | Manufacturer | Well volume | Volume of protease | Evosep column | Instrument cleaning |
| --- | --- | --- | --- | --- | --- | --- |
| 1 | Tryp | promega | 500 µl | 2 µl of 0.25 µg/µl |  |  |
| 1 | Ct | promega | 500 µl | 2 µl of 0.25 µg/µl |  |  |
| 1 | AspN | promega | 500 µl | 10 µl 0.05 µg/µl |  |  |
| 1 | Ct+Tryp | promega | 500 µl | 2 µl of 0.25 µg/µl + 2 µl of 0.25 µg/µl | changed to new column before Ct+Tryp |  |
| 2 | Tryp | promega | 500 µl | 2 µl of 0.25 µg/µl |  |  |
| 2 | Ct | promega | 500 µl | 2 µl of 0.25 µg/µl |  |  |
| 2 | AspN | promega | 500 µl | 5 µl 0.1 µg/µl | changed to new column before AspN |  |
| 2 | Ct+Tryp | promega | 500 µl | 2 µl of 0.25 µg/µl + 2 µl of 0.25 µg/µl |  |  |
| 3 | Tryp | promega | 1 ml | 2 µl of 0.25 µg/µl |  |  |
| 3 | Ct | thermo | 1 ml | 2 µl of 0.25 µg/µl |  |  |
| 3 | AspN | promega | 1 ml | 5 µl 0.1 µg/µl | changed to new column before AspN |  |
| 3 | Ct+Tryp | thermo/promega | 1 ml | 2 µl of 0.25 µg/µl + 2 µl of 0.25 µg/µl |  |  |
| 4 | Tryp | promega | 500 µl | 2 µl of 0.25 µg/µl |  |  |
| 4 | Ct | thermo | 500 µl | 2 µl of 0.25 µg/µl | changed to new column before Ct | QExactive HF was cleaned before Ct |
| 4 | AspN | promega | 500 µl | 5 µl 0.1 µg/µl |  |  |
| 4 | Ct+Tryp | thermo/promega | 500 µl | 2 µl of 0.25 µg/µl + 2 µl of 0.25 µg/µl |  |  |

**Supplementary Table 6:**
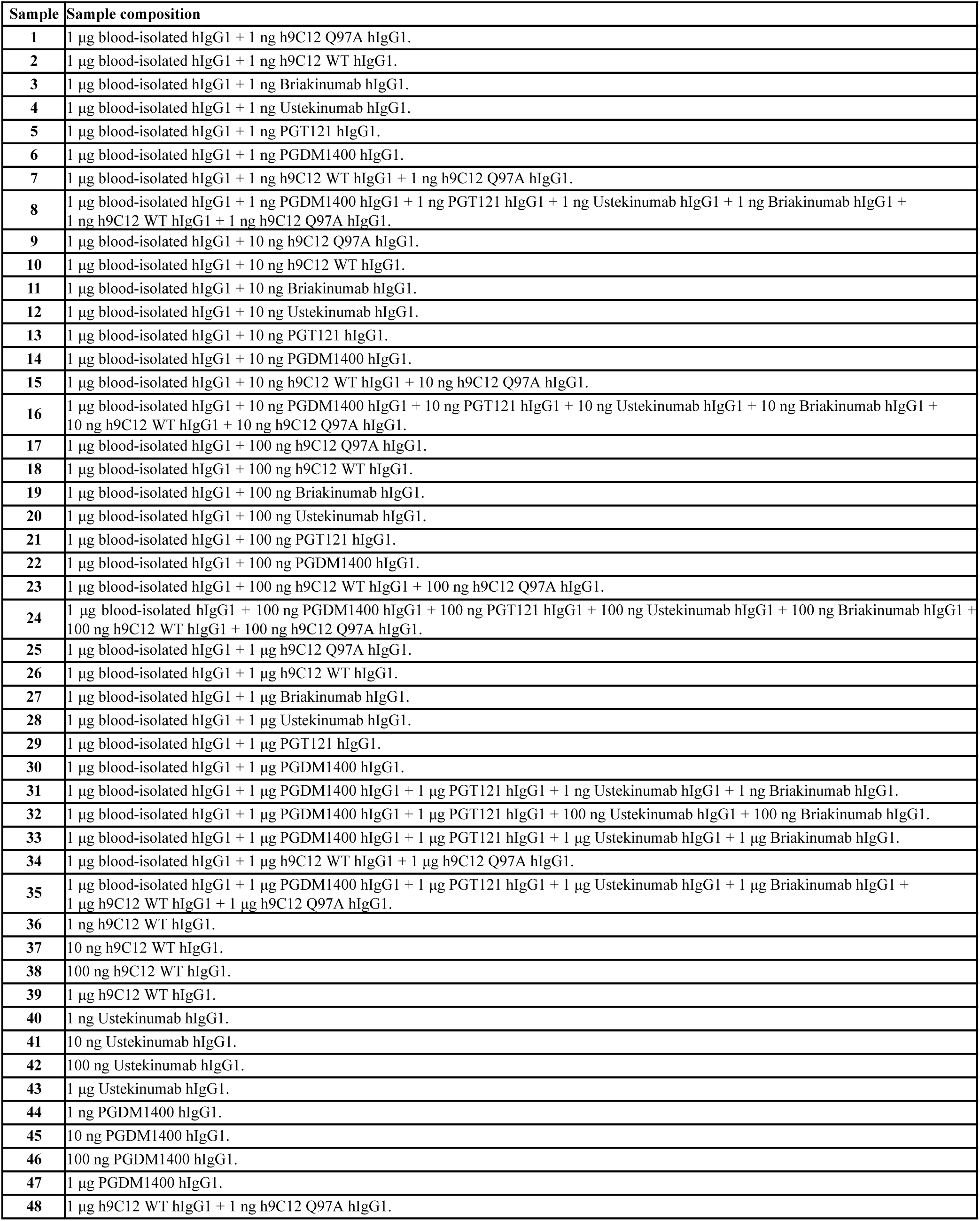
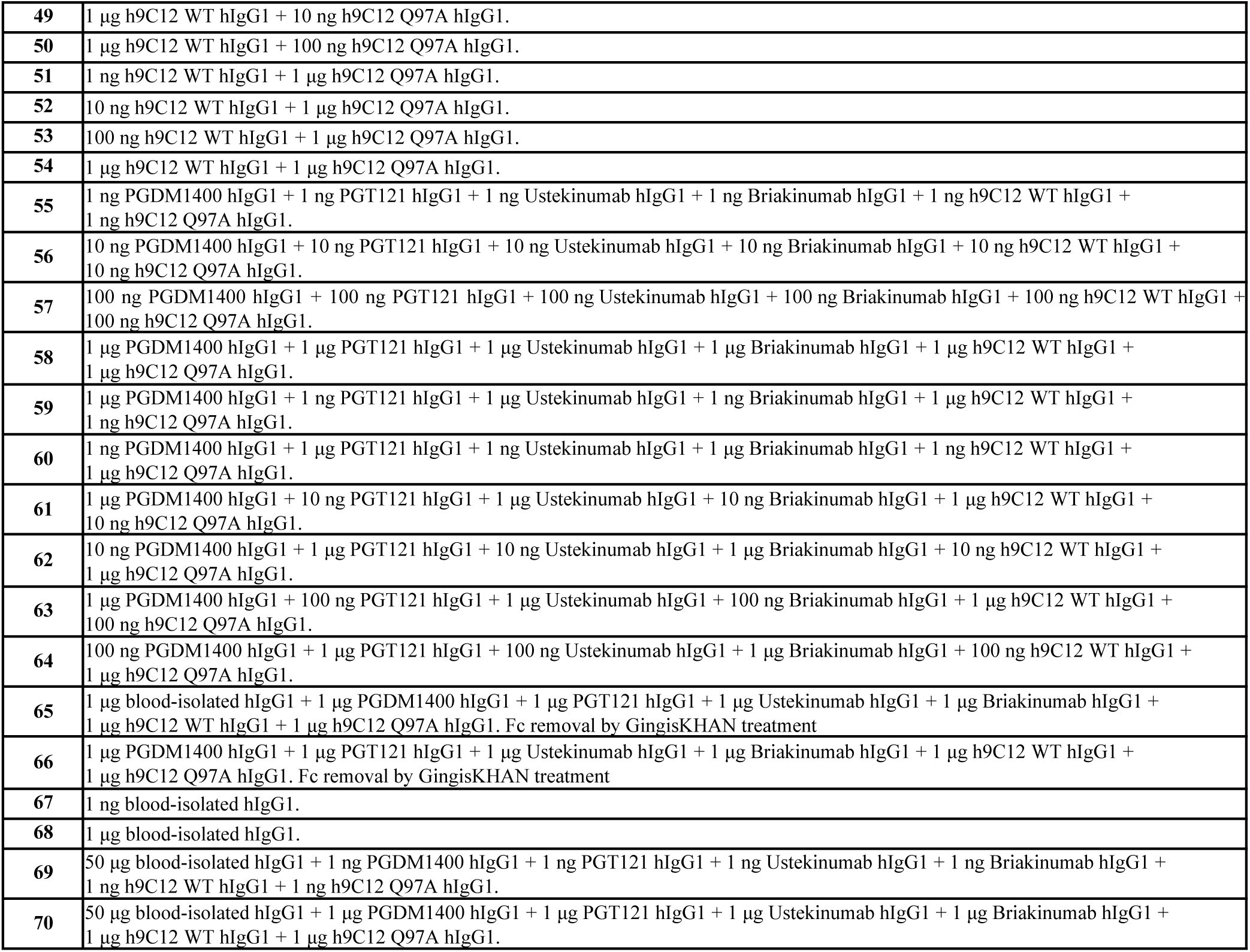
Sample description.

**Supplementary Figure 1:**
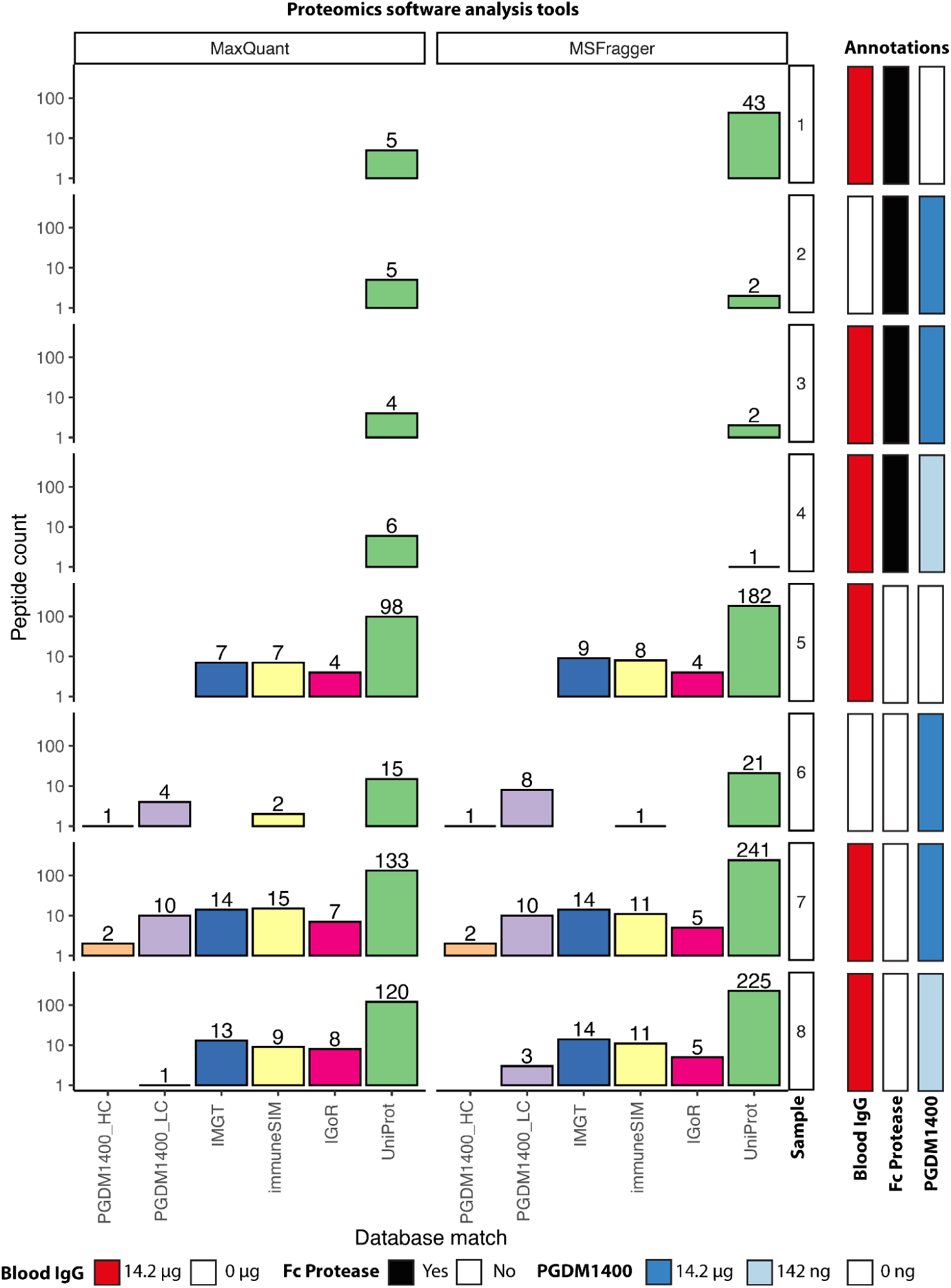
Pilot experiment: Fc protease treatment does not increase the detection of mAb-related peptides from the samples. We analyzed how GingisKHAN (a protease digesting human IgG1at a single site above the hinge region) treatment before enzymatic digestion influenced mAb-related peptide detection. To this end, we analyzed mAb-related peptide detection for blood IgG (14.2 μg) either alone or mixed with the monoclonal antibody PGDM1400 (142 ng–14.2 μg) with and without Fc protease treatment. We found that overall, the number of mAb-related peptides (peptides that aligned to the variable region of a reference Ab) identified was higher for samples without Fc protease treatment (samples 5–8) than for samples with Fc protease treatment (samples 1–4). Chymotrypsin+Trypsin were used to digest antibodies into peptides for this experiment. This pilot experiment was performed prior to the main experiment shown in Figure 2. **Background for this experiment:** Although previous reports used Fc cleavage to increase signal-to-noise-ratio ^29,31,59^, our efforts to analyze mAb-related peptides by utilizing GingisKHAN protease for Fc cleavage yielded unsatisfactory results (Supplementary Figure 1), which may be attributed to multiple factors. Notably, antibody concentrations employed in our study spanned 10 ng/mL to 10 µg/mL. Given that median concentration of all IgGs in human adults are around 10900 µg/mL ^124^, and that the antibody repertoire follows a power law distribution ^125,126^, our spike-in mAbs inputs would be more representative of individual clonotypes present in the serum obtained from a blood draw. However, this range of concentration is substantially below the manufacturer’s recommended lower threshold of 0.5 mg/mL per reaction column ^127^. This concentration is also markedly less than the quantities reported in previous studies, which typically range from 1 mg/mL to 5 mg/mL ^128,129^. At the low concentrations of Ab considered in our study, catalysis efficiency of the enzyme may be compromised, requiring a longer incubation time, which can in turn may have led to enzyme degradation. Therefore, our data suggest that while Fc cleavage can enhance peptide detection in the V(D)J region for high-concentration single mAb samples, its efficacy is diminished in diluted and polyclonal antibody samples.

**Supplementary Figure 2:**
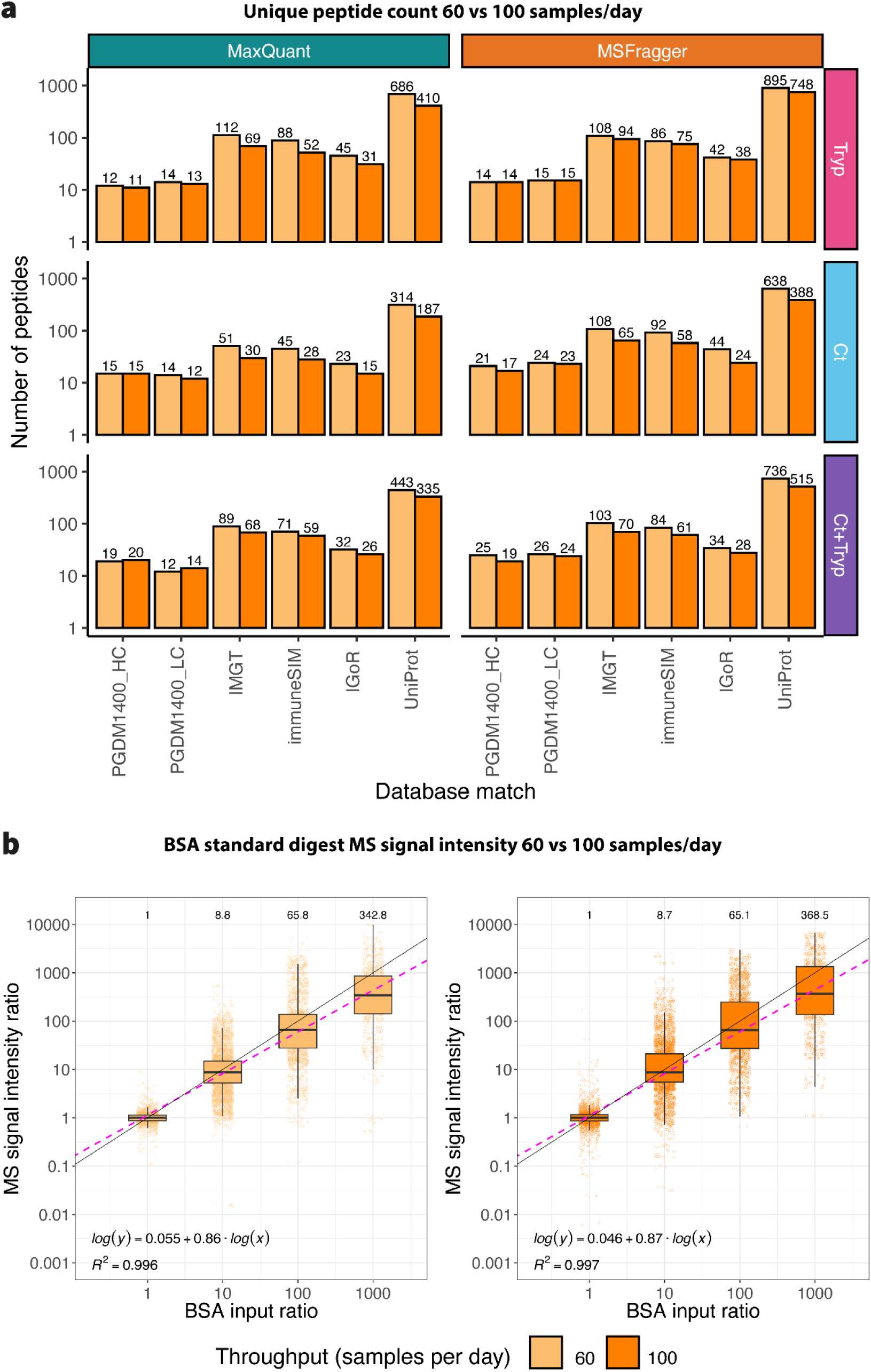
Pilot experiment: comparing two different Evosep methods (60 samples per day and 100 samples per day) shows that using a higher throughput LC-method does not compromise the quality of peptide detection and qualification. (a) We analyzed 14.2 μg of the monoclonal antibody PGDM1400 with three different protease combinations (Tryp, Ct, Ct+Tryp) using two different Evosep methods (60 samples per day; 21 min LC-separation gradient and 100 samples per day; 11.5 min LC-separation gradient) and did not observe a major difference in Ab-specific peptide counts on the HC and LC of PGDM1400. (b) We analyzed BSA protein digest standards at 5 pmol, 500 fmol, 50 fmol, and 5 fmol. The values of MS1 signal intensity ratios and mAb input ratios were log10-transformed, and the median MS1 signal intensity ratios were used for linear regression, weighted by the number of data points in each mAb input ratio. The dashed purple line represents the linear model that predicts the MS1 signal intensity ratios from the mAb input ratio, while the black solid represents the expected linear relationship of the two variables. The median MS1 signal intensity ratio for each mAb input ratio is shown above each boxplot, the predictor function is shown as log(y) = intercept + slope*log(x), and the goodness-of-fit is denoted by the coefficient of determination (R2). This pilot experiment was performed prior to the main experiment shown in Figure 2.

**Supplementary Figure 3:**
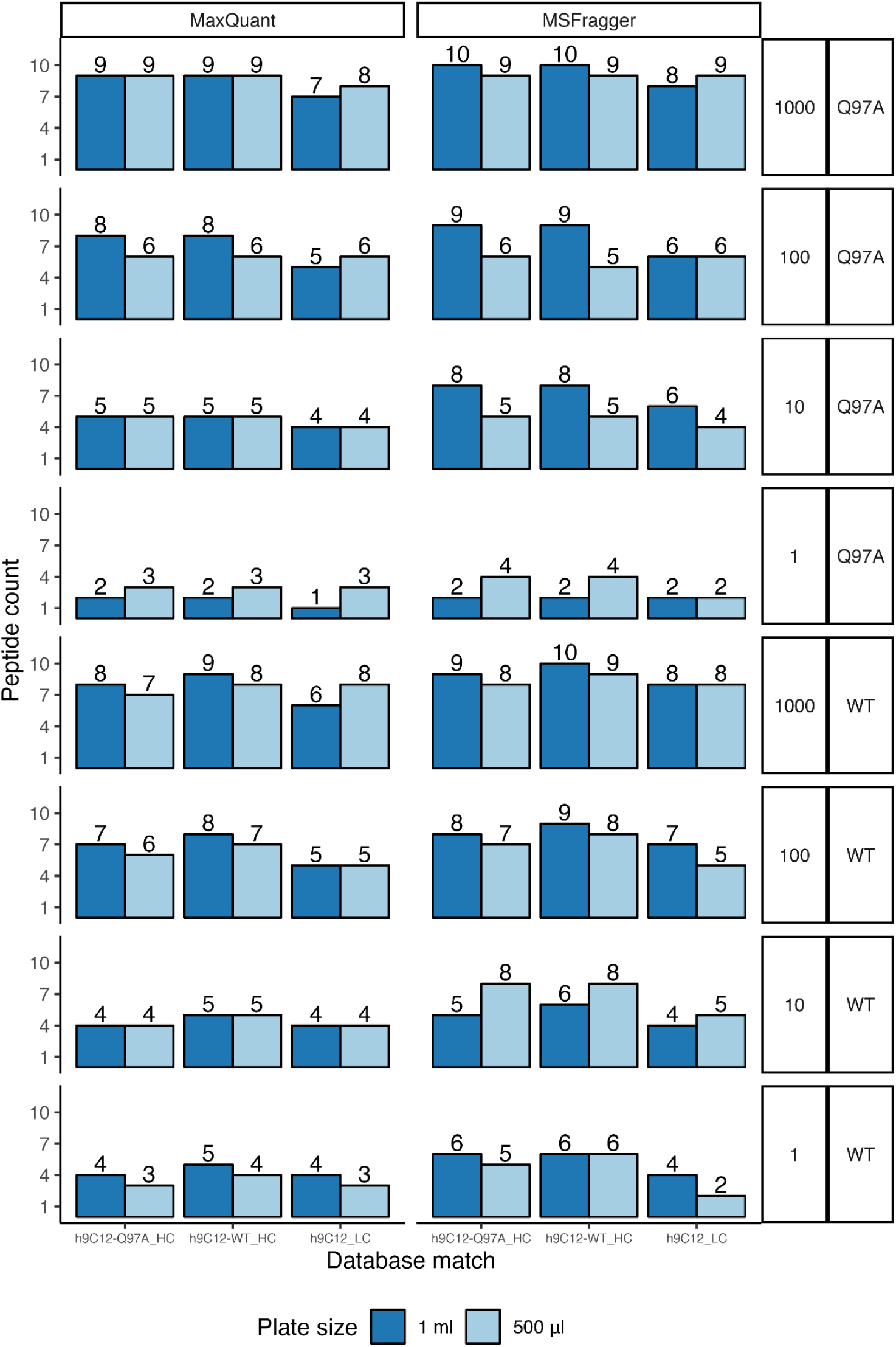
Pilot experiment: the 96-well plate well volume has no effect on peptide detection. We tested the influence of plate well volume (500 μl and 1 ml) on mAb-related peptide detection, due to the fact that we used different volume plates between the experimental replicates (Supplementary Table 5). Here, we used the protease trypsin and the two antibody variants h9C12 WT and h9C12 Q97A. We found that neither mAb-related peptides did not substantially differ by plate well volume. This pilot experiment was performed prior to the main experiment shown in Figure 2.

**Supplementary Figure 4:**
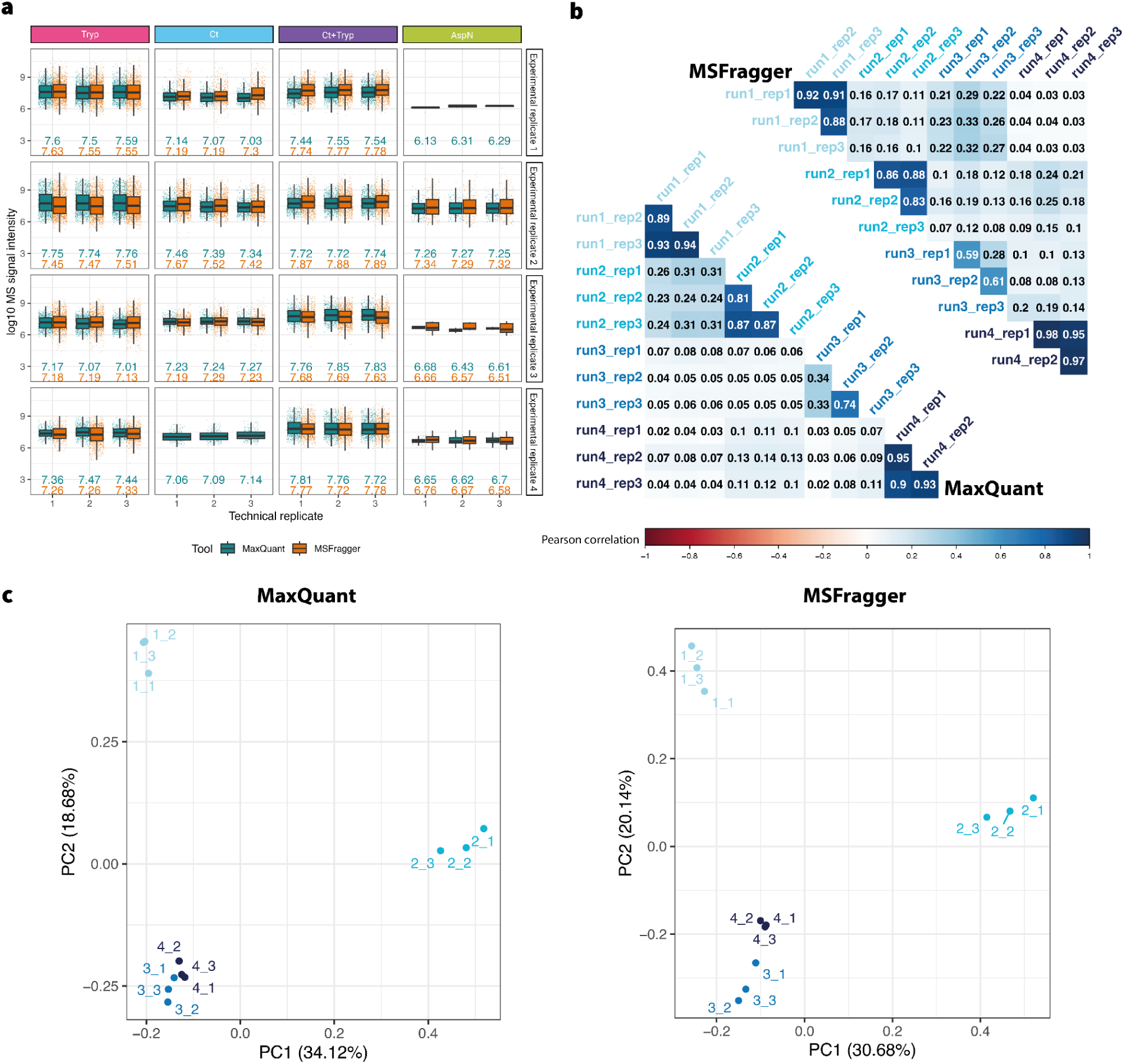
Technical replicates show high replicability in terms of signal intensity. Replicability was measured by comparing MS1 signal intensity values unique for each combination of: peptide sequence, sample number, and protease treatment used, with values compared across experimental and technical replicates. Peptides not detected in certain replicates are considered to have intensity of 0. **(a)** Distribution of the log10 transformed MS1 signal intensities, colored by proteomics software used. Numbers underneath each boxplot are the median values of the log10 transformed MS1 signal intensities. **(b)** Pearson correlation of the MS1 signal intensities, the number in each cell is the Pearson’s correlation value. Labels run*x*_rep*y* represent the experimental replicate (*x*) and technical replicate (*y*) number. **(c)** PCA of the MS1 signal intensities from MaxQuant and MSFragger, with each point (colored by experimental replicate) being one technical replicate in an experimental replicate, labels *x*_*y* represent the experimental replicate (*x*) and technical replicate (*y*) number, respectively. Relates to Figure 1c.

**Supplementary Figure 5:**
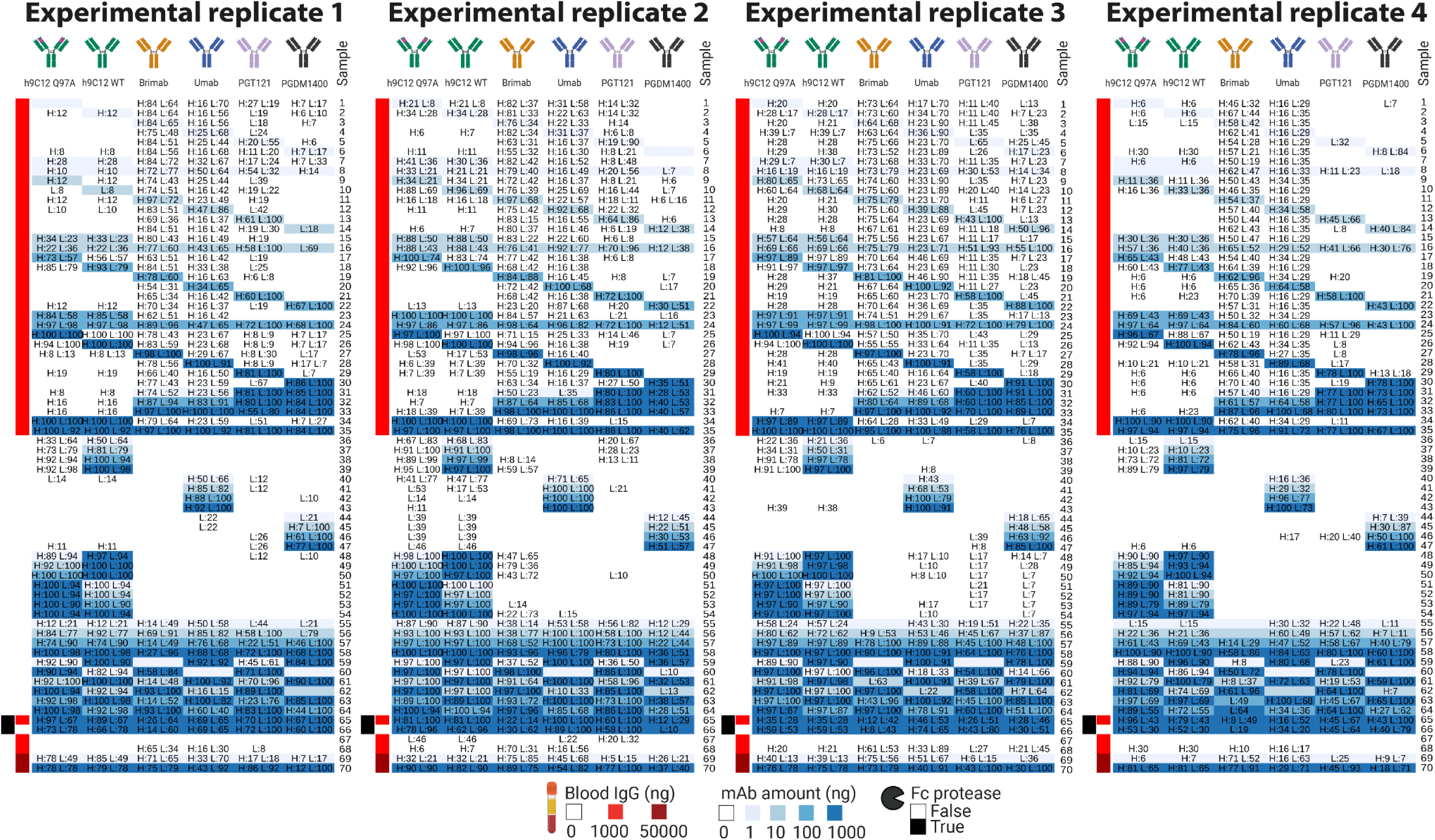
V(D)J sequence coverage in each sample by experimental replicate. Peptides merged from all bioinformatic software analysis tools after preprocessing were aligned with the mAb V(D)J sequences and coverage (displayed as percentage, calculated according to Supplementary Figure 12) were calculated for each sample. H:xx & L:yy are used to denote sequence coverage on the HC and LC, respectively. Pooled V(D)J coverage from all experimental replicates is shown in Supplementary Figure 11. Relates to Figure 2b.

**Supplementary Figure 6:**
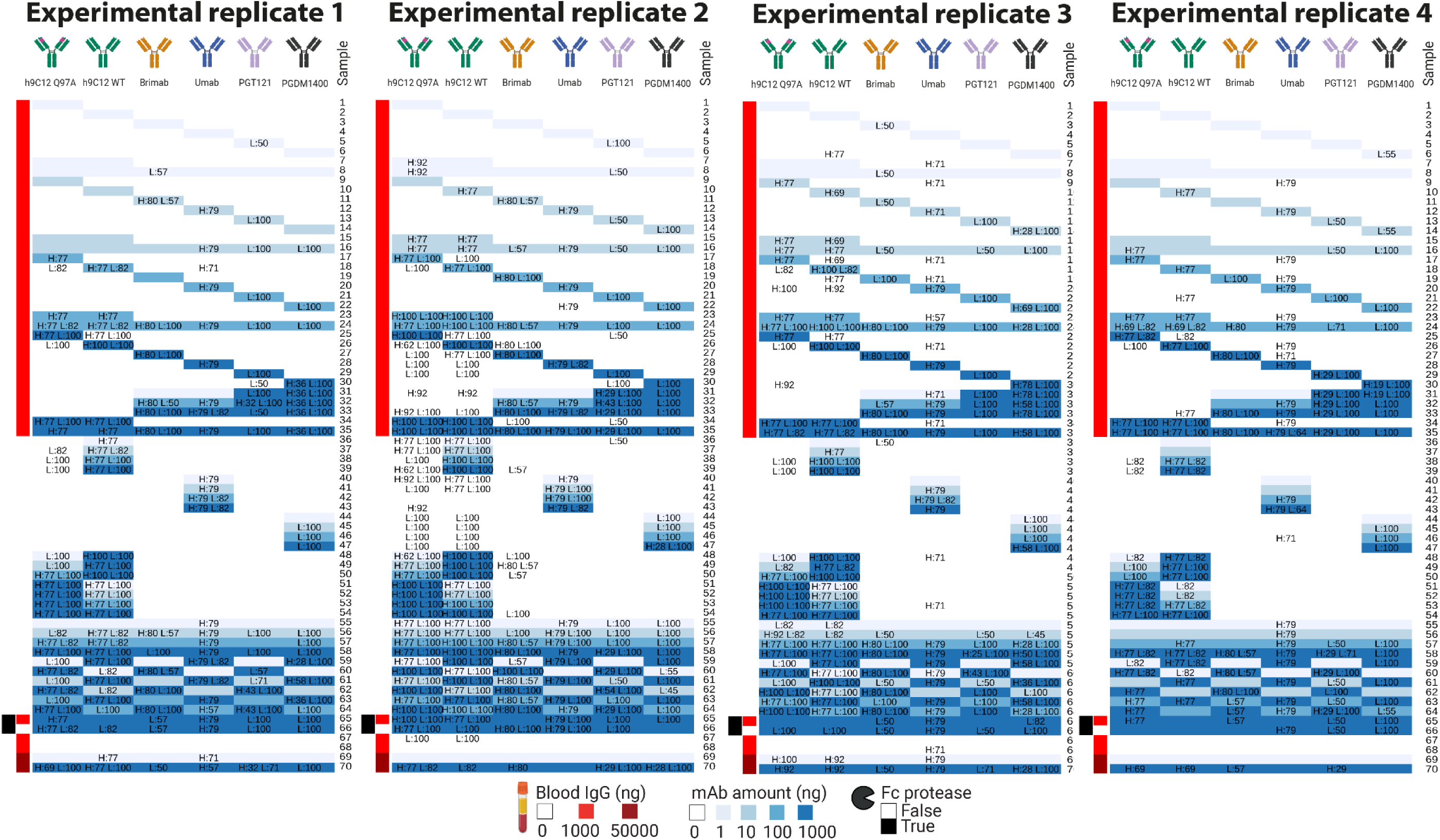
CDR3 sequence coverage in each sample by experimental replicate. Peptides merged from all bioinformatic software analysis tools after preprocessing were aligned with the mAb CDRH3/CDRL3 sequences and coverage (displayed as percentage, calculated according to Supplementary Figure 12) were calculated for each sample. H:xx & L:yy are used to denote sequence coverage on the HC and LC, respectively. All unique peptides in all experimental replicates were used in CDR3 sequence coverage shown in Figure 2b.

**Supplementary Figure 7:**
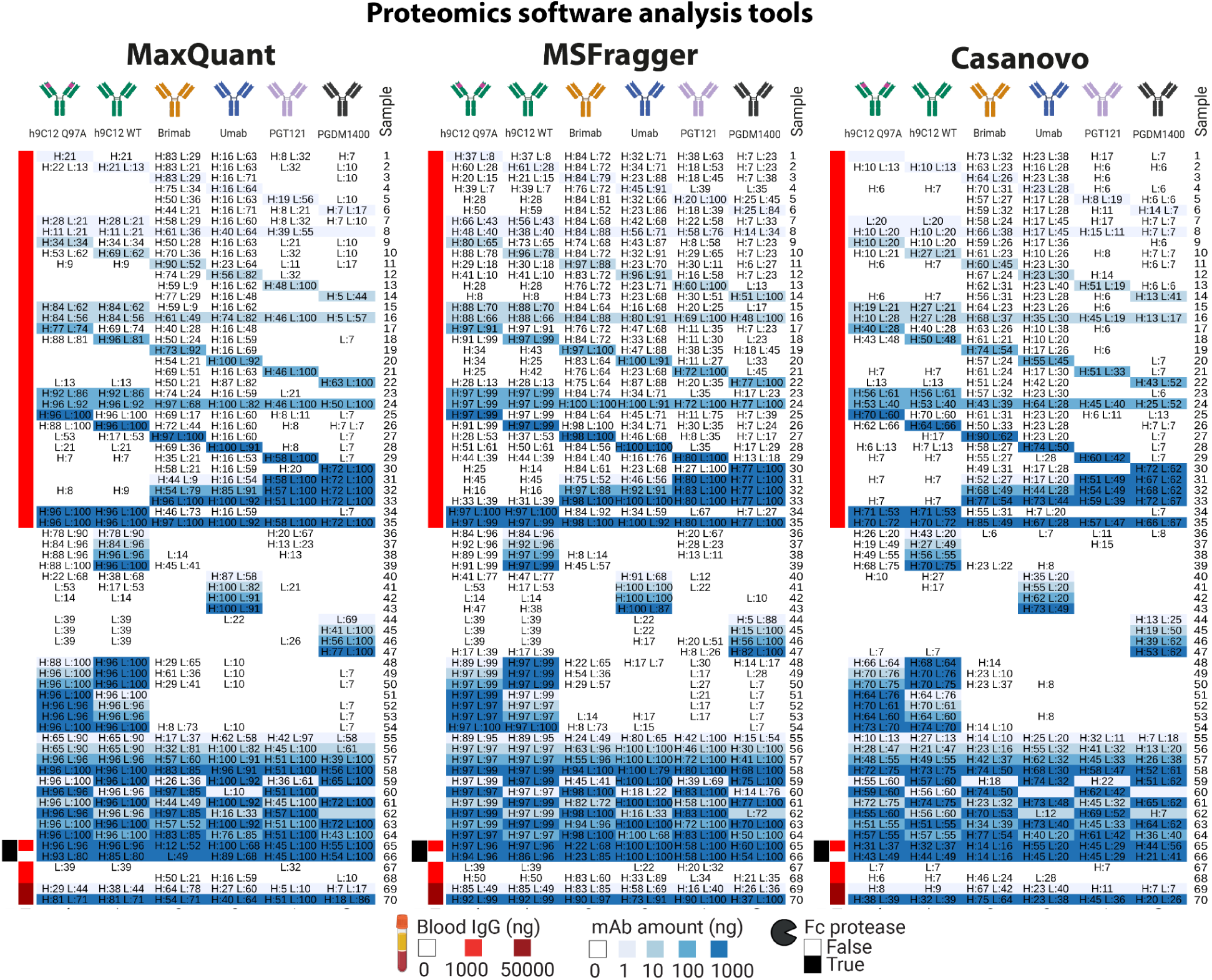
V(D)J sequence coverage in each sample by proteomics software analysis tool. Mass spectrometry raw data was used to determine peptide sequences by either MaxQuant, MSFragger, or inferred *de novo* by Casanovo. Peptides from all experimental replicates after preprocessing were aligned with the ground truth mAb V(D)J sequences and coverage (displayed as percentage, calculated according to Supplementary Figure 12) were calculated for each sample. H:xx & L:yy are used to denote sequence coverage on the HC and LC, respectively. Pooled V(D)J coverage is shown in Supplementary Figure 11 and pooled CDR3 coverage is shown in Figure 2b.

**Supplementary Figure 8:**
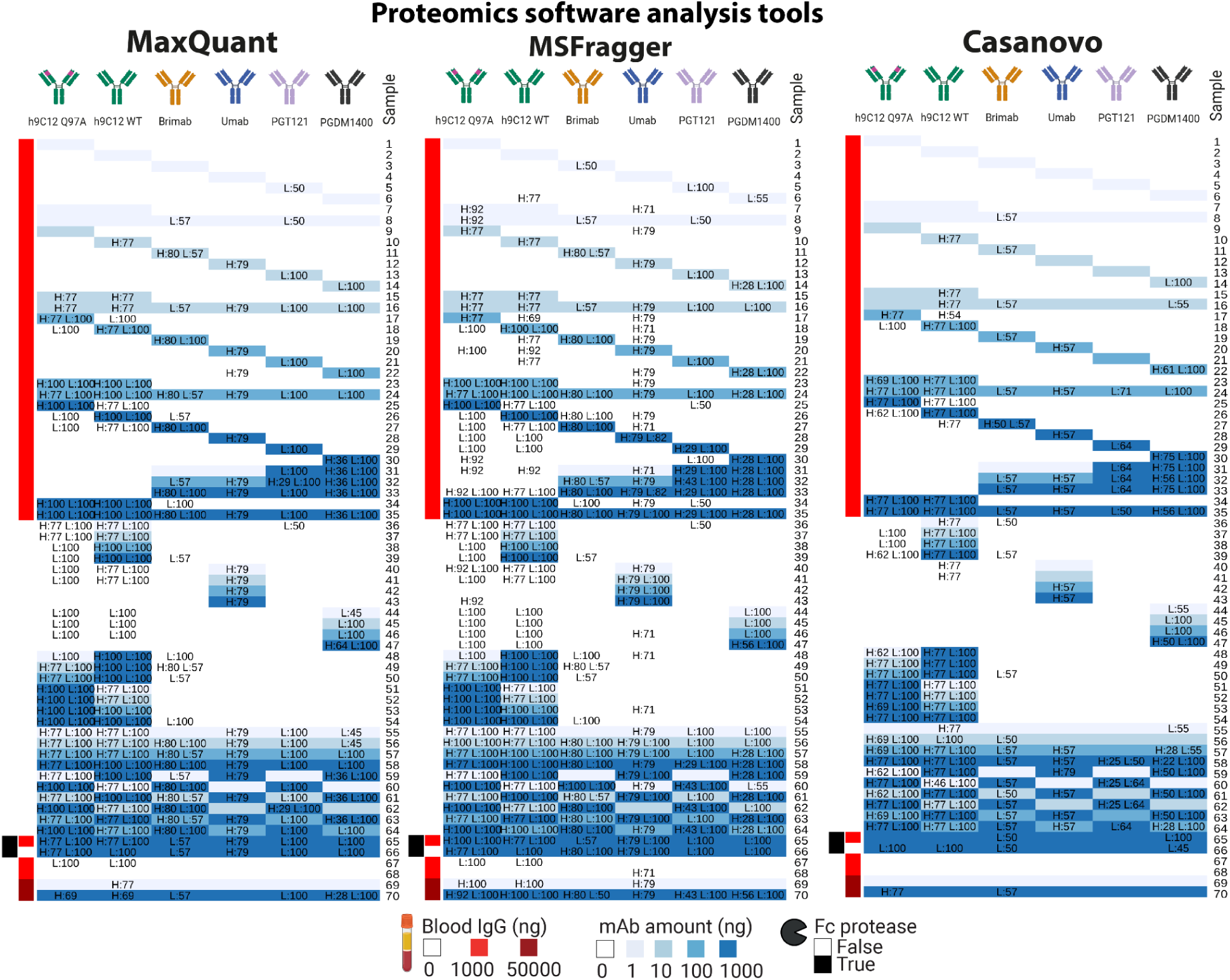
CDR3 sequence coverage in each sample by proteomics software analysis tool. Mass spectrometry raw data was used to determine peptide sequences by either MaxQuant, MSFragger, or inferred *de novo* by Casanovo. Peptides from all experimental replicates after preprocessing were aligned with the mAb CDRH3/CDRL3 sequences and coverage (displayed as percentage, calculated according to Supplementary Figure 12) were calculated for each sample. H:xx & L:yy are used to denote sequence coverage on the HC and LC, respectively. All unique peptides in all proteomics software analysis tools were used in CDR3 sequence coverage shown in Figure 2b.

**Supplementary Figure 9:**
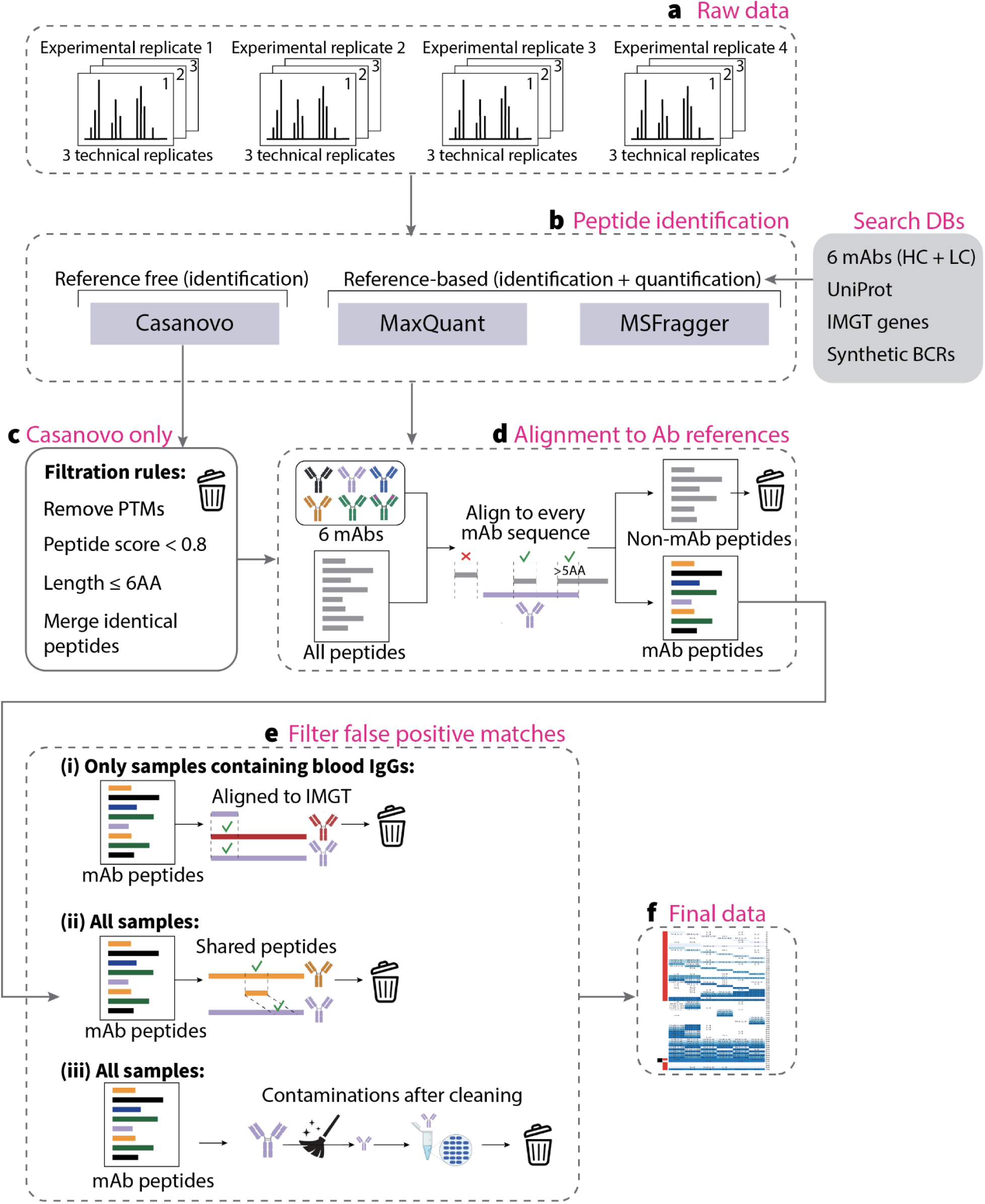
Data preprocessing pipeline. **(a)** Raw data merged from four experimental replicates and three technical replicates per each experimental replicate **(b)** was analyzed using three proteomics tools — Casanovo for *de novo* peptide identification^45^ and two PSM tools for peptide identification and quantification — MaxQuant ^75^ and MSFragger ^76^. The following search databases were used for the PSM tools (MaxQuant and MSFragger): sequences of the HC and the LCs of the 6 monoclonal antibodies, UniProt (“UniProt: The Universal Protein Knowledgebase in 2023” 2023), IMGT (database of V and J genes for IGH, IGK, and IGL chains, downloaded July 2020), and a set of synthetic IGH, IGK, and IGL (30000 in total) human sequences generated with immuneSIM ^117^. **(c)** We further preprocessed the Casanovo-predicted peptides by removing PTMs, as well as eliminating peptides with a Casanovo search engine score lower than 0.8 or a length less than 6 amino acids. Subsequently, we merged identical peptides within each sample and technical replicate. We additionally preprocessed Casanovo-predicted peptides by removing PTMs and removing peptides with a Casanovo search engine score lower than 0.8 or length less than 6AA. After that, we merged identical peptides within every sample and every technical replicate and assigned the maximum search engine score to the final peptide among the corresponding merged peptides. **(d)** All identified peptides were aligned to the six monoclonal antibody HC and LC variable region sequences. A peptide was considered as an mAb peptide, if it can be matched (aligned with 100% identity) or overlapped (with at least 5AA) to at least one of the mAb HC or LC references. Otherwise, a peptide was considered as non-mAb, and thus, removed. **(e)** After the alignment step, we processed the false positive peptides, i.e., peptides that map to a mAb that is not present in the sample by design. Among false positive peptides, we filtered the mAb peptides if it: (i) belongs to a sample containing blood-derived IgGs and align to the IMGT database or (ii) align to more than one mAb sequence, where one mAb is present in a sample and the other mAb is not present. The first criteria allowed removing peptides that are shared between the six mAbs and blood IgGs. For instance, if a blood IgG is made of IGHV1-2, then some peptides from its V region may be also aligned to the PGDM1400 HC, because it is also composed of IGHV1-2. The second criteria allowed to filter the peptides shared between mAb sequences (Supplementary Table 4). For example, if we detect a peptide GVPDRFSGSG in the sample 6 (1 ng of PGD1400 with blood IgGs), then it will map both to the PGD1400 LC and h9C12 LC, and thus h9C12 will be a false positive match in this case. As a final step (iii), we removed the false positive peptides that are likely biological contamination because they were detected in the effluent of the previous cleaning cycle. To this end, we removed those peptides that corresponded to mAbs which should have been absent in a sample by experimental design if this peptide was also detected in the previous blank sample. **(f)** The preprocessed data was used for all main figures (Figure 2, Figure 3, Figure 4, Figure 5, Figure 6), however in Figure 2c-d, Figure 3, Figure 4, Figure 5, Figure 6 we only present results obtained from true positive peptides to exclude potential bias from FP peptides. Unless specified, all data visualized in the manuscript are from merged technical replicates (concordance between technical replicates is shown in Supplementary Figure 4). Peptides filtered out during preprocessing are shown in Supplementary Figure 10.

**Supplementary Figure 10:**
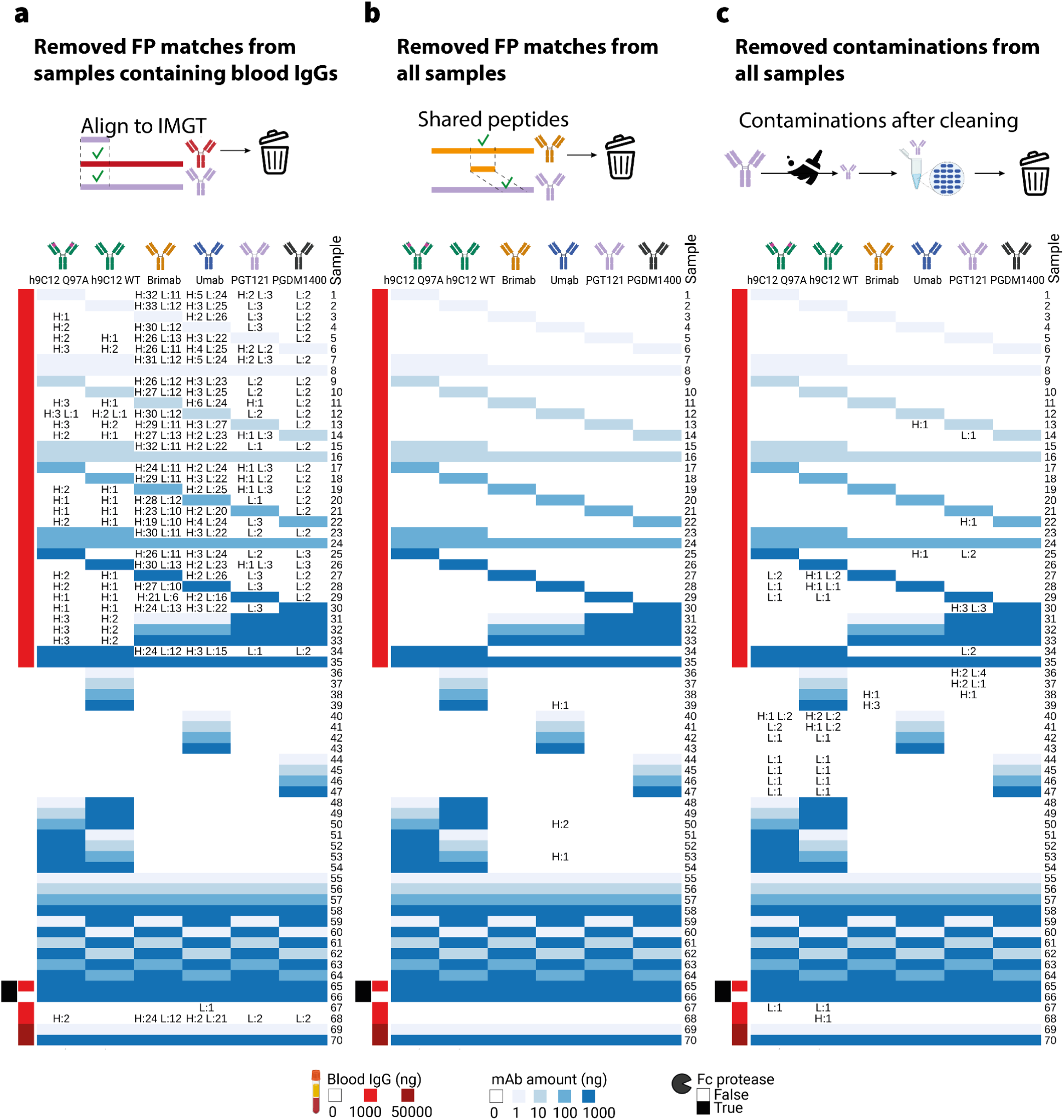
False positive peptides removed during data preprocessing. The dataset comprising peptide sequences from all experimental and technical replicates was subjected to a progressive filtering process designed to eliminate false positives—specifically, peptides matching monoclonal antibody (mAb) sequences absent in the respective samples (described in Supplementary Figure 9). This clean-up procedure involved sequential stages: **(a)** elimination of peptides corresponding to sequences registered in the IMGT V(D)J database, in order to exclude generic matches, **(b)** removal of peptides mapped to regions shared by multiple mAbs, and **(c)** exclusion of peptides detected in the intermediate MS purification runs carried out between samples. Only peptides that passed through all filtering steps were displayed in Figure 2.

**Supplementary Figure 11:**
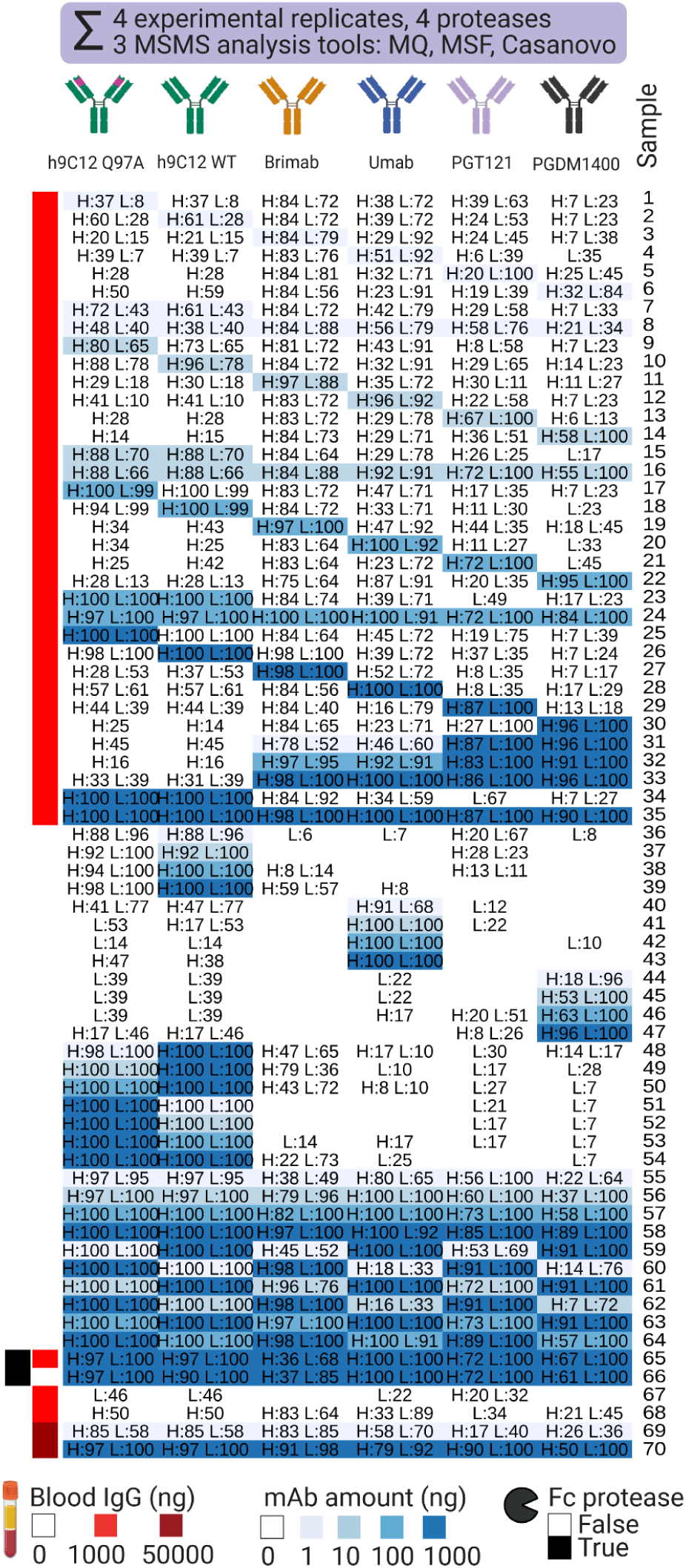
Pooled V(D)J coverage of all proteases, experimental replicates, and proteomics software analysis tools. Peptides that passed through all filtering steps were pooled by sample and V(D)J sequence coverage was calculated for each sample (displayed as percentage, calculated according to Supplementary Figure 12). H:xx & L:yy are used to denote sequence coverage on the HC and LC, respectively. Pooled CDR3 coverage is shown in Figure 2b.

**Supplementary Figure 12:**
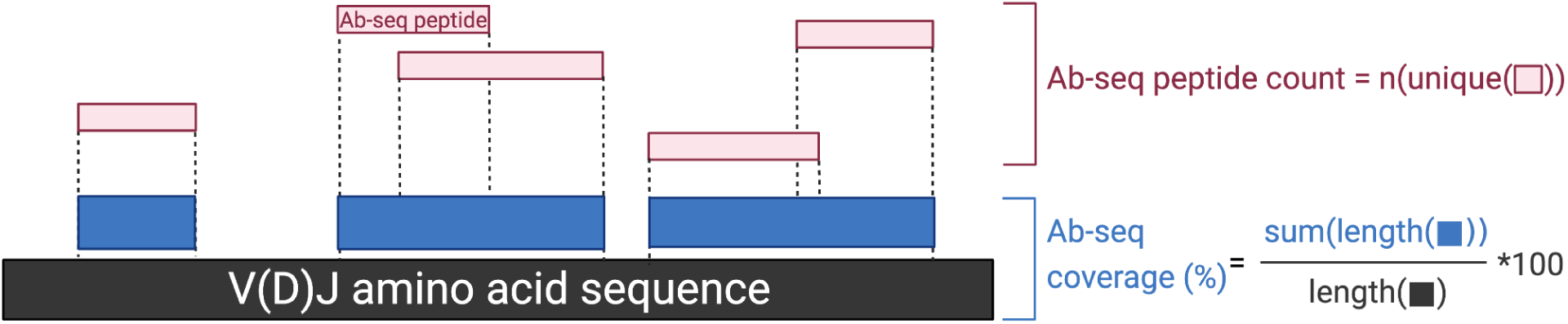
Difference between Ab-seq coverage and Ab-seq peptide count in terms of recovery of the antibody V(D)J sequence and rationale for using Ab-seq coverage as the main Ab-seq performance metric. Ab-seq peptide count (n(unique(peptide))) is the number of unique Ab-seq peptides, which can be mapped to a given V(D)J sequence. Ab-seq coverage is the percentage of covered amino acid positions of a given antibody V(D)J sequence, and can be calculated as the number of V(D)J sequence amino acids covered by Ab-seq peptides (sum(length(▪)) divided by the amino acid (number of amino acids) of the V(D)J sequence (length(▪)). We used Ab-seq coverage and not Ab-seq peptide count as a measure of Ab-seq performance because our focus in this work was on measuring the extent of the recovery of mAb V(D)J sequences. As visualized, Ab-seq peptides may overlap sequence-wise, and therefore, Ab-seq peptide count may not represent a faithful reflection of the extent of recovery of the V(D)J sequence. Consequently, an increasing Ab-seq peptide count may not lead to higher recovery of the antibody V(D)J sequence. Coverage metrics were utilized in Figure 2, Figure 3, Figure 4, Figure 6.

**Supplementary Figure 13:**
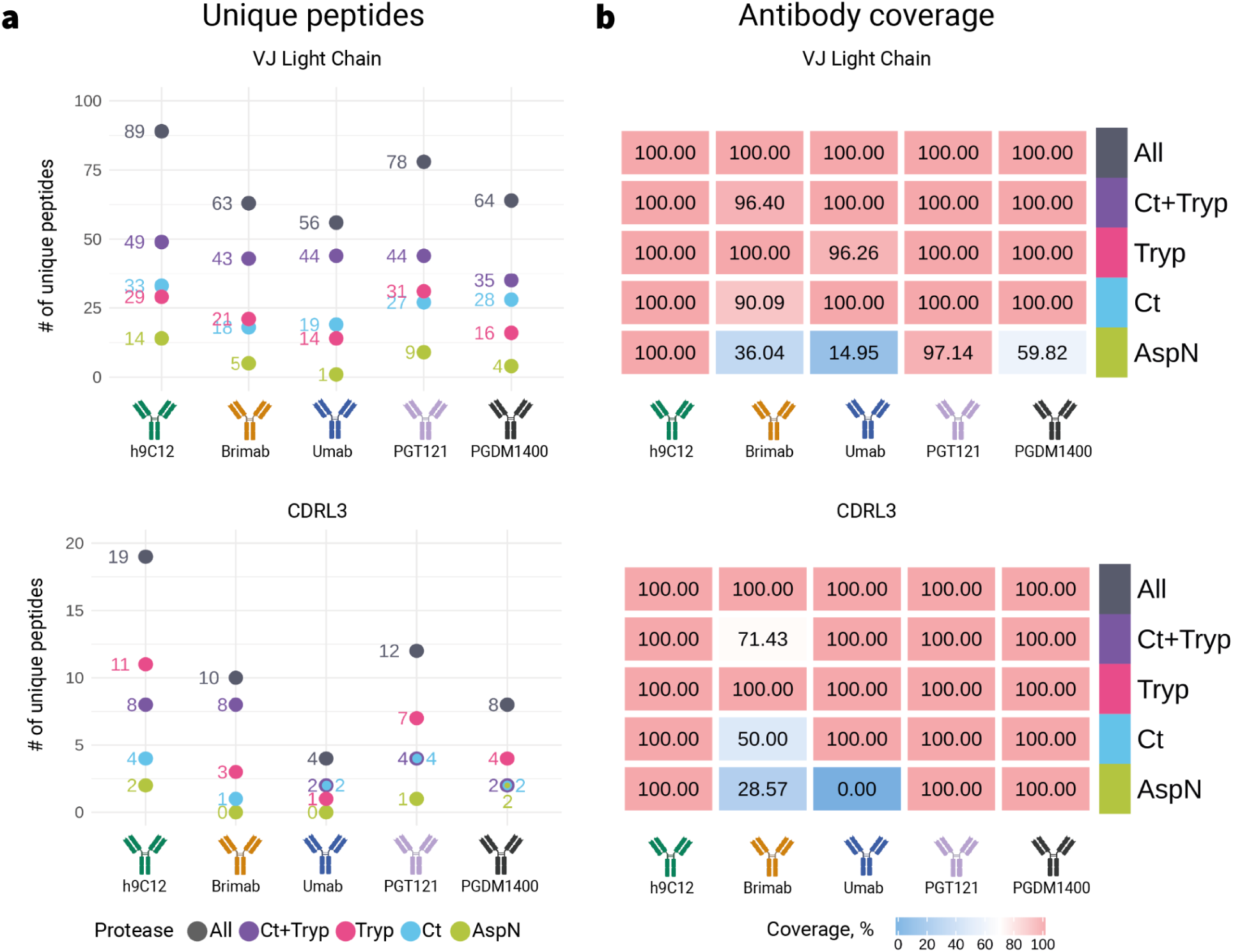
Peptide count and sequence coverage on the LC by protease. (a) Number of unique true positive peptides in the full-length VJ region and overlapping with the CDRL3 region of antibody LC. Peptide counts are pooled for all experimental and technical replicates. (b) Full-length LC VJ and CDRL3 coverage (the percentage of the reference Ab sequence detected by Ab-seq) of true positive peptides pooled from four experimental replicates achieved by every protease digestion. Peptide count and sequence coverage on the HC by protease was displayed in Figure 2c, d.

**Supplementary Figure 14:**
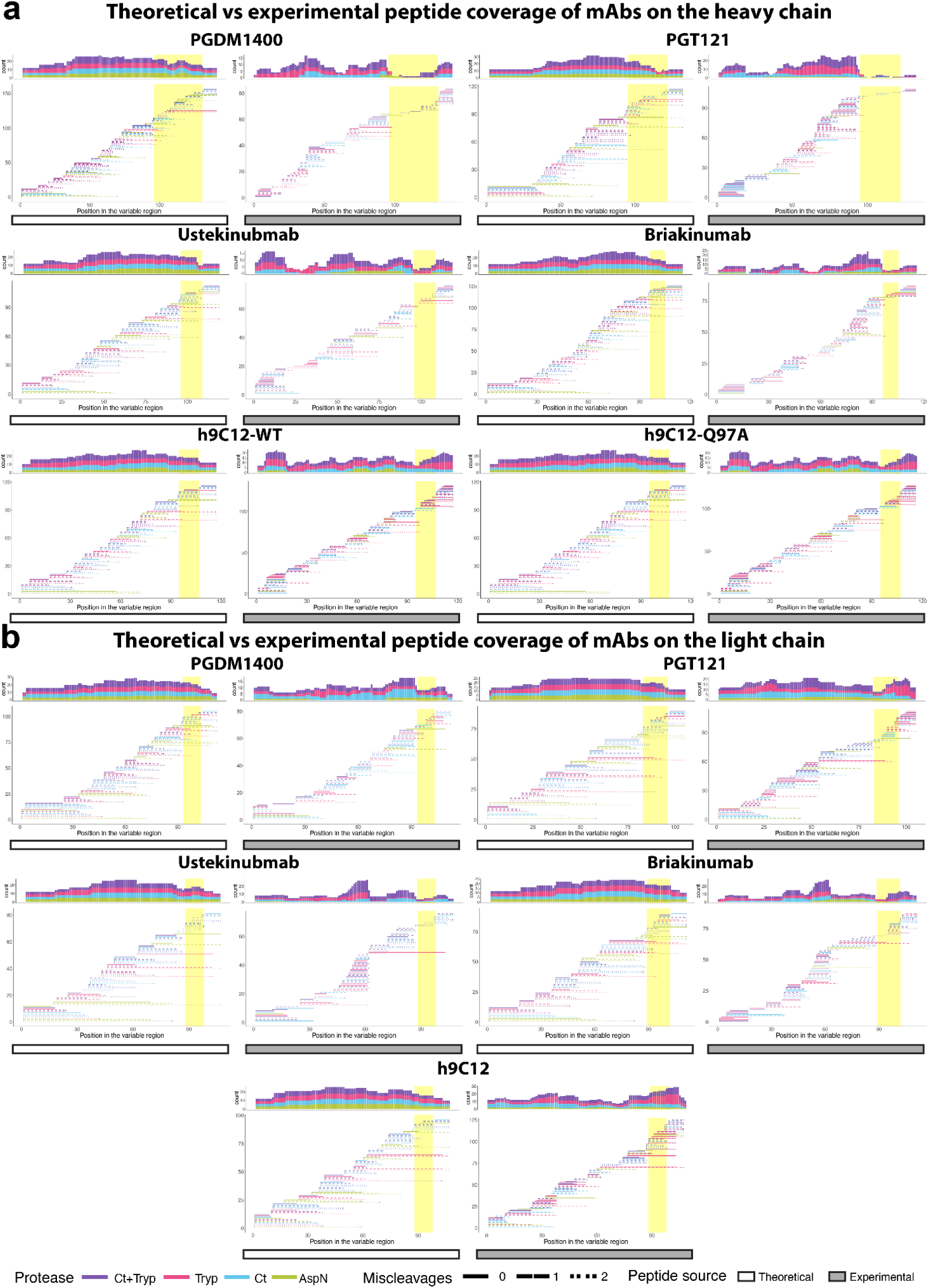
Theoretical and experimental peptide coverage of mAbs (V(D)J and CDR3) on the HC and LC by protease. The same theoretical peptides derived from in silico cleavage described in Supplementary Figure 16 were compared to the peptide output from mass spectrometry experiments. Resulting theoretical and experimental peptides (as lines colored by protease used and patterned by number of miscleavages) were then aligned to the V(D)J region of the mAbs’ sequences on the (a) HC and (b) LC (CDR3 region highlighted in yellow) to visualize the coverage of MS peptides in regard to the whole V(D)J region. Histograms above each coverage plot represent the absolute number of peptides covering each amino acid position in the respective V(D)J region. All preprocessed experimental peptides were shown as cdr3 coverage percentage in Figure 2b and VDJj coverage percentage in Supplementary Figure 11.

**Supplementary Figure 15:**
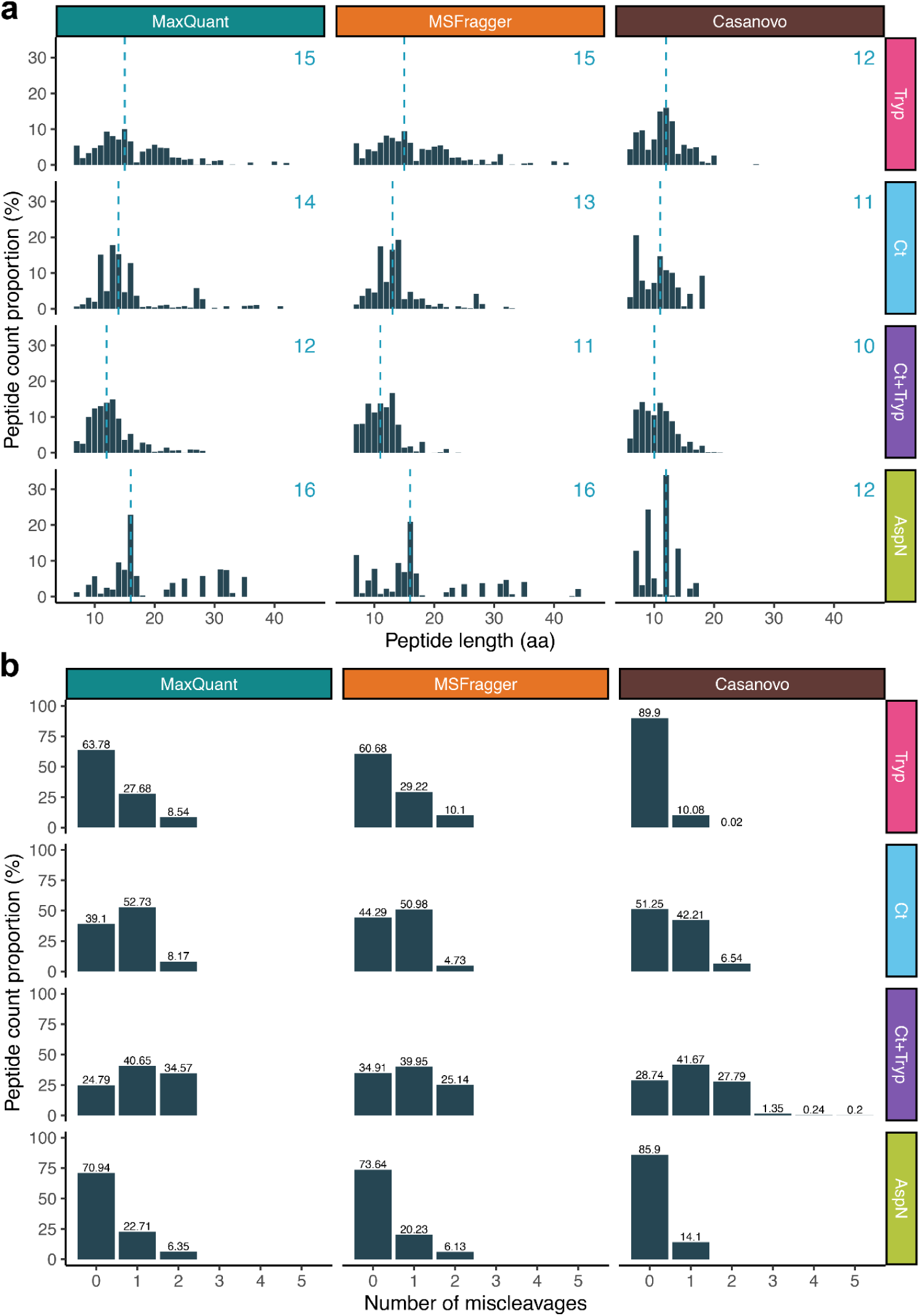
Peptide length and miscleavage distribution. Mass spectra from 1120 samples were processed by either MaxQuant, MSFragger, or Casanovo respectively and peptide sequences were determined. **(a)** Distribution of peptide length (aa) as percentage of all peptides. The median peptide length is shown per combination of software and protease in the upper right corner. **(b)** Number of miscleavages distribution as a percentage of all peptides. For MaxQuant and MSFragger, a maximum of two miscleavages were allowed for downstream analyses, while Casanovo did not have a limit of miscleavages allowed. After preprocessing, the experimental peptide sequences displayed protease-specific disparities in terms of length and miscleavage distribution. Median peptide lengths recorded were within 10–12 aa for Ct+Tryp, 12–15 aa for Tryp, 11–14 aa for Ct, and 12–16 aa for AspN. Notably, peptides generated through *de novo* sequencing (Casanovo) were typically 1–4 aa shorter relative to those derived via PSM methods (MaxQuant & MSFragger). With respect to miscleavages, the proteases Ct+Tryp and Ct exhibited the highest proportion of peptides with minimum one miscleavage, yielding respective ranges of 65.09–75.21% and 48.75–60.90%. This was followed by Tryp (10.1%–36.22%) and AspN (14.10–29.06%). The substantial miscleavage prevalence among Ct+Tryp peptides was foreseeable, given the premature termination of the Ct digestion process, designed to prevent complete digestion thereby creating a different peptide distribution as compared to Ct peptides. On the other hand, AspN peptides displayed lower instances of miscleavage, deviating from the expectations based on existing literature ^74^. All preprocessed peptides were shown in Figure 2.

**Supplementary Figure 16:**
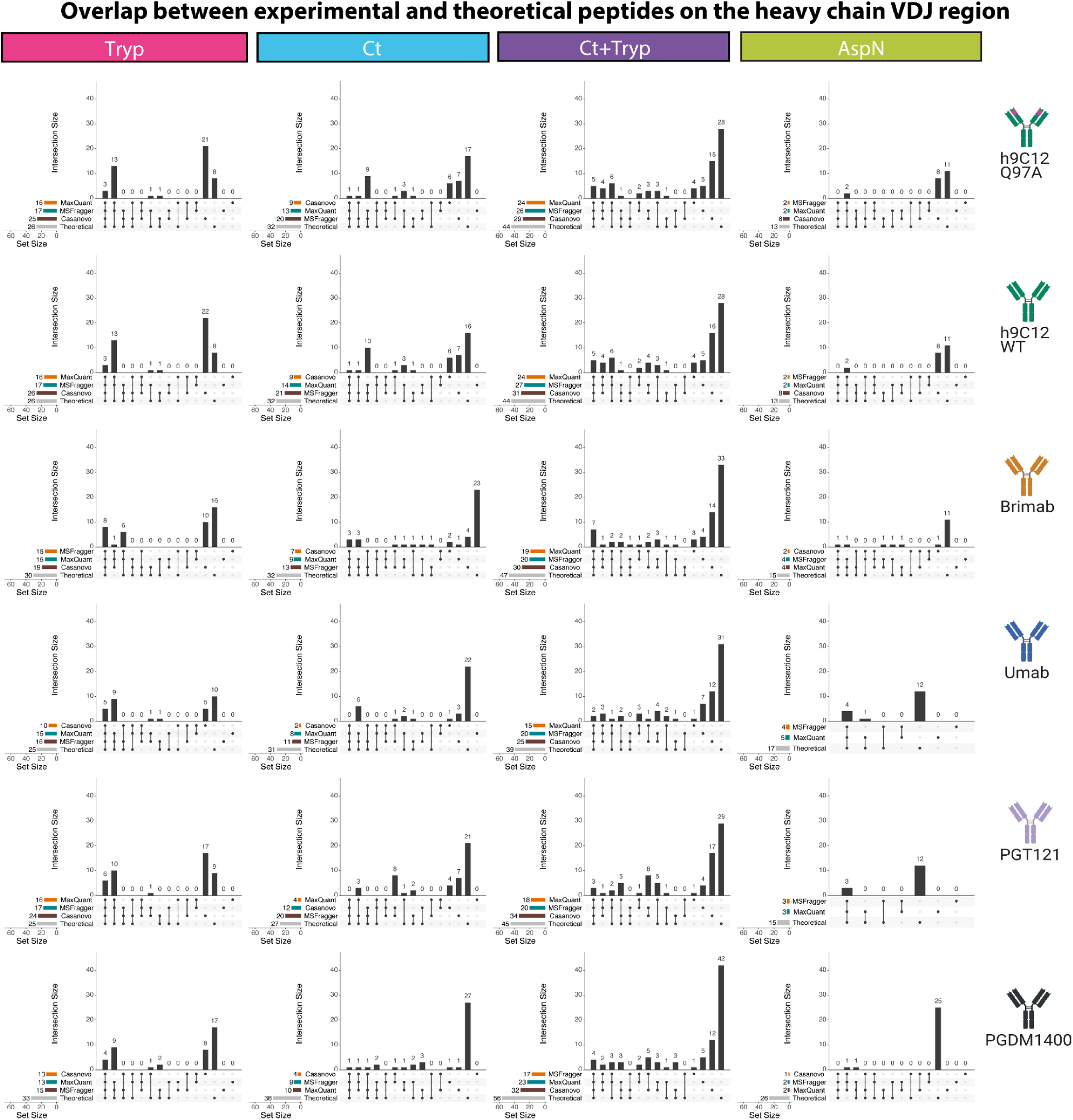
Overlap between peptide sequences from experimental data and from in silico cleavage of mAb sequences on the HC VDJ region. The variable region sequences of the six mAbs were cleaved in silico according to the stated specificity of the protease in ExPASy120 (Tryp, Ct, Ct+Tryp, AspN), with the options of having 0, 1, or 2 miscleavages allowed (denoted as theoretical) and compared to the peptide output from mass spectrometry experiments from all replicates in three proteomic tools (MQ, MSF, and Casanovo). The vertical bars show the number of peptides in exclusive intersection between sets, while the horizontal bars show the number of unique peptides in each set. All preprocessed experimental peptides were shown in Figure 2, Supplementary Figure 20.

**Supplementary Figure 17:**
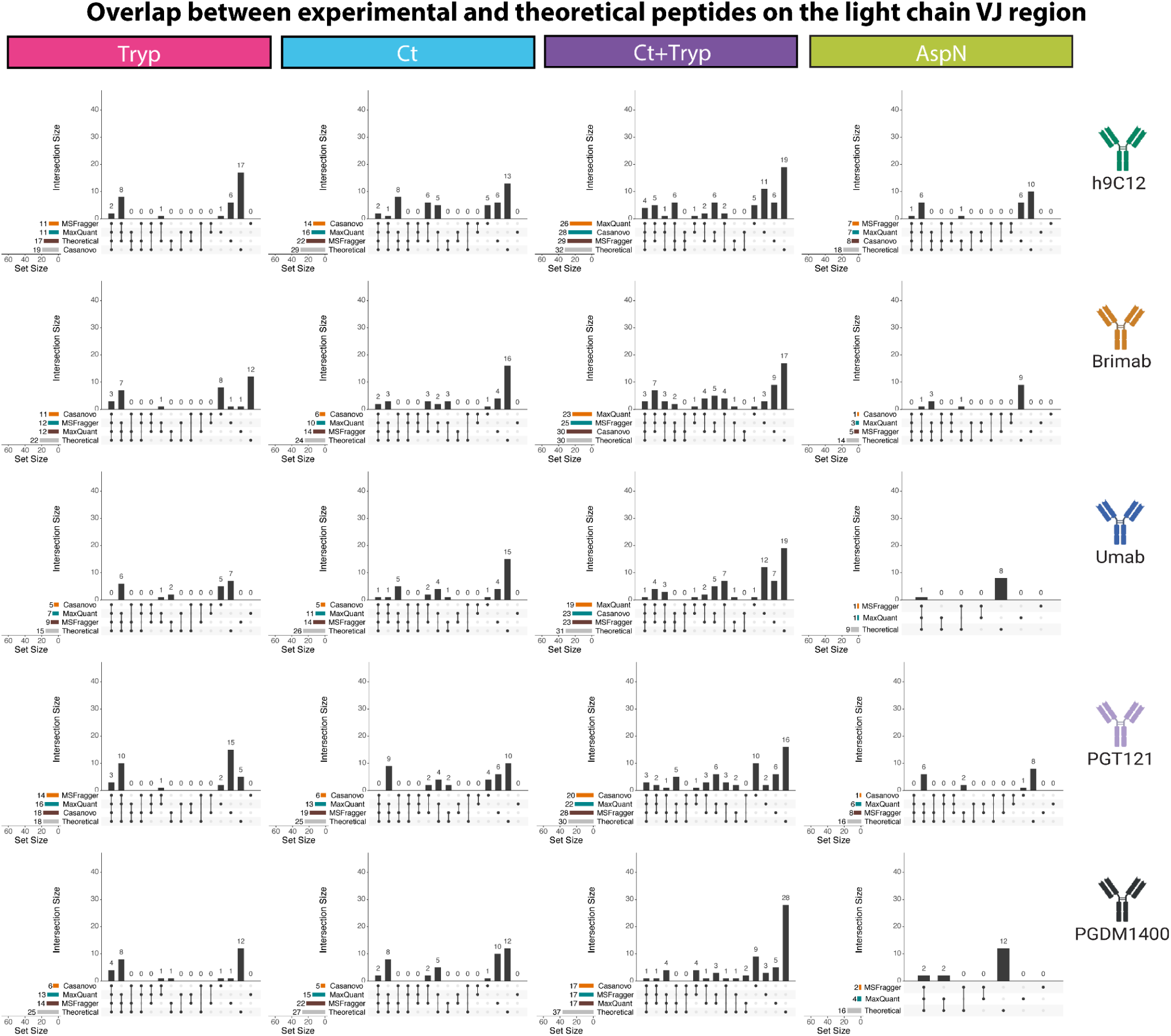
Overlap between peptide sequences from experimental data and from in silico cleavage of mAb sequences on the LC VJ region. The variable region sequences of the six mAbs were cleaved in silico according to the stated specificity of the protease in ExPASy120 (Tryp, Ct, Ct+Tryp, AspN), with the options of having 0, 1, or 2 miscleavages allowed (denoted as theoretical) and compared to the peptide output from mass spectrometry experiments from all replicates in three proteomic tools (MQ, MSF, and Casanovo). The vertical bars show the number of peptides in exclusive intersection between sets, while the horizontal bars show the number of unique peptides in each set. All preprocessed experimental peptides were shown in Figure 2, Supplementary Figure 20.

**Supplementary Figure 18:**
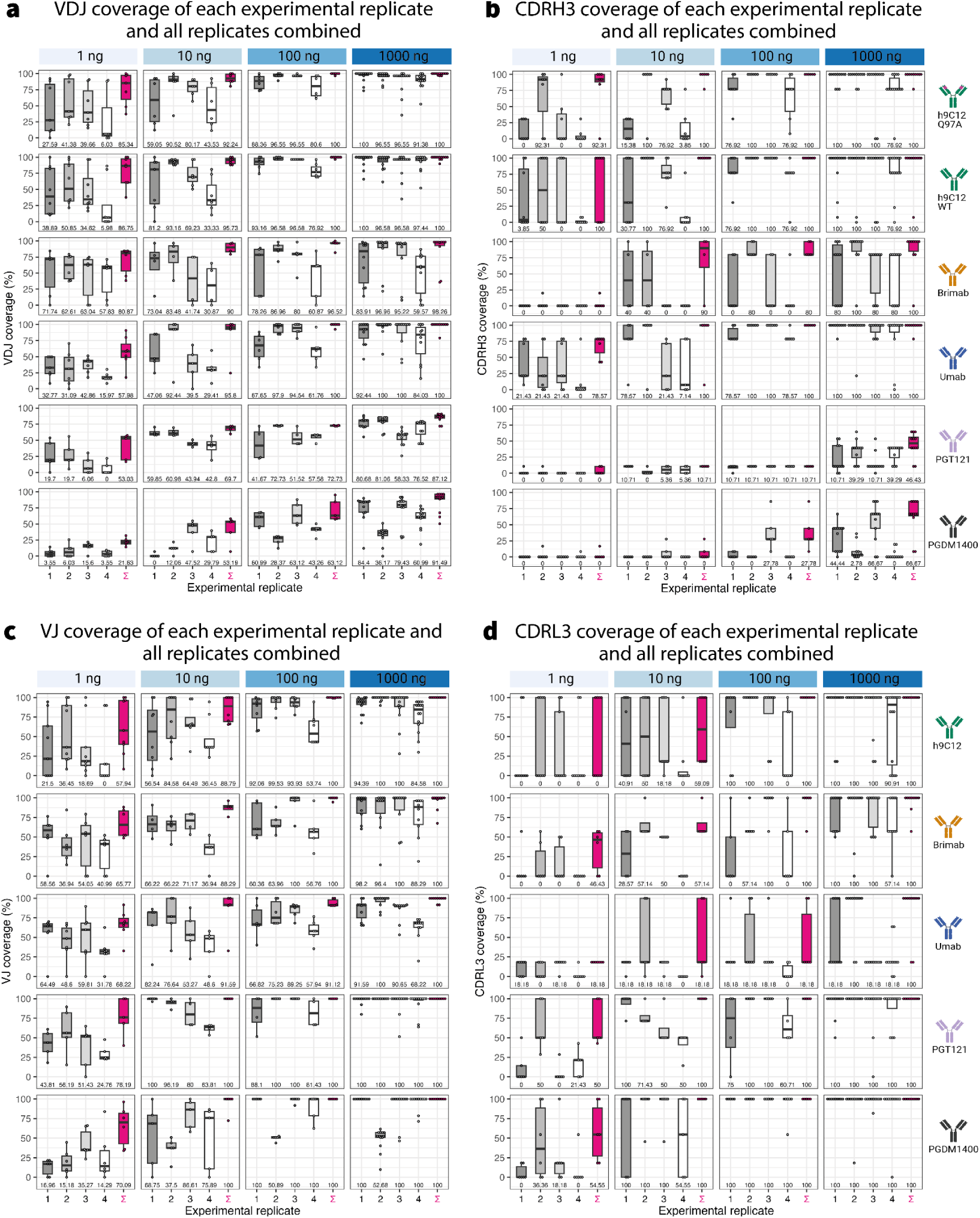
Coverage for each mAbs in the V(D)J and CDRH/L3 region by mAb input amount. Preprocessed peptides from all proteases were pooled, separated by mAb and input amounts from 1 ng to 1000 ng. Sequence coverage percentage for the (a) VDJ, (b) CDRH3, (c) VJ, and (d) CDRL3 for each experimental replicate individually and for all experimental replicates pooled together (∑). Cumulative coverage from merging experimental replicates for each mAbs in the VDJ and CDRH3 region by mAb input amount was shown in Figure 3.

**Supplementary Figure 19:**
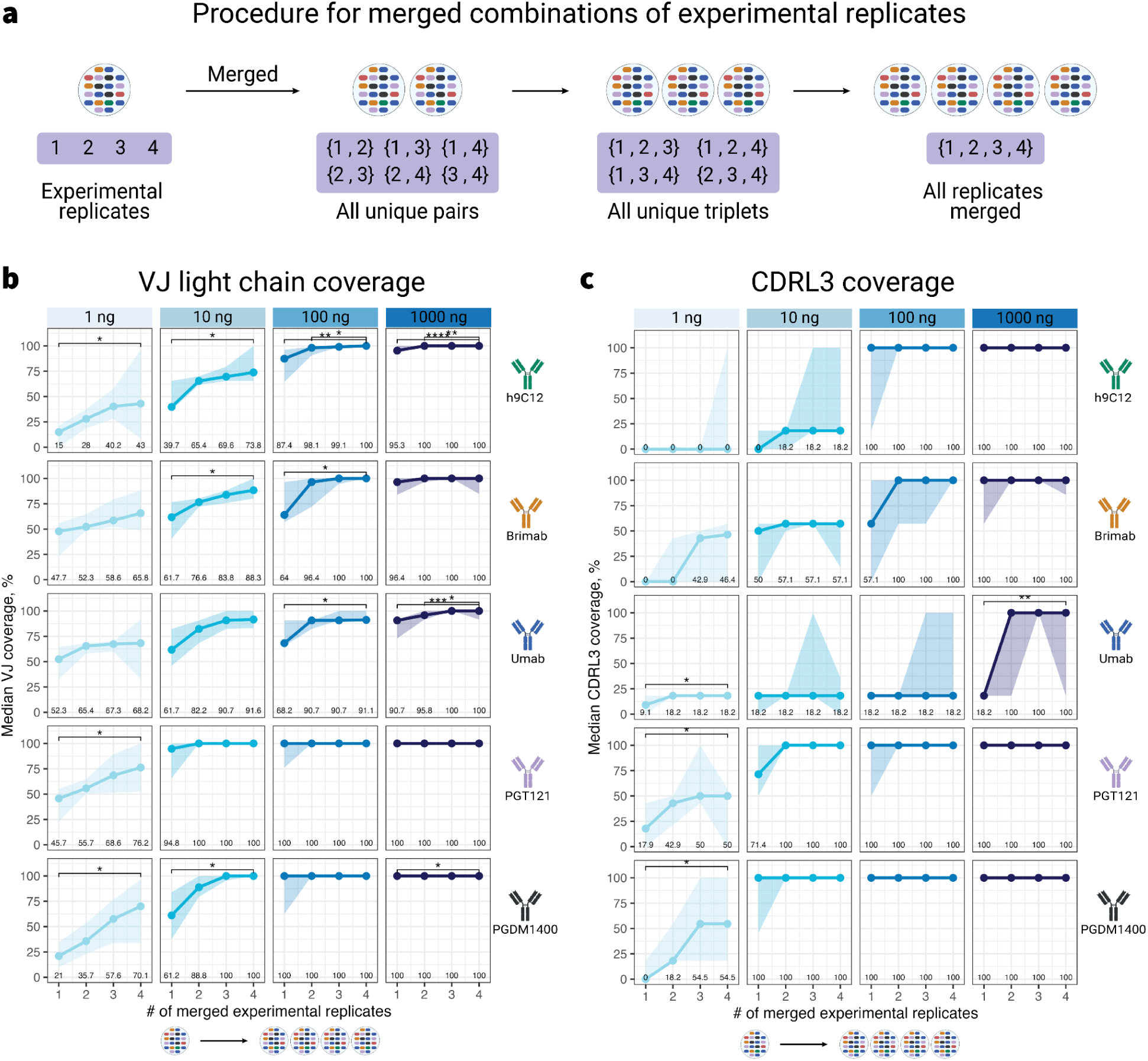
Experimental replicates are required for comprehensive V(D)J and CDR3 coverage. (a) Preprocessed peptides from all proteases, technical replicates, and proteomics analysis software programs were pooled, separated by mAb, and input amounts from 1 ng to 1000 ng. Sequence coverage for each sample in groups of cumulative merged experimental replicates from 1 (individual replicates), 2 (all unique pairs of replicates merged), 3 (all unique triplets of replicates merged), to 4 (all replicates merged) was shown for the (b) VJ sequence, (c) CDRL3 sequence on the LC. Values underneath each boxplot represent the median coverage in each group. Global differences between the sequence coverage values were determined using the Kruskal–Wallis (KW) test (adjusted for multiple testing by Benjamini–Hochberg method), and pairwise differences were determined by the Wilcoxon Rank Sum test, with p-values adjusted for multiple testing by Benjamini–Hochberg correction. All adjusted p-values lower than 0.05 are displayed above brackets as symbols: *(p.adj < 0.05), **(p.adj < 0.01), ***(p.adj < 0.001). Coverage of the VDJ and CDRH3 regions with similar groupings was shown in Figure 3. The number of unique peptide sequences for each experimental replicates and their pairwise overlap of peptide sets were shown in Supplementary Figure 20.

**Supplementary Figure 20:**
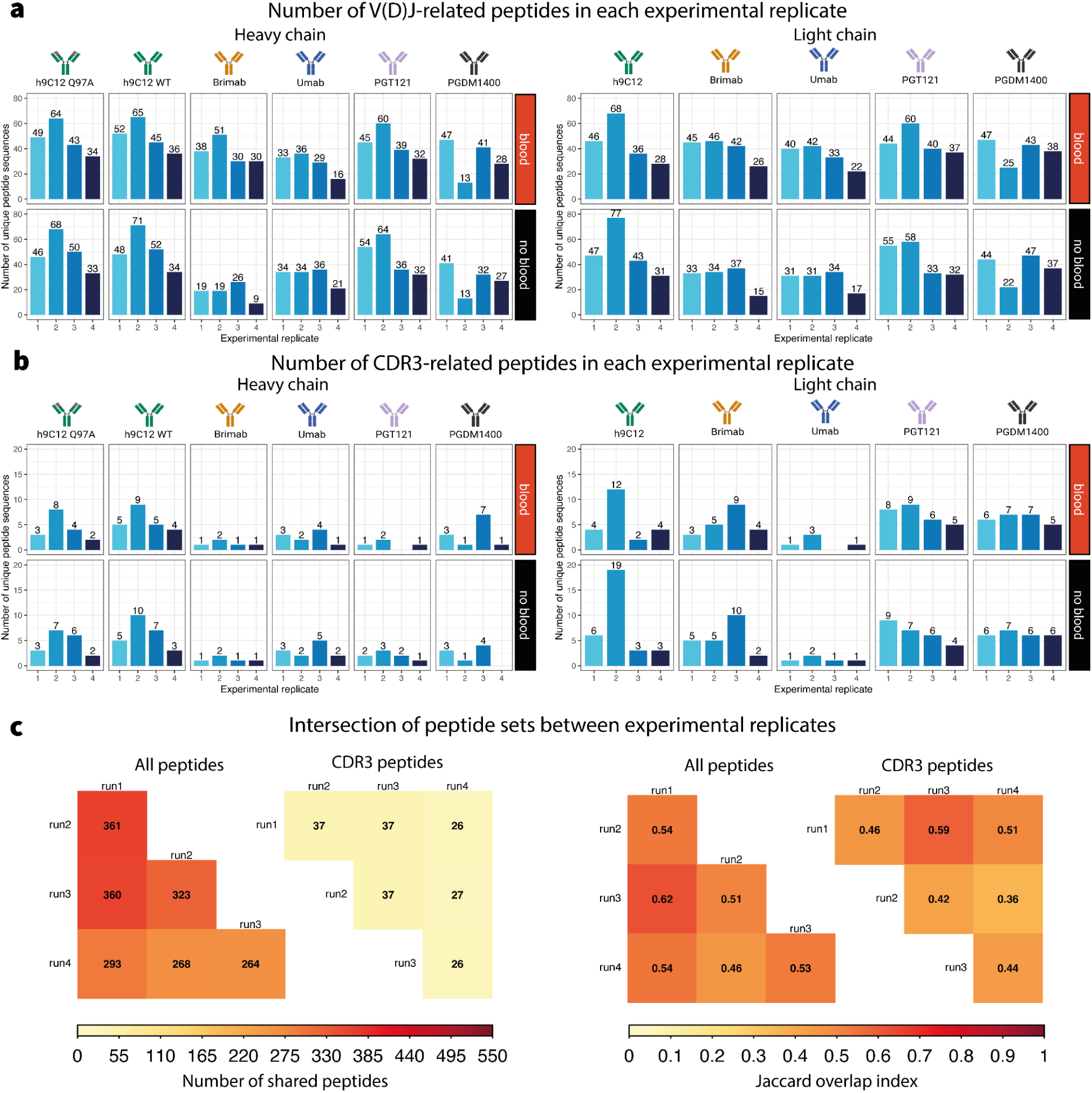
Number of unique peptide sequences by experimental replicates. Preprocessed peptides from all samples and technical replicates were pooled and grouped by experimental replicates. (a) Number of unique peptide sequences aligning to the reference mAbs’ VDJ region in samples with blood IgG (blood) and without blood IgG (no blood). (b) Number of unique peptide sequences aligning to the reference mAbs’ CDR3 region in samples with blood IgG (blood) and without blood IgG (no blood). (c) Overlap of peptide sets between experimental replicates in terms of number of unique peptide sequences shared (left panel) and Jaccard overlap index (right panel), with values ranging from 0 (mutually exclusive sets) to 1 (identical sets) calculated using the following formula: 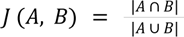, where A and B represent the peptide sets of two experimental replicates. The levels of sequence coverage provided by these peptides by individual experimental replicates are shown in Supplementary Figure 18 and replicates merged together are shown in Figure 3.

**Supplementary Figure 21:**
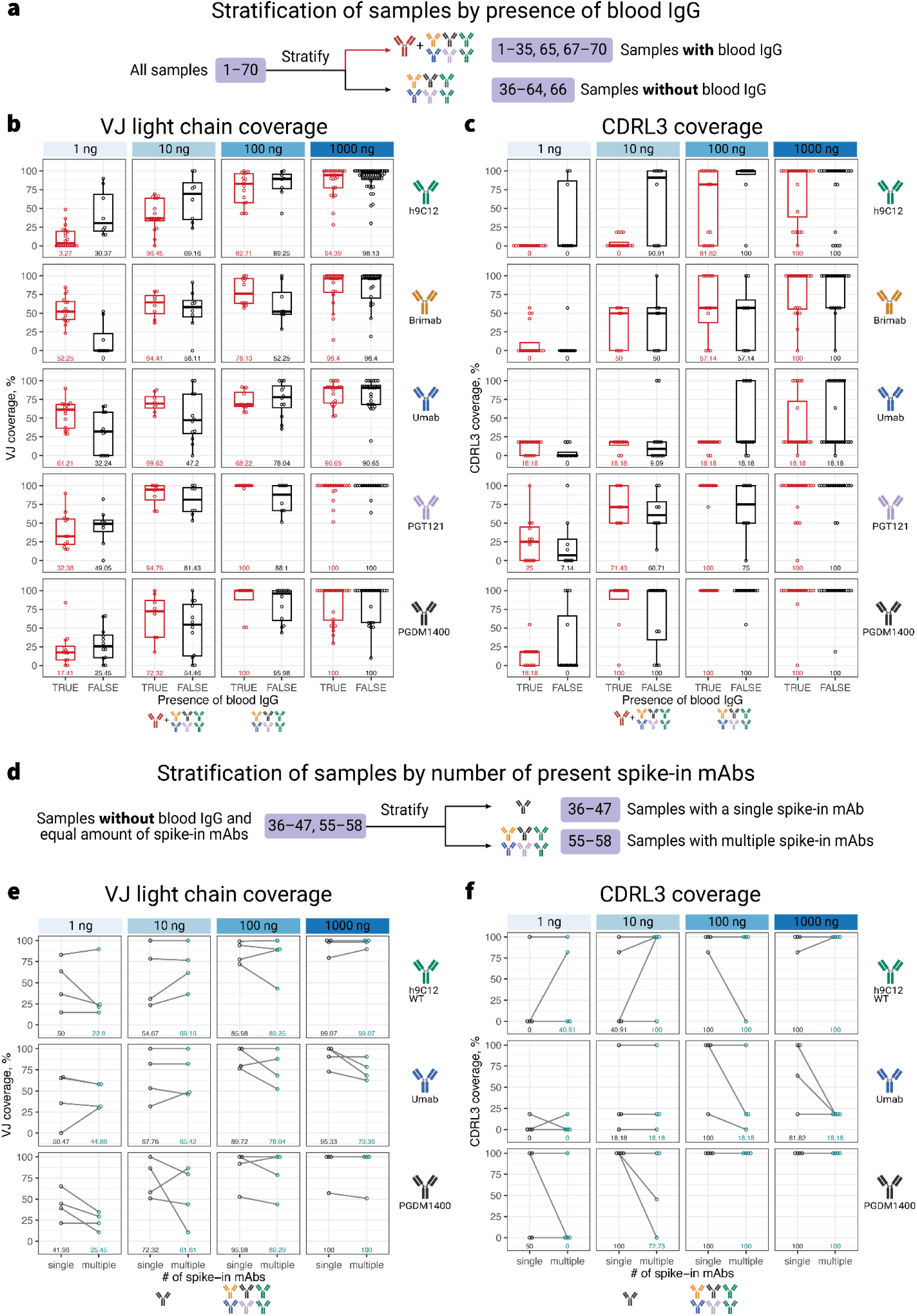
Analysis of LC sequence coverage with respect to presence of blood IgG and antibody diversity. **(a)** We aggregated true positive preprocessed peptides originating from all proteolytic treatments, technical replicates, proteomics software programs, and categorized the Ab-seq samples according to the diversity of antibodies, delineating by monoclonal antibody presence and ranging input levels from 1 ng to 1000 ng. Samples were further stratified by the absence (sample 36–64, 66) or presence (sample 1–35, 65, 67–70) of blood IgG background, and subsequent coverage metrics were computed for **(b)** the VJ region, **(c)** the CDRL3 region on the light chain. **(d)** Samples without blood IgG were further subdivided into single mAb (sample 36–47) and multiple mAbs samples (sample 55–58), with samples from the same experimental replicate connected with lines. Coverage metrics were computed for **(e)** the VJ region, **(f)** the CDRL3 region on the HC. Displayed beneath each boxplot is the group’s median coverage value. Pairwise differences were determined by the Mood’s median test, with p-values adjusted for multiple testing by Benjamini–Hochberg correction. All adjusted p-values were not statistically significant (p.adj > 0.05) and thus were not displayed in the figure. Comparative analysis of sequence coverage on the HC is shown in Figure 4.

**Supplementary Figure 22:**
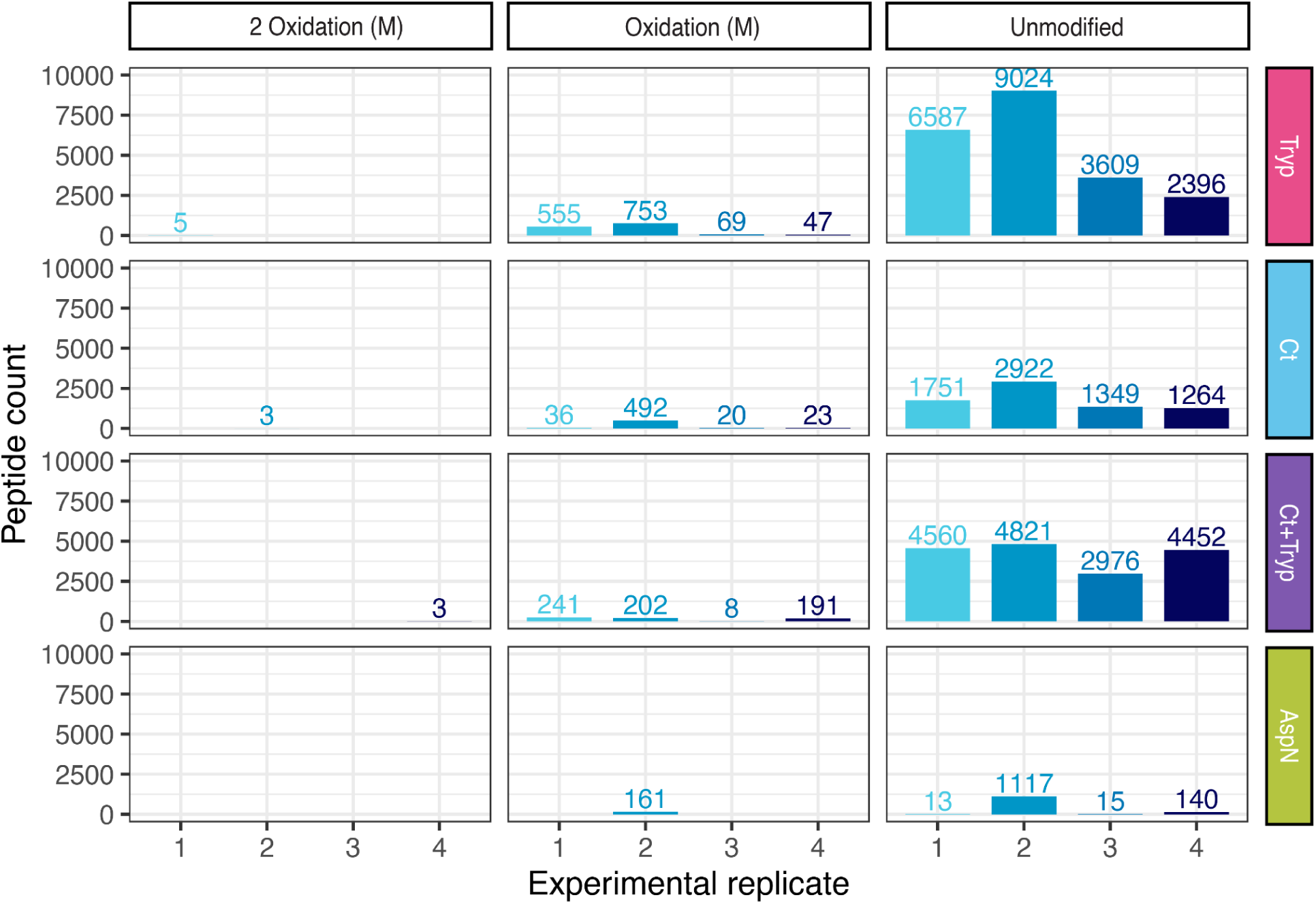
Distribution of peptide modifications with regard to experimental replicate and protease used. The number of unique peptides, considering their modifications, were calculated for each experimental replicate and protease treatment as reported by MaxQuant. Modification information was extracted from the “Sequence” and “Modification” columns in the evidence.txt file. The data includes only two modifications: 2 Oxidation (M) and Oxidation (M), with the majority of peptides being unmodified. Only unmodified peptides were utilized in the main figures (Figure 2, Figure 3, Figure 4, Figure 5, Figure 6).

**Supplementary Figure 23:**
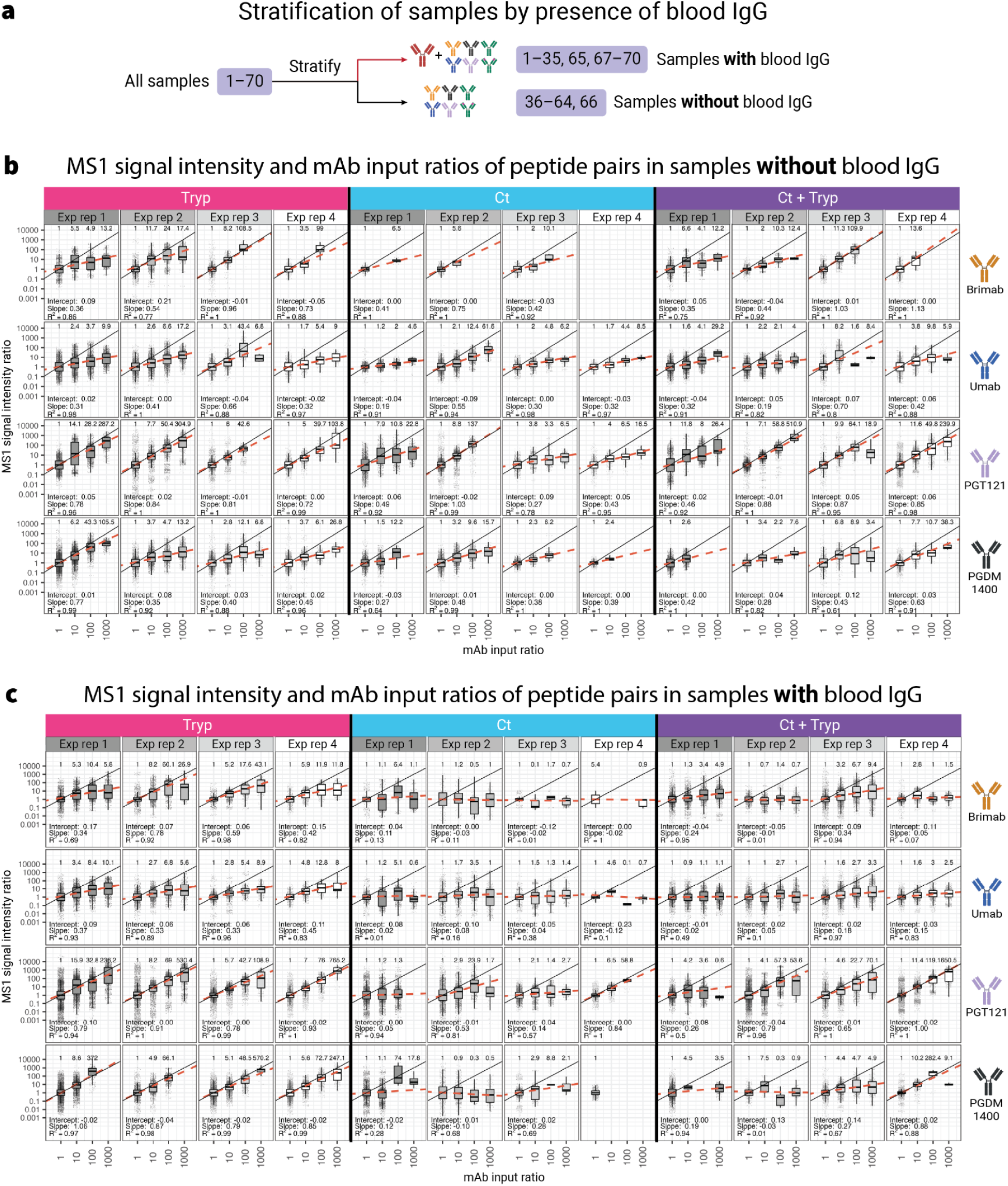
MS1 signal intensity ratio is a biased estimator of relative mAb input amount, and is biased by presence of blood IgG background, mAb sequence, and proteases utilized in digestion. **(a)** We stratified 70 samples based on the presence (sample 1–35, 65, 67–70) or absence (sample 36–64, 66) of blood IgG background. True positive preprocessed peptides from all protease treatments, technical replicates, and proteomics software programs (MQ and MSF only, as Casanovo does not report MS1 signal intensity for each peptide) were pooled together. Pairs of LC peptides aligning to the same mAb from the same experimental replicate (here denoted as exp rep for brevity) with identical amino acid sequences were formed and grouped according to the ratio of mAb input (1:1 to 1000:1, denoted as 1 to 1000 on the axis labels), and their reported MS1 signal intensity ratio was calculated (see Methods). h9C12 WT and h9C12 Q97A were excluded from this analysis because their nearly identical sequences would skew the MS1 signal intensity ratio on the peptide level. Peptides derived from ApsN were also excluded from this analysis due to a lack of data points. **(b)** MS1 signal intensity ratios in relation to mAb input ratios in peptide pairs from samples without blood IgG. **(c)** MS1 signal intensity ratios in relation to mAb input ratios in peptide pairs from samples with blood IgG. The values of MS1 signal intensity ratios and mAb input ratios were log10-transformed, and the median MS1 signal intensity ratios of each mAb-experimental replicate group were used for linear regression, weighted by the number of data points in each mAb input ratio. The dashed bright orange line represents the linear model that predicts the MS1 log10-signal intensity ratios from the log10-mAb input ratio, while the black solid represents the expected linear relationship of the two variables in log-log scale (slope 1, intercept 0). The median MS1 signal intensity ratio for each mAb input ratio is shown above each boxplot, while the intercept, slope, and the goodness-of-fit of the linear model – denoted by the coefficient of determination (R^2^), are shown below each boxplot. MS1 signal intensity ratios by mAb input for peptides on the HC is shown in Figure 5.

**Supplementary Figure 24:**
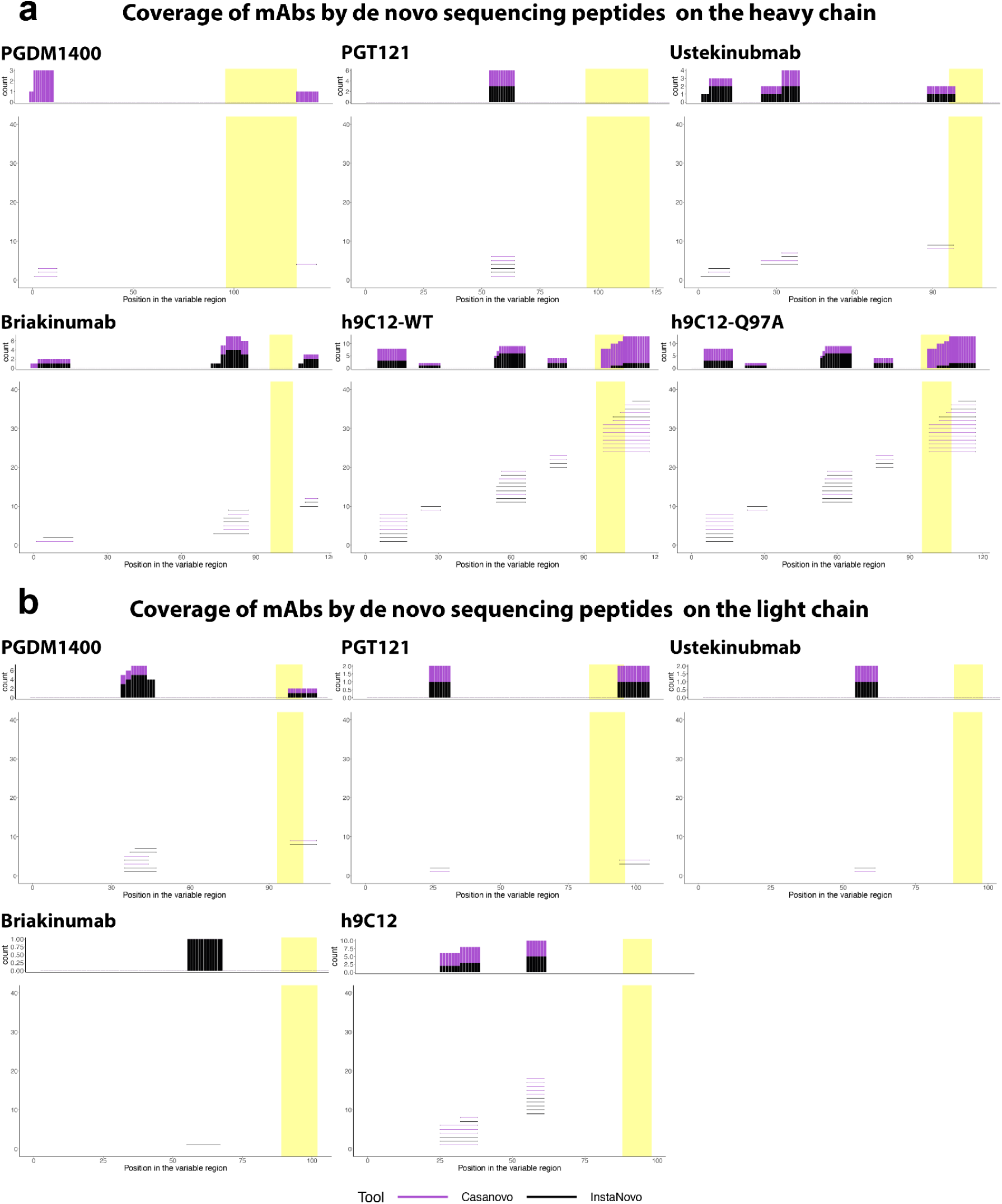
Coverage comparison between Casanovo and InstaNovo. A representative subset of the main dataset (described in Figure 2a): 6 mAbs samples at 1 ng + serum IgG background, 3 mAbs samples at 1000 ng without background IgG, and 1 combined all 6 mAbs at 1000 ng + background IgG, was utilized to compare the performance of Casanovo and InstaNovo regarding the coverage of the V region of the six mAbs on the (a) HC and (b) LC. De novo inferred peptides (as lines colored purple for Casanovo and black for InstaNovo) were then aligned to the V(D)J region of the mAbs’ sequences (CDR3 region highlighted in yellow) to visualize the coverage of MS peptides. Identical peptide sequences from different samples were not collapsed into a single sequence in order to visualize the ability of each tool to recover peptides from different input amounts. In brief, both Casanovo and InstaNovo managed to infer similar peptides mapped to the mAbs V(D)J region, and resulted in similar V(D)J sequence coverage.

**Supplementary Figure 25:**
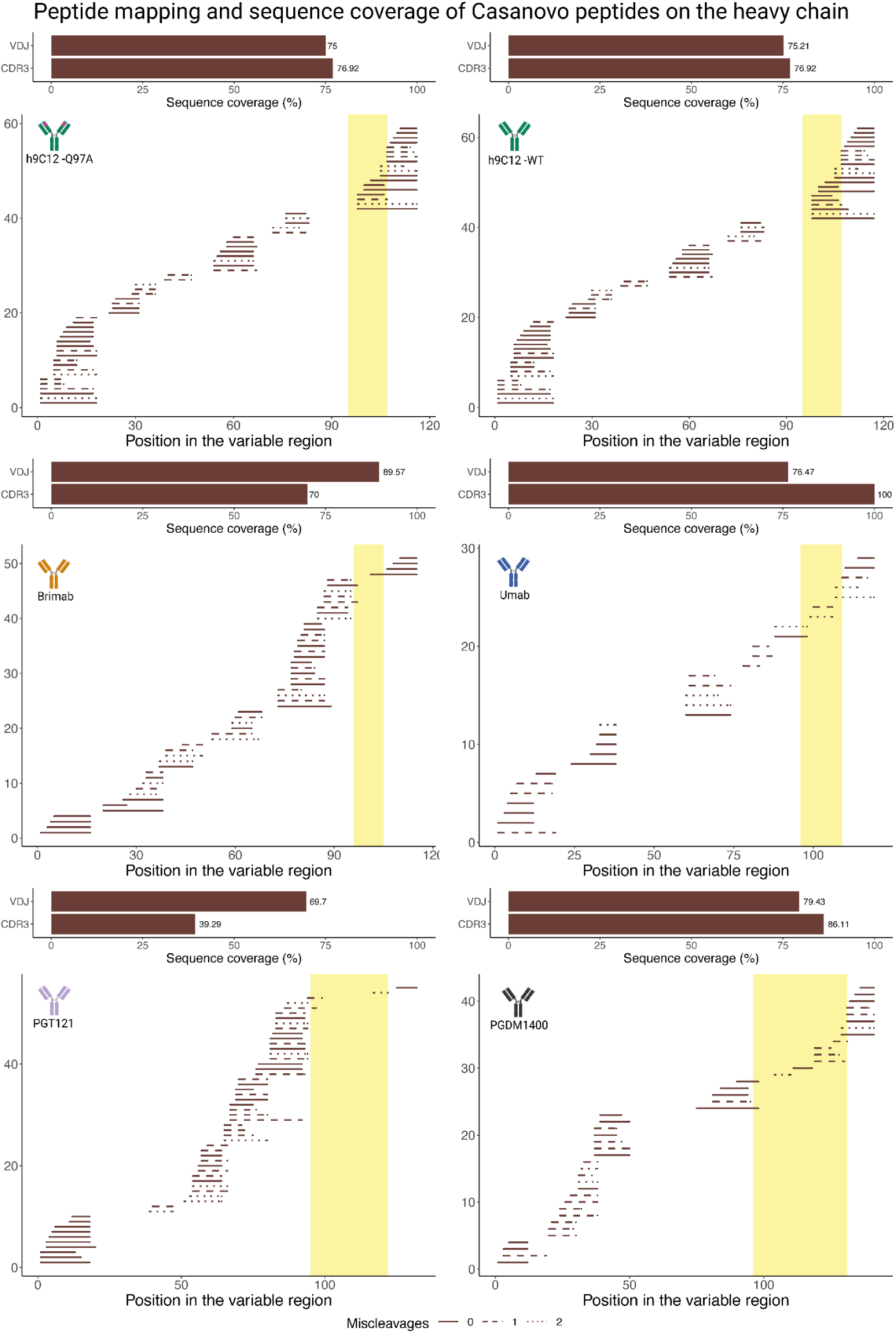
Position of Casanovo peptides on the reference sequence and sequence coverage of the mAbs on the HC. Peptides were pooled from all experimental and technical replicates, mAb inputs, and protease digestions. The yellow region represents the CDRH3, and line type represents the number of missed cleavages. Relates to Figure 6.

**Supplementary Figure 26:**
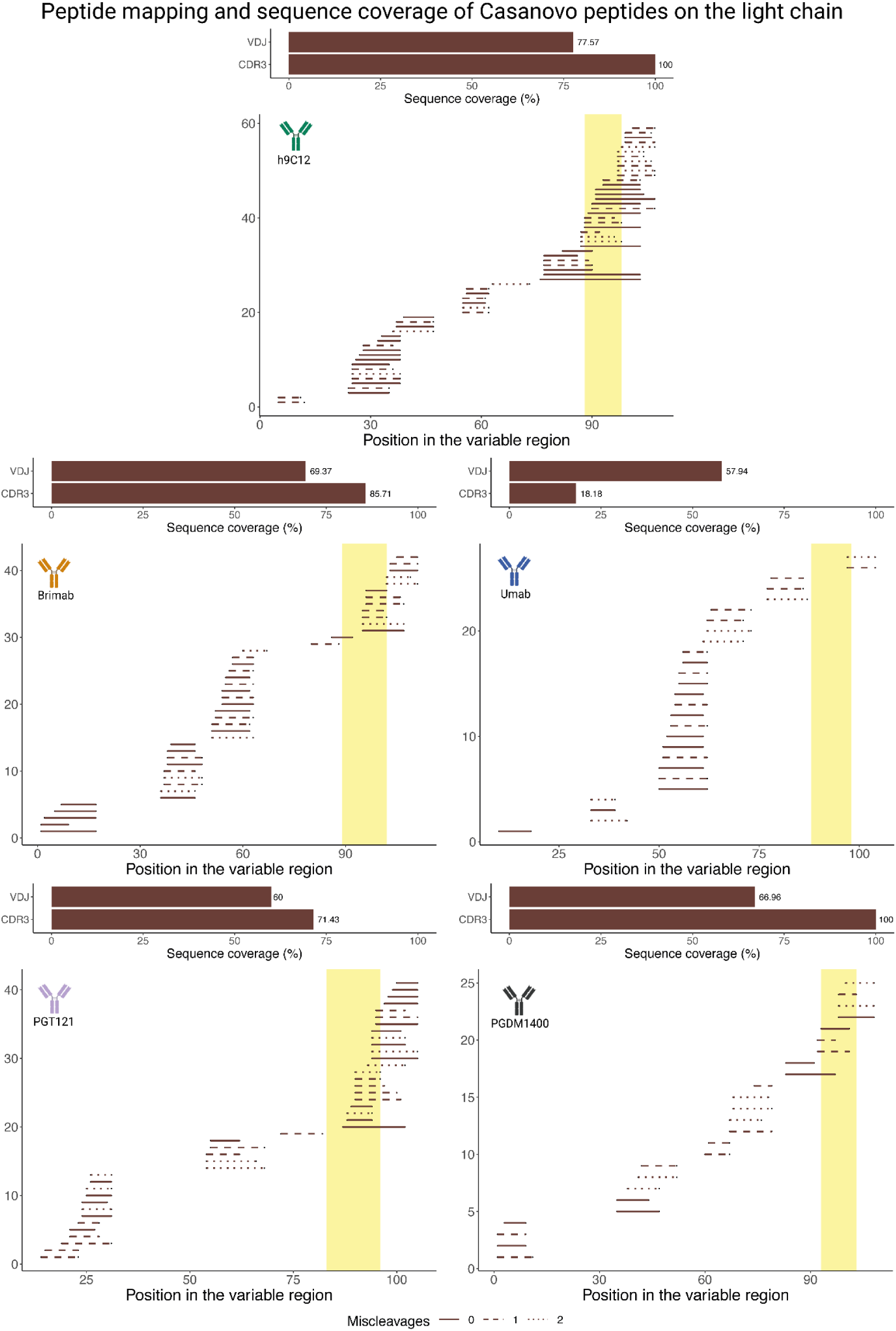
Position of Casanovo peptides on the reference sequence and sequence coverage of the mAbs on the LC. The yellow region represents the CDRL3, and line type represents the number of missed cleavages. Relates to Figure 6.

**Supplementary Figure 27:**
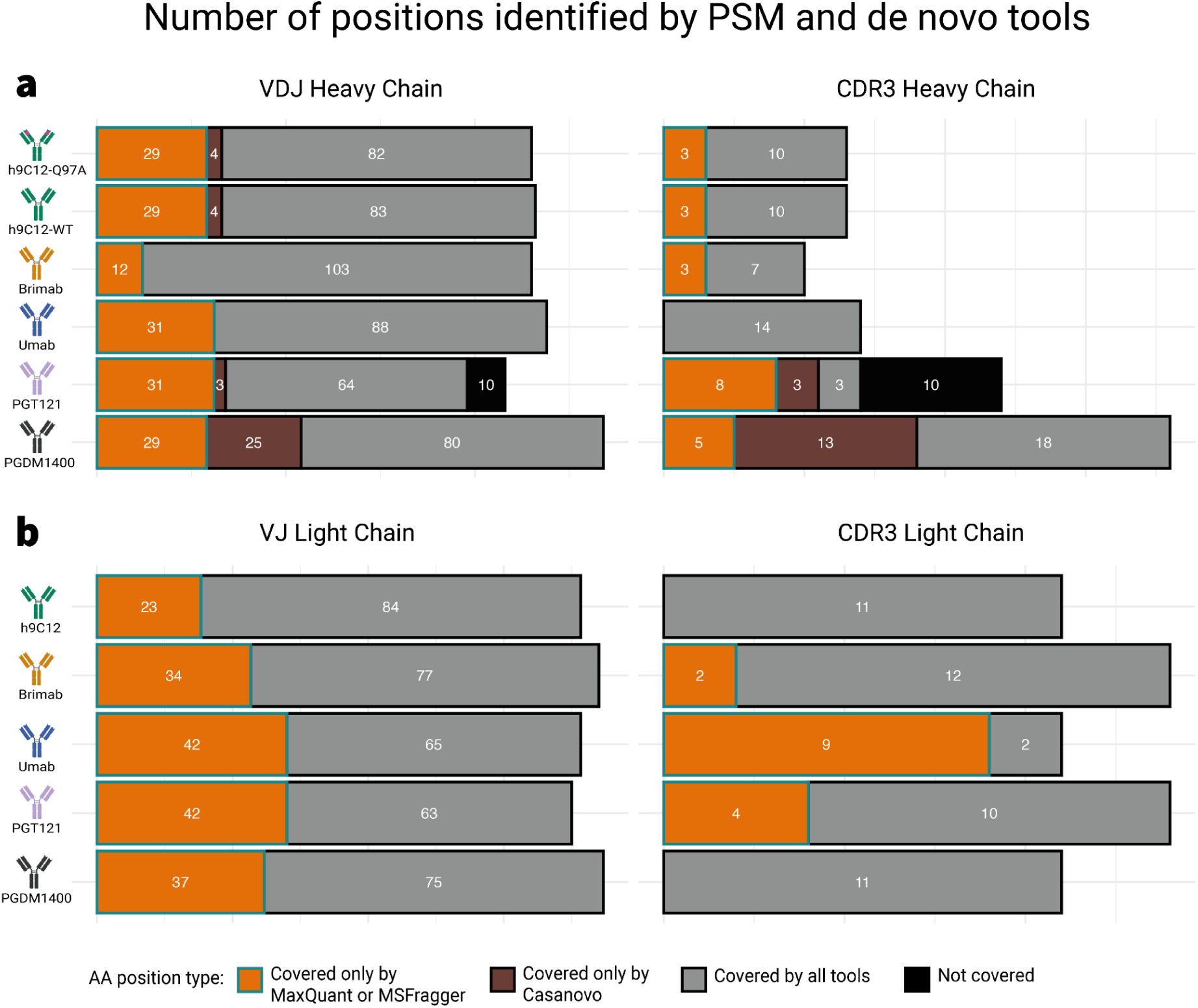
Number of positions in the V(D)J and CDR3 sequence covered by peptides from PSM tools (MaxQuant & MSFragger) and Casanovo. Bars indicate the number of residues covered only by MQ or MSFragger (orange with green outline), only by Casanovo (brown), covered by all tools (gray), or not covered by any tools (black) on the **(a)** HC and **(b)** LC. In contrast to the HC sequence, Casanovo-derived peptides did not cover any position that was not already covered by PSM tools (MaxQuant & MSFragger). Sequence coverage from each tool as percentage was shown in Figure 6c.

**Supplementary Figure 28:**
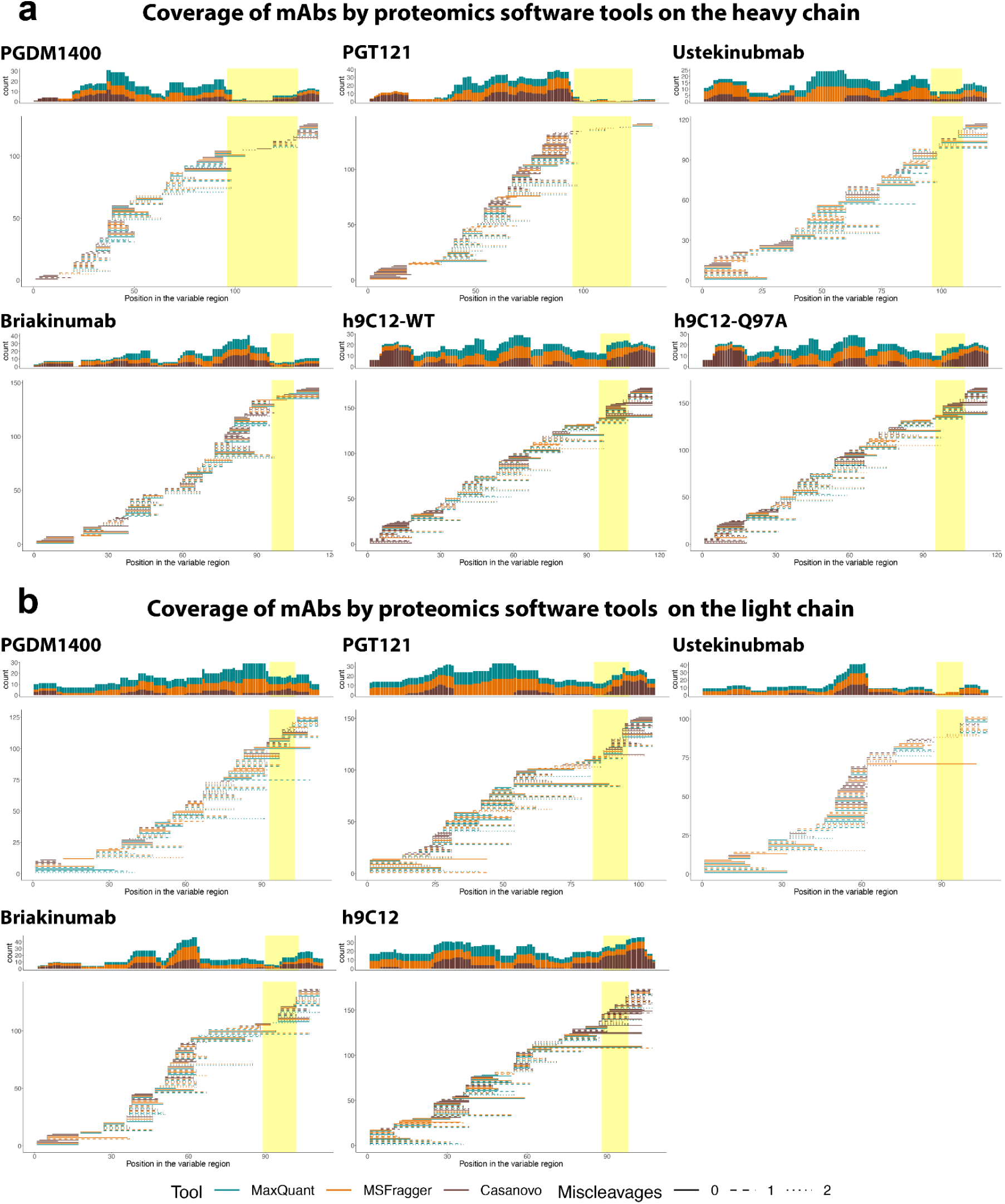
Experimental peptide coverage of mAbs (V(D)J and CDR3) on the HC and LC by proteomics software tools. Peptides pooled from all samples, technical replicates, experimental replicates, proteases, and proteomics software tools (as lines colored by software used and patterned by number of miscleavages) were aligned to the V(D)J region of the mAbs’ sequences on the **(a)** HC and **(b)** LC (CDR3 region highlighted in yellow) to visualize the coverage of MS peptides in regard to the whole V(D)J region. Histograms above each coverage plot represent the absolute number of peptides covering each amino acid position in the respective V(D)J region. All preprocessed experimental peptides were shown in Figure 2b as coverage percentage.

**Supplementary Figure 29:**
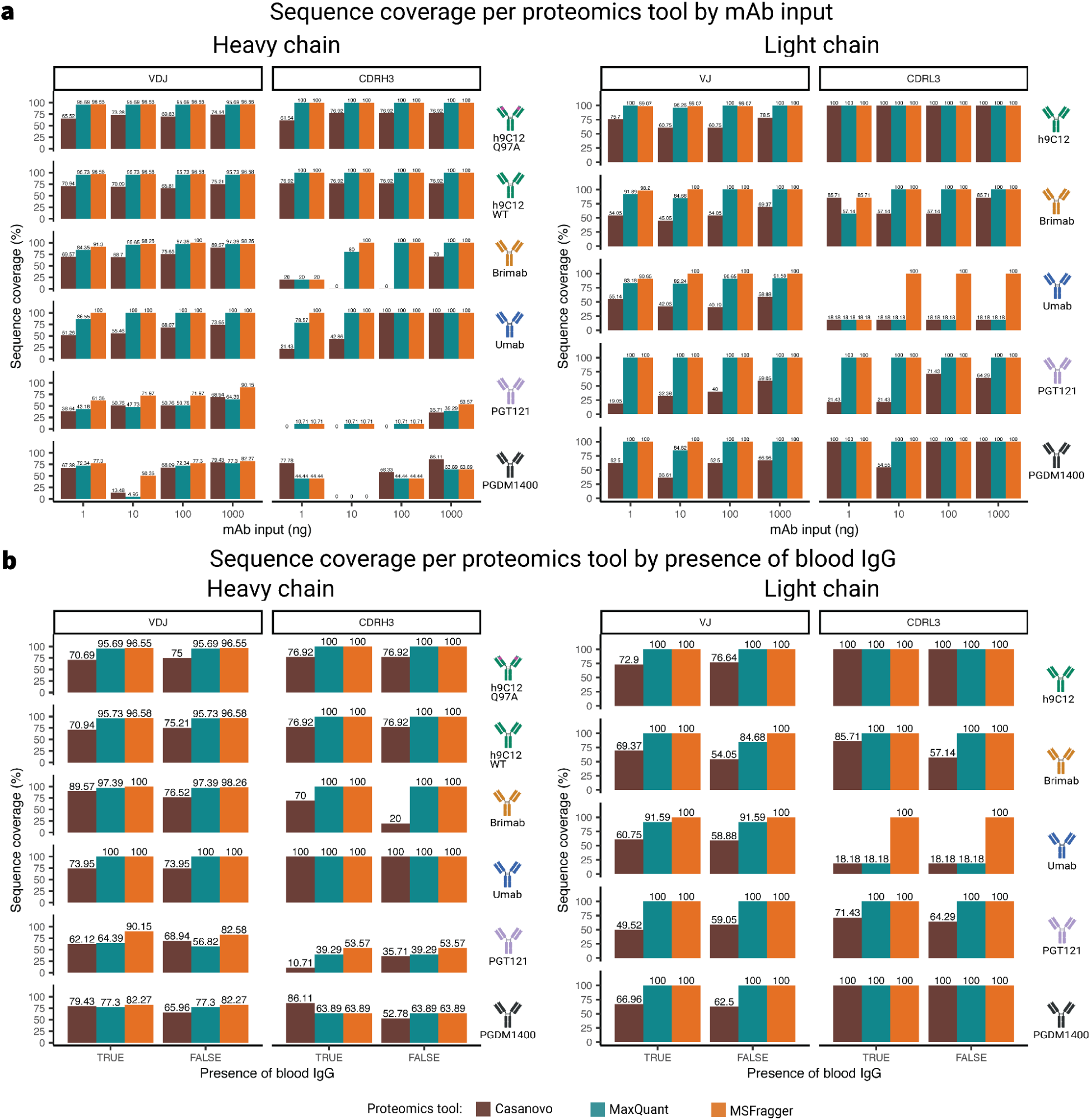
Sequence coverage of peptides per proteomics tools by mAb input levels and presence of blood IgG. (a) Sequence coverage of Casanovo peptides for each mAb at inputs ranging from 1 ng to 1000 ng on the HC and LC. (b) Sequence coverage of Casanovo peptides for each mAb in samples with or without the presence of blood IgG background on the HC and LC. Relates to Figure 6 and Supplementary Figure 25, Supplementary Figure 26.

**Supplementary Figure 30:**
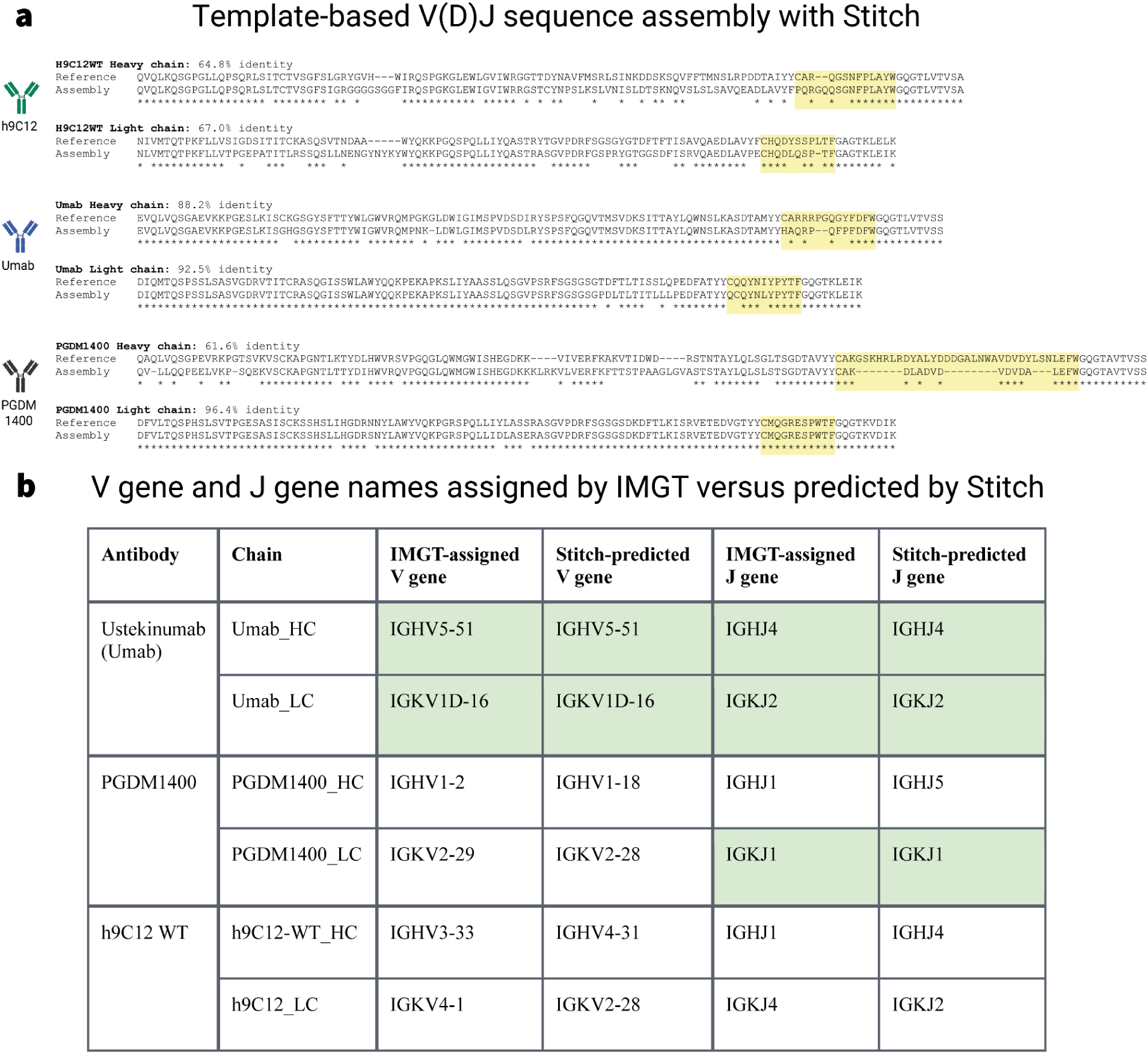
Assembly of de novo sequencing peptides by Stitch. Samples with only one mAb present and without blood IgG background from all replicates (samples 36–47) and sequenced de novo by Casanovo having prediction score of ≥0.8 were selected for V(D)J sequence assembly. The Stitch software22 was used to align de novo peptides to germline V, (D), and J gene segments from the IMGT reference, assigned a score for each gene segment, and recombined the most probable V(D)J region sequence (here, the term “recombined” is used in the sense of the Stitch manual). (a) The resulting consensus sequences were aligned to the reference mAb sequence, and the sequence identity percentage was calculated. The yellow region denotes the CDR3 region on the HC and LC. (b) V and J gene names for the mAbs assigned by IMGT were compared to those predicted by Stitch. The green background indicates instances when the gene name predicted by Stitch overlapped with the gene name assigned by IMGT. Relates to Figure 6.

## Supplementary Files

Supplementary File 1: MaxQuant parameter file example for the tryptic samples from run1

Supplementary File 2: MSFragger parameter file example for the tryptic samples from run1

Supplementary File 3: Casanovo config file

Supplementary File 4: Stitch example config file for h9C12 WT

All Supplementary files are accessible on GitHub: https://github.com/csi-greifflab/Systematic-benchmarking-of-mass-spectrometry-based-antibody-sequencing-reveals-methodological-biases

