## Supplementary File 3 for "Systematic benchmarking of mass spectrometry-based antibody sequencing reveals methodological biases": Supplementary_file_3.yaml

```
###
# Casanovo configuration.
# Blank entries are interpreted as "None"
###

# Random seed to ensure reproducible results.
random_seed: 454

# Spectrum processing options.
n_peaks: 150
min_mz: 50.0
max_mz: 2500.0
min_intensity: 0.01
remove_precursor_tol: 2.0  # Da
max_charge: 10
precursor_mass_tol: 50  # ppm
isotope_error_range: [0, 1]

# Model architecture options.
dim_model: 512
n_head: 8
dim_feedforward: 1024
n_layers: 9
dropout: 0.0
dim_intensity:
custom_encoder:
max_length: 100
residues:
  "G": 57.021464
  "A": 71.037114
  "S": 87.032028
  "P": 97.052764
  "V": 99.068414
  "T": 101.047670
  "C+57.021": 160.030649 # 103.009185 + 57.021464
  "L": 113.084064
  "I": 113.084064
  "N": 114.042927
  "D": 115.026943
  "Q": 128.058578
  "K": 128.094963
  "E": 129.042593
  "M": 131.040485
  "H": 137.058912
  "F": 147.068414
  "R": 156.101111
  "Y": 163.063329
  "W": 186.079313
  # Amino acid modifications.
  "M+15.995": 147.035400    # Met oxidation:   131.040485 + 15.994915
  "N+0.984": 115.026943     # Asn deamidation: 114.042927 +  0.984016
  "Q+0.984": 129.042594     # Gln deamidation: 128.058578 +  0.984016
  # N-terminal modifications.
  "+42.011": 42.010565      # Acetylation
  "+43.006": 43.005814      # Carbamylation
  "-17.027": -17.026549     # NH3 loss
  "+43.006-17.027": 25.980265
n_log: 1
tb_summarywriter:
warmup_iters: 100_000
max_iters: 600_000
learning_rate: 5e-4
weight_decay: 1e-5

# Training/inference options.
train_batch_size: 32
predict_batch_size: 1000
n_beams: 5

logger:
max_epochs: 30
num_sanity_val_steps: 0

train_from_scratch: True

save_model: True
model_save_folder_path: ""
save_weights_only: True
every_n_train_steps: 50_000
```
